# Coordinated Brain-Network Dynamics of Maternal Adaptation

**DOI:** 10.64898/2026.09.17.751574

**Authors:** Sara B. Mitchell, Yilin Wang, Himali Gilliland, Zhihan Chen, Ian Hultman, Alli Jimenez, Allan Ngumo, Kiriana Williams, Sanvesh Srivastava, Kung-Sik Chan, Hanna Stevens, Rainbo Hultman

## Abstract

The maternal brain exhibits impressive plasticity, undergoing many anatomical and functional alterations over the transition to parenting^1–3^. Such alterations can reflect maternal adaptation to offspring needs^4–6^, aligning with the onset and maintenance of maternal care behaviors^7–9^. These brain-wide neural changes occur across stages, starting during pregnancy and continuing through the peripartal transition^10,11^. We therefore wanted to understand brain changes with pregnancy, parenting, and care behaviors over time. Many brain regions have been implicated in the coordination of maternal adaptation and corresponding care behaviors ^10,12,13^. We used computational modeling of multi-region neural oscillation data to identify distinct electrical networks during naturally-occurring care behaviors and across multiple maternal stages in outbred, highly maternal mice^14^,. This work demonstrates that multi-region oscillatory electrical dynamics at high spatiotemporal resolution reflect these functions. Distinct, discriminative networks predict maternal behavior and stage, showcasing network adaptation with parental transition. Maternal behavior and stage also had substantial effects on other distinct, multi-region oscillatory electrical networks relevant to stress susceptibility ^15^, further linking maternal adaptation to the neurophysiology of stress. While these stress networks were perturbed by maternal experience of early life stress, networks trained on maternal behavior and stage remained robust against such perturbation. This suggests a uniquely persistent strength of maternal electrical dynamics. Together, this work supports the role of oscillatory electrical dynamics as networks underlying fundamentally conserved behaviors and transitions, demonstrating wide relevance and significance to brain–behavior discovery.

## Introduction

Numerous regions across the brain have been implicated in different aspects of maternal behavior. Human neuroimaging studies show changes in structure, regional activity, and connectivity (i.e. fMRI) throughout the maternal transition, including altered cortical volume and increased functional network connectivity, such as between the amygdala and nucleus accumbens in association with maternal behavior^16^. In rodent studies, behaviors for which specific neural substrates have been identified include: the onset and regulation of pup-directed behavior^17,18^, the structuring of highly coordinated care behaviors (i.e., licking/grooming of pups, pup retrieval, nest building)^19–22^, reinforcement learning for repetitive stereotyped parental behavior^23^, individual maternal motivation^24^, pup defense behavior ^25^, and social aspects of pup interaction^12^. Some regions that have ties to these behavioral elements include anatomical correlates of the prefrontal cortex (PFC): the prelimbic (PrL) and infralimbic cortices (IL)^22,25^; the nucleus accumbens (NAc)^24^; multiple sub-regions of the amygdala (AMY): the medial (MeA)^12^, central (CeA)^26^, and basolateral (BLA) amygdala^27,28^; the ventral tegmental area (VTA)^20,23^; and the ventral hippocampus (VHipp)^19^. Collectively, these regions imply maternal network involvement. Therefore, we sought to identify the spatiotemporal relationships of oscillatory electrical dynamics between these regions over the maternal transition and during care behavior. We hypothesized that maternal stage and emergent maternal care behavior have unique network activity involving these regions. We further hypothesized that the strength of network activity has distinctive trajectories over this transitional period. Thus, we investigated the strength of these networks over time.

Throughout this investigation, we evaluated neural oscillations via local field potential (LFP) recordings. These recordings were captured using multi-site in vivo electrophysiology in adult female mice prior to conception, during pregnancy, and over several postpartum timepoints. We further time-locked these LFPs to behavioral measurements and used a machine learning approach to train several maternal-relevant electrical brain network signatures. While the plastic maternal brain undergoes extensive alterations as a means of adaptation, maladaptive brain disorders linked to stress and emotional dysregulation also involve altered connectivity across multiple brain regions^29,30^. Many of the same neural circuits engaged in maternal brain adaptation are likewise engaged in stress responsive anxiety and mood disorders^31^, strongly suggesting that stress vulnerability is relevant to the maternal transition. With this in mind, we applied our LFP measurements to previously established stress-relevant networks^15^, characterizing their dynamics across this transition.

We also evaluated the influence of prior stress on the maternal experience at both the behavioral and brain-wide network levels. It has been shown that adversity in early life has lasting impacts on the brain and can heighten negative reactions to future life stressors^29^, such as the maternal transition^32^. Therefore, an early life stress (ELS) paradigm was performed in a subset of the experimental animals to enhance neuropsychiatric vulnerability^33–35^ throughout pregnancy and postpartum. We assessed the impacts of ELS on novel networks in association with maternal stage and behavior as well as on distinct stress-relevant networks. While ELS did perturb some of the known stress networks, we observed multiple signatures of maternal-relevant network activity robust to stress. Overall, we identified multiple simultaneous network dynamics governing different aspects of the parental transition, a highly conserved mammalian experience that provides rich information for understanding brain plasticity.

## Methods

### Animal Care and Use

Animal experiments were performed in accordance with the University of Iowa Animal Use and Care Committee. CD1 mice (Charles River) were group housed by sex prior to breeding and maintained on a 12-hour light/dark cycle with ad libitum access to food and water. Studies were performed during the light cycle. Mice were single-housed following surgical implantation of recording electrodes (Fig. 2a). Cage changes were carried out weekly, apart from during the postpartum period, when they were left undisturbed until being changed at 10 days postpartum^36^.

### Breeding, Rearing, and Early Life Stress (ELS)

Experimental animals were generated by breeding pairs of CD1 mice 9-12 weeks of age, a minimum of one generation removed from shipping stress. To generate control and early life stress (ELS) experimental females for electrode implantation, dams were singly housed during pregnancy and following parturition. All litters were culled to 8 pups on postnatal day one (P1), making efforts to evenly balance sex distribution. Standard cage conditions consisted of Beta Chip®, a non-nestable woodchip bedding, instead of corncob which may have varied hormonal impacts^37^. Nine grams of Enviro-dri® (Shepherd Specialty Papers, Richland, MI) supplemental nesting material was additionally provided. At P2, a randomly selected subset of litters was assigned to the ELS condition. The paradigm was primarily modeled after similar work in CD1s^36^, yielding emotionally-relevant molecular and behavioral impacts on exposed animals. To mimic conditions of neglect and limited resources, the ELS paradigm involved the combination of daily maternal separation, limited nesting materials, and early weaning^36,38–40^. ELS litters had 6g of supplemental nesting material removed from the cage and were separated from their dams for periods of 4 hours on days P2-P5 (increased to 8 hours on days P6-16). During daily separations, dams were placed in a separate cage (the same cage was used for all separation days), and pups were removed to an adjacent room to prevent dams from hearing pup vocalizations. Pups were placed on a heating pad to aid in regulating body temperature. ELS litters were early weaned at P17 and co-housed with same sex siblings. Water-softened chow and Napa Nectar (SE Lab Group Inc., Hickory, NC) were provided for the ELS animals to support hydration until P21, the control wean date, when they were group housed by sex in standard cage conditions. All experimental animals were ear-punched at P21 for individual identification purposes.

### Adolescent Behavioral Assays

Behavioral studies were performed across an age range equivalent to adolescence in mice^41^ to evaluate early impacts of ELS. Time of day was maintained for the completion of each behavioral task. Following a 30-minute minimum habituation to test room and dimmed white light conditions, mice performed the following tasks (one per day with immediate return to their homecage). Statistics: differences between groups (control vs. ELS) were assessed via student’s un-paired t-test (open field) or two-way repeated measures ANOVA (3-chamber social). Tasks were performed in order of increasing stress intensity, as listed below.

#### Open Field

At P32-34, experimental mice were placed in a rectangular plexiglass arena (40 × 39 × 30 cm) for 30 minutes. ANY-maze Software tracked and recorded all movements for the duration of the trial. Basal locomotor activity was assessed through total distance traveled.

#### 3-Chamber Social Interaction

At P34-36, experimental mice were evaluated for social behavior. This experiment required two consecutive trials per mouse, 5 minutes per trial. Mice were placed into a plexiglass arena (15.5□ x 23□ x 8.5□) separated into 3 chambers of equivalent size (i.e. left chamber, middle chamber and right chamber). For both trials, a cylindrical cage (4.2□ diameter x 8□ high) was placed in the left and right chamber. For the first trial, both cylindrical cages were empty. The experimental mouse was placed in the middle chamber and given freedom to explore any of the 3 chambers at will, being returned to its homecage after 5-minutes. The chamber was cleaned between trials (Super Sani Cloth®, PDI Inc., NJ). For the second trial, an unfamiliar, but sex and aged-matched conspecific CD1 mouse was placed in one of the cylindrical cages. The curved surface of the cylinder was barred to allow for subject interaction. The placement of the conspecific was counter-balanced among experimental animals to control for unforeseen spatial biases of the mice. The experimental mouse was then placed back into the middle chamber and was given freedom to explore all 3 chambers for an additional 5 minutes. For each trial, a video tracking system (EthoVision XT, Noldus, Leesburg, VA) was used to assess social behavior via the duration of time experimental mice spent in each chamber and in the direct interaction zones around the social target (proximal) or empty cylindrical cage (distal).

### Selection of Animals for Maternal Behavior and Neurophysiological Studies

At 6 weeks of age, one female from each control or ELS assigned litter that had undergone evaluation of adolescent behavior was selected at random for additional procedures. These procedures included surgical electrode implantation, neurophysiological data acquisition, breeding, and subsequent assessment of maternal experiences, including maternal care observations. Animal weights were accounted for to ensure that an unbiased range was selected during random sampling across litters.

Females used as non-implanted controls did experience breeding, but they were not subject to surgical implantation nor any behavioral observations or assays until their first observation timepoint postpartum (P3).

### Electrode Implantation Surgery

Mice were anesthetized with 1-5% isoflurane during the surgical implantation of recording electrodes. A stereotaxic device (Kopf, Tujunga, CA) was used to secure the skulls of surgical animals. Ground screws (P1 Technologies, Roanoke, VA) were placed above the cerebellum and olfactory regions for stability of the implant and were further connected to a ground wire used for overall signal referencing. Holes were carefully drilled in the skull above target regions, and then specially designed tungsten wire (California Fine Wire Co. Grover Beach, CA) bundles were placed at the desired depth within the brain. Electrode bundles were designed to target the following regions: PFC (PrL and IL), NAc, AMY (MeA, BLA, and CeA), VTA, and VHipp.

Coordinates were guided by prior work in C57BL/6 mice^15^ (Table S1) after being verified or altered slightly to account for differences of the CD1 brain. Regions were chosen based on their relationship to maternal care and their overlap with regions in recent molecular and neurophysiology stress vulnerability studies^15,42^. The only non-overlapping region included in this design is the MeA, a maternal-relevant region that was added on to the AMY bundle. Young adult experimental animals were implanted between 6 and 7 weeks of age. Recovery time was a minimum of 2 weeks prior to the start of any testing post-surgery (Fig. 2a).

### Neurophysiological Data Acquisition

The CerePlex Direct neural signal processor (© Blackrock Neurotech, Salt Lake City, UT) was used to sample low noise neuronal activity at 30 kHz. A bandpass filter of 0.5-250 Hz was applied to the local field potentials (LFPs), which were collected and stored at 1000 Hz. Mice were briefly anesthetized with 5% isoflurane to allow for their efficient connection to a CerePlex µ headstage (© Blackrock Neurotech, Salt Lake City, UT). Mice were always provided with 30 minutes minimum habituation time to adjust to their system connection prior to the start of recording. Once started, neurophysiological recordings paired with time-locked video were obtained over a range of durations spanning 10 minutes to 4 hours during naturally occurring, freely moving behaviors (see below). Experiments were carried out across the maternal transition in three separate cohorts of animals.

### Homecage Recordings

Homecage recordings were collected across the maternal transition for varying minimum durations at the following timepoints: preconception (10 minutes), E17/gestation (10 minutes), P1 (4 hours), P3 (3 hours), P8 (2 hours), P14 (1 hour), and P20 (10 minutes). Due to experimental limitations, longer recording durations were prioritized over the early postpartum period, when maternal care is most essential. Preconception recordings were captured as an important baseline measurement, the late gestation timepoint (E17) was chosen to observe change with pregnancy, and the postpartum timepoints were selected to cover the stages of maternal care occurring across pup development through independence. More timepoints were selected in early development when care is most crucial. Experimental animals had access to food, but not water for the duration of recordings. Pups remained in the cage for all postpartum timepoints except for P20, when they were removed to prevent their disturbance of the recording cable and to eliminate risk of animal injury.

### Timed Pregnancies of Implanted Animals

Once implanted animals were fully healed, sexually mature, and had completed baseline/preconception recordings (preconceptual FIT, homecage, and pup retrieval screen – see below), they were paired with control, non-implanted male CD1 mice at approximately 12 weeks of age for breeding. Following pairing, females were checked for the presence of a vaginal plug each morning to see if successful copulation had occurred. If a vaginal plug was detected, then the male was removed from the cage, and the female’s weight was recorded. The female’s weight was monitored to confirm the pregnancy and conception date, which was noted as embryonic day zero (E0).

### Assessments of Maternal Behavior with Pups

Naturalistic maternal behaviors were assessed in the homecage following electrode implantation and breeding (or with non-implanted controls) as described in the following sections. Pup retrieval behaviors were also evaluated prior to breeding and again in early postpartum as outlined below. Statistics: to determine differences between groups on each naturalistic maternal observation and retrieval test, measurements were assessed via student’s un-paired t-test or a non-parametric equivalent (Mann Whitney) when normality requirements were not satisfied. For tests across multiple maternal timepoints, the two-way repeated measures ANOVA or linear mixed model approach (with unbalanced or missing data) was used to evaluate effects of maternal stage.

### Observations and Scoring of Naturalistic Maternal Behaviors

Postpartum mice were recorded in their homecage alongside pups over multiple observations covering many hours and timepoints of care. During these observations, there were no disturbances to the cage, apart from the dam’s recording electrode being plugged into the neurophysiological recording equipment. The experimenter left the room after the start of the recordings to prevent disruption of natural behavior by the dam. In addition to the neurophysiological data collected, time-locked video of behavior was also captured. This behavior was later hand-scored for various maternal behaviors both “on” and “off” the nest. The experimenter was not blinded to experimental group for the continuous scoring of maternal behavior. However, a blinded experimenter performed maternal behavior scoring at fixed observation timepoints (once every 4 minutes at P3). General “on-nest” maternal behavior was scored across all postpartum timepoints. At P3, the following sub-behaviors were additionally documented: nursing, nest-building, pup retrieval, and licking/grooming of pups^43,44^. Total time on nest and number of care visits were additionally assessed for differences across timepoints and between groups. Dam self-care behaviors such as eating and self-grooming were also scored. Observations were further performed with a cohort of non-implanted animals at P3 to evaluate whether the neural implant altered general “on-nest” maternal behavior.

### Preconception Pup Retrieval Screen

Pup-naïve, adult female mice with neural implants were screened for pup retrieval behavior, which is typically not observed in mice lacking maternal experience^45^. Experimental mice were placed in a clean cage resembling their homecage and left to habituate for 20 minutes. The “clean homecage” had the same bedding and nesting materials present as the subject’s homecage. After 20 minutes, several pups (5-8 in number) between one and four days old were added to the corner of the cage where the clean nesting material had been placed. The experimental mouse was allowed to acclimate to the presence of the pups for two minutes. Once two minutes had passed, one of the pups was moved from the nest to a far point in the cage. The experimental mouse was given two minutes to retrieve the pup back to the nest. If the mouse did not retrieve the pup during the two minutes, then the trial was scored as a failure, and the isolated pup was returned to the nest. This process was repeated for a total of ten trials, and the pup selection and isolation placement in the cage was varied over the course of the trials. If the experimental animal showed any aggression towards the pups at any point following their introduction to the cage, the retrieval screen was stopped immediately. Reliable retrieval was scored as 2 or more successful retrieval trials (out of the 10 total)^46^. In addition to experimenter observation, trials were video recorded, and for implanted animals, neurophysiological data was gathered simultaneously.

### Homecage Pup Retrieval at P4

A pup retrieval assay was conducted in parous mice with the same procedure as the retrieval screen, except the dams’ own pups and homecage were used at P4. Retrieval trials were evaluated on the completeness of retrieval (partial retrieval vs. complete retrieval back to nest), the percentage of complete retrievals, and the average time to retrieve.

### Building *Electome Factors* of Maternal Experience and Behavior

We used homecage LFP data collected over the maternal transition and postpartum period to identify electrical brain network activity features that work together in a coordinated way across maternal stages. These collections of electrical brain network activity features, or electrical network signatures, can be thought of as factors of the functional electrical connectome, which we refer to as *Electome Factors* (*EF*s). LFP data, acquired via neurophysiological recordings, were summarized within eight targeted regions (PrL, IL, NAc, BLA, CeA, MeA, VTA, and VHipp). Within each recording, channels mapped to the same region were averaged to obtain one regional time series per region. Signals were resampled from 1000 Hz to 100 Hz (for predefined frequency bands; defined below) or 200 Hz (for 1 Hz frequency steps) using polyphase resampling with low-pass filtering, and the resampled signals were segmented into consecutive, non-overlapping 3-second windows. For all model development, we used a supervised autoencoder (SAE) with a Non-negative Matrix Factorization (NMF) decoder^47^ to train the model on control data only. The specific SAE approach employed in this study was chosen due to its prior success in identifying stress and social behavior-associated multi-region networks^47,48^. For details on maternal *EF* model development, see Supplemental Methods: “Model development detailed overview (SAE-NMF)”, “Training, validation, and calculation of *Electome Factors*”, and “Feature Thresholding”.

#### Frequency banded networks

For each 3-second window, spectral power and magnitude-squared coherence within three predefined frequency bands (2-7 Hz, 8-12 Hz, and 14-23 Hz) were computed using Welch’s method, yielding 108 spectral features per time window. Feature normalization was performed separately within each recording (mouse x stage). Powers were log-transformed (base 10) and then min-max normalized, and coherences were min–max normalized within the same recording.

#### Networks trained in 1Hz frequency steps

In parallel with the banded-frequency analysis, we also constructed network features using consecutive 1-Hz frequency bins from 2 to 56 Hz. Using the same 3-second non-overlapping windows, spectral power within each region and magnitude-squared coherence between regions were computed in each 1-Hz band using Welch’s method. Aside from replacing the three predefined frequency bands with 1-Hz frequency steps, all preprocessing and normalization procedures were identical to those described above for the banded-frequency networks.

#### Data labeling with behavioral metrics

Three behavioral events were manually annotated from video recordings: “on-nest” behavior, licking, and self-grooming. For each 3-second window, a binary label (1/0) was assigned to each behavior based on whether it occupied more than half the window duration (≥1.5 s). Maternal stage labels (preconception, E17/gestation, P1, P3, P4, P8, P14, P20) were assigned as categorical variables corresponding to each recording session.

#### Maternal Engagement EF training

To identify an *EF* of maternal engagement, we used general “on-vs.-off” nest maternal behavior as the discriminator, which was hand-scored from early postpartum (P1: 4 hour, and P3: 3 hour) video recordings paired with time-locked neurophysiological data. The discriminative performance of the *EF* was summarized using ROC AUC from leave-one-animal-out cross-validation.

#### Maternal Stage EF training

To further identify an *EF* of maternal stage reflective of neuroplasticity over time during the maternal transition, we focused our discrimination across maternal timepoints. Specifically, the preconception stage (10-minute recording) vs. the early postpartum stage homecage recordings (P1: 4 hours, P3: 3 hours, and P4: 20 minutes) were used to distinguish maternal timepoint and train the model on control data.

#### Model testing

Additional out-of-sample evaluation was performed using data collected in parallel but not involved in *EF* training. For *EF_on-nest_*, animals with a history of ELS were used for external validation. For *EF_stage_*, LFP data from maternal timepoints collected but not used in model generation (E17/gestation: 10 minutes, P8: 2 hours, P14: 1 hour, and P20: 10 minutes) were projected onto the trained model to examine how *EF* strength changed across the maternal transition and postpartum timecourse (see “Training, validation, and calculation of Electome Factors” in Supplemental Methods for additional details). All LFP data used in the generation and testing of the new maternal *EF*s was limited to only that which included validated electrode placement for all 8 targeted regions.

### Calculating Stress-related *Electome Factor* Network Scores

In order to evaluate the activity of previously identified *Electome Factor* (*EF*) brain networks related to stress susceptibility^15^ in the experimental animals, we projected LFP data collected during homecage, behavioral, and FIT recordings onto these established models as formerly described (for *EF*s 1-3, 6)^15,42^.

### Forced Interaction Test

The Forced Interaction Test (FIT) for negative affect response to an aggressor was performed as previously described ^49^ with minor adjustments. Immediately following a 10-minute homecage recording, mice were placed in a cylindrical sub-chamber (4.2□ diameter x 8□ high) in the middle of a clean standard cage. The curved surface of the sub-chamber was slotted to allow for sensory engagement with the environment. After 5-minutes in the sub-chamber, either a male CD1 mouse or a female Wistar rat screened for aggressive behavior against female mice was introduced into the cage for an additional five minutes. The sub-chamber protected experimental mice from physical harm while forcing them into sensory contact/exposure with the more intimidating animal. Neurophysiological and video recordings were collected during the FITs to evaluate brain activity and monitor behavior during this encounter. This assay was performed at both preconception and early post-weaning timepoints.

### *Electome Factor* Score Analysis

To analyze *Electome Factor* (*EF*) differences by group (control, ELS) or changes over time in network strength, we used mean *EF* scores collapsed by observation timepoint and further sorted by time-locked behavior when applicable. Prior to statistical evaluation, mean *EF* scores were fourth root transformed to improve data normalization^42^. *EF* scores sorted by on/off nest behavior at a single timepoint were analyzed using the paired t-test (2 behavioral conditions, 1 animal condition), one-way repeated measures (RM) ANOVA (3 behavioral conditions, 1 animal condition), or two-way RM ANOVA (2 behavioral conditions, 2 animal conditions). These analyses were carried out using GraphPad Prism. *EF* scores evaluated across 2+ timepoints were affected by missing datapoints. Such maternal timecourse-relevant assessments were carried out using the one, two, or three-way RM Restricted Maximum Likelihood (REML) mixed-effects approach depending on whether animal condition (control, ELS) or behavior (i.e., “on” vs. “off-nest”) were included in the analysis design. If a main effect involving maternal timepoint was detected, it was followed up via multiple comparisons testing, with False Discovery Rate (FDR) corrected p-values being reported throughout.

### Histological Validation of Electrode Placements

Following the completion of all experiments, implanted subjects underwent cardiac perfusion under deep anesthesia (pentobarbital, 150mg/kg, Sagent Pharmaceuticals, Schaumberg, IL) with 4% PFA in PBS, pH 7.4. Brains were dissected from skulls and stored in 4% PFA for 24 hours before being cryoprotected in 30% sucrose for an additional 24 hours, or until they had sunk in solution. The cryoprotected brains were subsequently frozen down in Tissue-Plus® O.C.T. Compound (Thermo Fisher Scientific, 23730571, Waltham, MA) on dry ice and stored at −80°C until being sliced via cryostat (Leica Biosystems, Deer Park, IL) at −20°C into 35µm sections. Brain sections were stained with NeuroTrace™530/615 red fluorescent Nissl Stain (Thermo Fisher Scientific, N21482, Waltham, MA) and mounted onto Superfrost Plus™ microscope slides (Thermo Fisher Scientific, 1255015, Waltham, MA) for imaging. A condition-blinded experimenter imaged and evaluated brain sections to determine the accuracy of implanted electrode placements. This experimenter also did not take part in neurophysiological or behavioral data collection or analysis. Neurophysiological data analysis was revised as needed following the finalized histological evaluation of all electrode placements.

## Results

### Modeling brain network function from observations of naturalistic maternal behavior

We utilized a supervised autoencoder (SAE) approach to investigate a predictive relationship between brain-wide spatiotemporal dynamics and naturalistic maternal behavior. This approach identifies electrical dynamic features from across multiple brain regions which define a network that is highly predictive of outcomes or behaviors^47,48^. This process involves the training of a simultaneous collection of latent networks, where one network emerges as the most highly predictive. We applied this method to a dataset of simultaneous LFP recordings from eight brain regions (Fig. 1a) to identify maternal-relevant brain networks. These networks are hereafter referred to as *Electome Factors* (*EF*s), where “*Electome*” represents the functional electrical connectome. Only animals with verified electrode placement in all eight regions were used to train these models (Supplemental Fig. S1). All neurophysiological recordings were paired with time-locked video for simultaneous behavioral assessment. Behavioral observations in non-implanted dams confirmed that the implants did not significantly alter general maternal care behavior (Supplemental Fig. S2).

**Figure 1:**
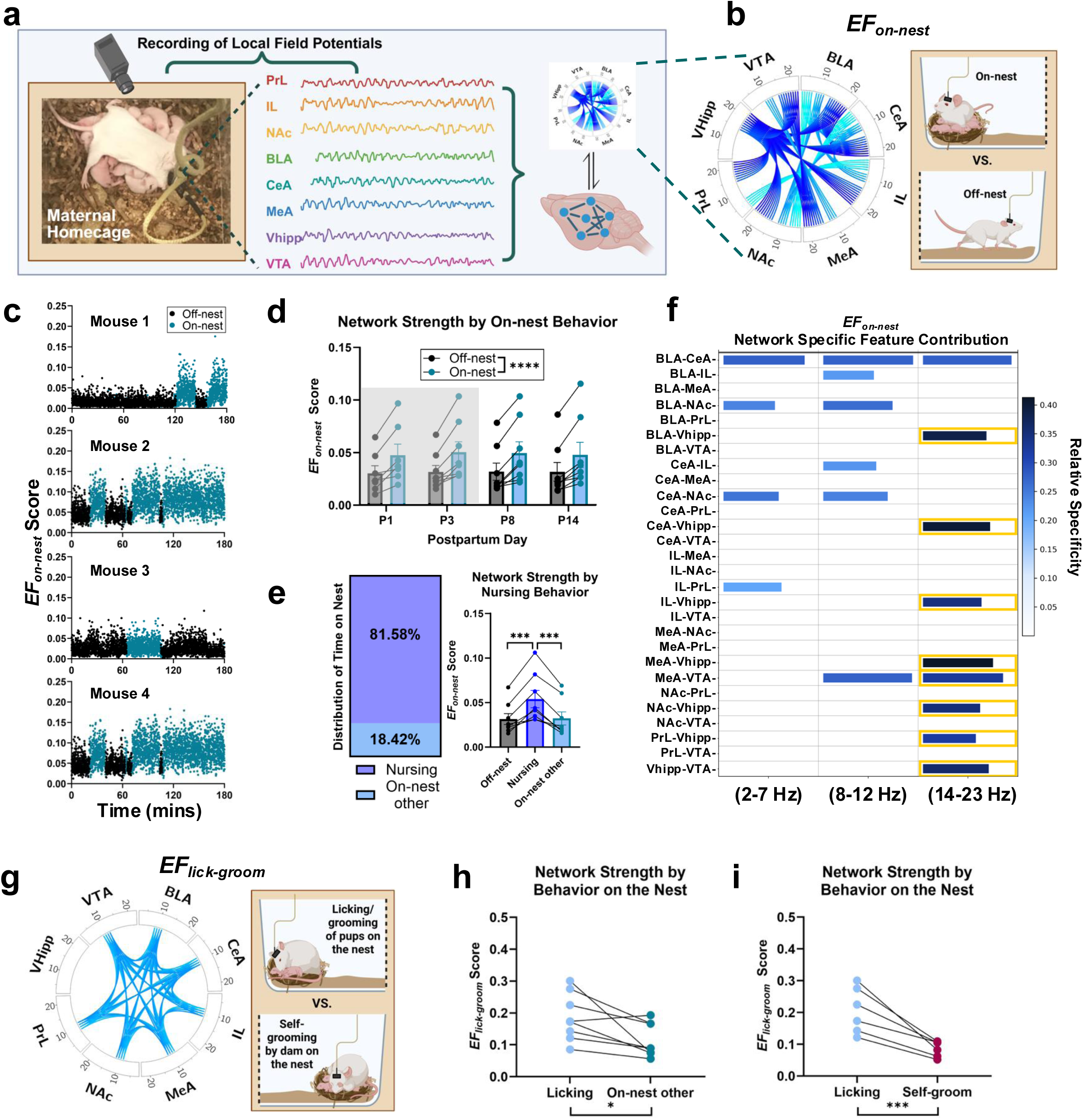
Identifying a novel *Electome Factor* of “on-nest” maternal behavior. a, Schematic showcasing the general project methodology. b, Circos plot for the “on-nest” *Electome Factor* (*EF_on-nest_*) and visual depiction of the behavioral classifier that was used (“on” vs. “off-nest” behavior) to train a network for “on-nest” behavioral discrimination. The regions included in the network are presented around the outside of the Circos plot alongside corresponding frequency bands. Any regional power contributing to the network is shown within the outside band of the plot (none present here). Synchronous regions with contributing network coherence are indicated by connections across the center of the plot. The frequency bands are color-coded moving from lighter to darker blue with increasing frequency. c, Four example traces of *EF_on-nest_* strength scores paired with associated “on” (blue) or “off-nest” (black) observed location over a 3-hour observation time window at P3. Each dot represents the network strength over a 3-second time bin. d, Mean *EF_on-nest_* strength scores separated by “on” and “off-nest” behavior location across four observed postpartum timepoints. The grey background indicates the timepoints from which data was used for network training. There is a significant main effect of behavior location (REML, *P* = 4.25E-15, *n* = 7-8), with *EF_on-nest_* scores increasing when dams are “on-nest” at each timepoint of observation (Table S2). e, Percent of time on the nest spent nursing at P3 and mean *EF_on-nest_* score separated by “off-nest”, nursing (a subset of “on-nest”), and “on-nest-other” behavior. There is a significant effect of behavior subtype (RM ANOVA, *p* < 0.0001, *n* = 8), with the EF*_on-nest_* score elevated during nursing as compared to “off-nest” and “on-nest other” (Table S3). f, Network specific feature contribution for *EF_on-nest_* sorted by guided frequency bands. The length of the bar indicates absolute strength of network contribution, and depth of color represents specificity of the feature’s contribution to the “on-nest” network. The yellow outlines indicate features with significant network contribution through absolute strength and specificity. g, Circos plot for the “lick-groom” *Electome Factor* (*EF_lick-groom_*). This network was trained using the behavioral classifier of dams “licking” pups on the nest vs. dams grooming themselves on the nest. One control animal was excluded from training due to extreme class imbalance. The Circos description provided for Panel b is also applicable here. h, Strength of *EF_lick-groom_* during licking (of pups) on the nest vs. all other behavior on the nest to assess behavioral discrimination of the network (paired t-test, *P* = 0.0150, *n* = 8). i, Strength of *EF_lick-groom_* during licking (of pups) on the nest vs. self-grooming on the nest (paired t-test, *P* = 0.0008, *n* = 7). Data shown are mean ± sem. \**P* < 0.05, \*\*\**P* < 0.001, \*\*\*\**P* < 0.0001. All statistical analyses of mean *EF* scores were performed on fourth root transformed data. This was done to help stabilize variance and make the data more normally distributed. All data shown are non-transformed. Statistical analyses include the two-way RM Mixed-Effects Model [REML] (d), the one-way RM ANOVA (e), and the paired t-test (h-i). If a significant effect of the repeated measure was found, a False Discovery Rate (FDR) correction for multiple comparisons was performed to specify which timepoints or measures were significantly different. PrL = Prelimbic Cortex, IL = Infralimbic Cortex, NAc = nucleus accumbens, BLA = basolateral amygdala, CeA = central amygdala, MeA = medial amygdala, VHipp = ventral hippocampus, VTA = ventral tegmental area, *EF* = *Electome Factor*, P = Postpartum Day, REML = restricted maximum likelihood, RM = repeated measures, ANOVA = analysis of variance, FDR = False Discovery Rate correction for multiple comparisons. Panel a, Created in BioRender. Mitchell, SB. (2026) https://BioRender.com/6vkvhof. Panel b, Created in BioRender. Mitchell, SB. (2026) https://BioRender.com/589usn1. Panel g, Created in BioRender. Mitchell, SB. (2026) https://BioRender.com/5pxfmxo.

We started with the most general behavioral assessment of maternal care: dams’ engagement with pups, on the nest, or away from pups, off the nest. Homecage data were therefore labeled as “on” or “off” the nest over observation epochs, and a model network was trained on this labeled data of neurophysiological recordings from early postpartum timepoints (P1, P3). For training this model, pre-determined guided oscillatory frequency bands of 2-7 Hz, 8-12 Hz, and 14-23 Hz were utilized given their effectiveness in prior network dynamic analyses using the regions recorded from in this study^49^. These guided frequency bands capture directional information flow between pairs of brain regions, have clear biological meaning ^49^, and enabled us to approach this work with a reasonable set of parameters to identify reliable models of complex, naturally occurring behaviors over the long course of the maternal experience. Such a strategy is consistent with human studies in which brain activity is monitored over longer periods by EEG, with roughly parallel delta/theta, alpha, and beta frequency bands ^50,51^. Novel network activity predictive of “on-nest” maternal behavior was identified (Fig. 1a-b) that had greater strength during “on-nest” behavior (Fig. 1c-d, Table S2); the activity of this network was determined to discriminate “on” vs. “off-nest” behavior using a leave-one-out (LOO) testing approach (receiver operating characteristic area under the curve [AUC] mean ± sem = 0.71 ± 0.03, *n* = 8). This network was further validated using P8 recordings from the same animals (AUC mean ± sem = 0.72 ± 0.03, *n* = 8). We named this the “on-nest” *Electome Factor* (*EF_on-nest_*) (Fig. 1b, Supplemental Fig. S3). Further evaluation of network strength found *EF_on-nest_* to primarily reflect engagement in nursing behavior (Fig.1e, Table S3).

The *EF_on-nest_* network comprises 19 selected features, with coherence between all eight brain regions and across all guided frequency bands providing strength contributions (Fig. 1b and Supplemental Fig. S3). Coherence between IL and PrL, as well as between AMY and NAc, were prominent network aspects in the 2-7 Hz and 8-12 Hz bands, and intra-amygdala coherence between the BLA and CeA contributed to the network across all three frequency bands. Coherence involving the VTA and VHipp contributed more within the 14-23 Hz frequency band. Notably, coherence of VHipp with all other recorded brain regions contributed significantly to EF_on-nest_ in the 14-23 Hz range, indicating that this region plays a particularly important role in *EF_on-nest_* at those higher frequencies. While power within each region did not contribute substantially to *EF_on-nest_*, the wide range of contributing coherence across regions and frequency bands highlights the importance of the brain-wide network approach when aiming to understand maternal-relevant neural dynamics. To better comprehend contributing features of *EF_on-nest_*, we also considered whether features were highly specific to this network (Fig. 1f). During training, many latent networks were identified having varying predictive performance for “on-nest” behavior, with *EF_on-nest_* performing best. Feature specificity to *EF_on-nest_* was therefore evaluated relative to other latent networks that had poorer prediction in this context. For instance, the 14-23 Hz coherence of VHipp with all other regions showed definitive specificity in addition to significant absolute strength contributions to *EF_on-nest_*.

Models were trained using guided frequency bands for scientific reasons, yet this approach leaves the possibility of overlooking important network feature combinations outside of these ranges. As such, we examined an *EF* network trained in 1Hz steps (*EF_on-nest-1-Hz_*), which contained vastly more parameters but yielded largely similar results for both the degree of prediction achieved and features identified. A single exception arose for *EF_on-nest-1-Hz_*, which included additional low-frequency power contributions across regions (Supplemental Figs. S4-S5, Supplemental Results). Because the approach with guided bands and fewer parameters enabled us to more reliably achieve generalizable results, for the remainder of our studies, we report networks trained in this way. More information on the 1Hz-trained networks can be found in the supplement [Supplemental Figs. S4-S5, S14-S15, Supplemental Results].

To more fundamentally examine maternal behavior in a manner independent of nest environment, we turned to maternal licking, a primary behavior demonstrated in maternal care^8^. We trained *EF*s by discriminating licking from other “on-nest” behaviors (Supplemental Results). The first network looked at licking and grooming behavior discriminating between the dam licking/grooming pups vs. the dam grooming itself (*EF_lick-groom_*, Fig1g, Supplemental Fig. S9-10). This network had moderate performance (LOO AUC mean ± sem = 0.64 ± 0.04) and was successful in its intended behavioral discrimination (Fig. 1h-i). In observing selected contributing features of *EF_lick-groom_*, the 8-12 Hz oscillatory frequency band was essential, as all 18 of the identified network features were found to lie within this range (Supplemental Fig. S10). *EF_lick-groom_* excluded VHipp contributions in stark contrast to *EF_on-nest_*. Given the limited occurrence of maternal self-grooming, another network contrasting maternal licking and grooming of pups vs. the more abundant non-licking neural activity was trained (*EF_licking_*, Supplemental Fig. S6-8). This *EF_licking_* network performed well (LOO AUC mean ± sem = 0.78 ± 0.05)(Supplemental Fig. S7), but could not discriminate different kinds of licking (Supplemental Fig S8). This network also did not contain VHipp contributions but had features shared with both *EF_lick-groom_* and *EF_on-nest_*. The contributors to these *EF*s continue to demonstrate the importance of inter-region coherence from across multiple brain regions in the discriminative performance of identified networks underlying maternal behavior.

### Maternal stage as a classifier for training discriminative maternal brain networks

Another major motivation for the design of this study was to capture the adaptive nature of the maternal brain by measuring brain-wide spatiotemporal dynamics across the pregnancy and postpartum transitions, as done in multi-time point human studies ^1,2^. Therefore, LFP data collection began prior to conception and continued through gestation to several observational timepoints across postpartum (Fig. 2a). Although we found no difference in the percentage of time spent on the nest across postpartum days, we did observe a stage-dependent relationship of *EF_on-nest_* network strength with time on the nest (Supplemental Fig. S11). This finding supports the idea that the strength of behavior-trained networks can change over time, independently from an animal’s level of engagement in that behavior. Hence, the strength of the maternal behavior-associated networks (*EF_on-nest,_ EF_licking,_* and *EF_lick-groom_*) was evaluated across all recorded timepoints (Fig. 2b-c, Supplemental Fig. S12, Tables S4-S5).

**Figure 2:**
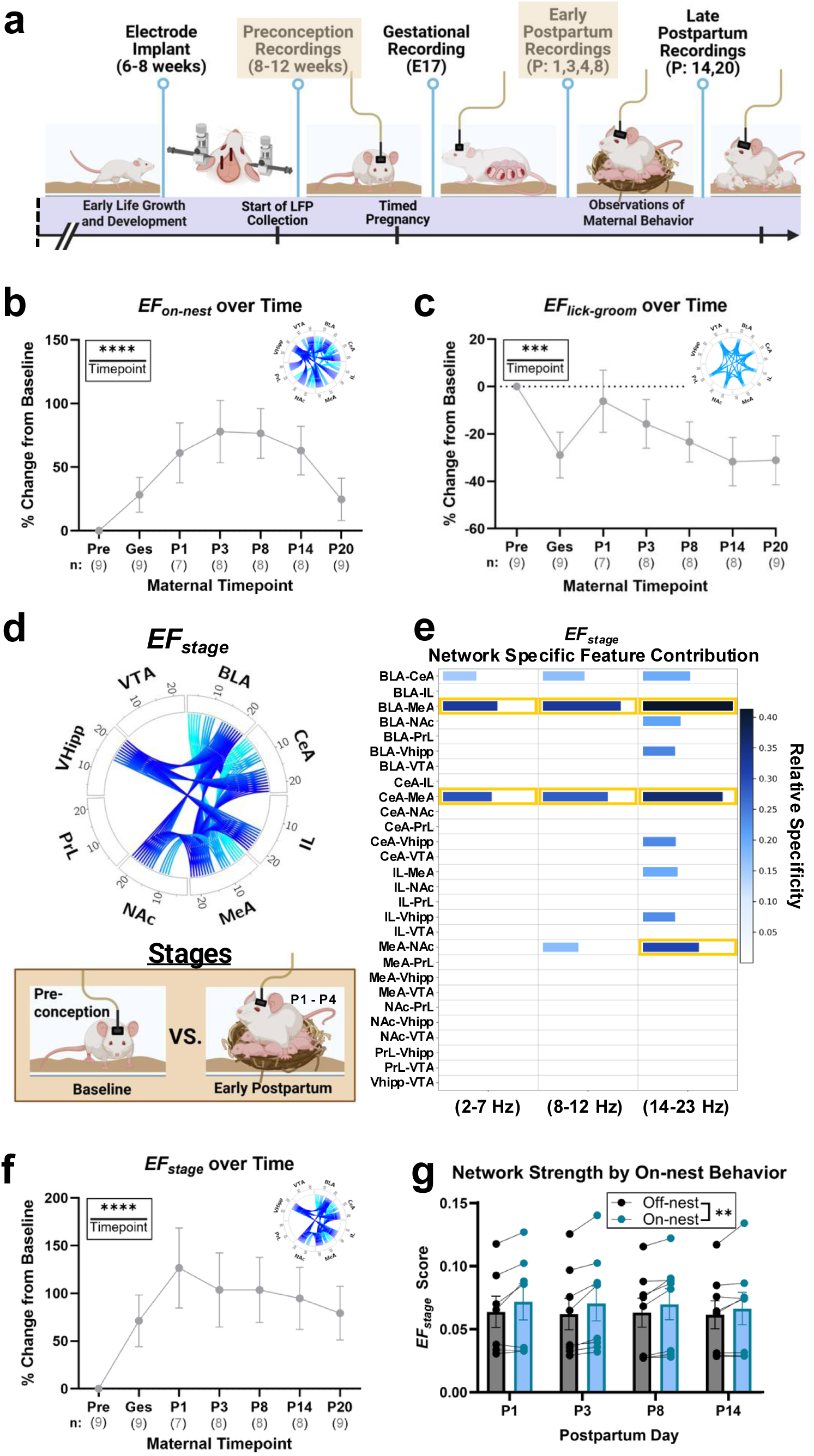
Maternal-relevant *EF*s display variable strength trajectories according to maternal stage. a, Schematic presentation of the experimental timeline from adolescence through maternal adulthood. Highlights at preconception and early postpartum indicate the recording timepoints used for training *EF_stage_*. b, *EF_on-nest_* strength scores across observed maternal timepoints normalized to the preconception baseline level (REML, timepoint effect, *P* = 6.99E-05, *n* = 7-9). Multiple comparisons of individual timepoints can be found in Table S4. c, *EF_lick-groom_* strength scores across observed maternal timepoints normalized to the preconception baseline level (REML, timepoint effect, *P* = 4.00E-04, *n* =7-9). Multiple comparisons of individual timepoints can be found in Table S5. d, Circos plot detailing the features of *EF_stage_* alongside a schematic depiction of the stage-wise classifiers (preconception baseline vs. early postpartum: P1, P3, P4 homecage) that were used in training a discriminative network of maternal stage. The description provided under Fig. 1b is also applicable here. e, Network specific feature contribution for *EF_stage_* sorted by guided frequency bands. The description provided under Fig. 1f is also applicable here. f, *EF_stage_* score across observed maternal timepoints normalized to the preconception baseline level (REML, timepoint effect, *P* = 4.17E-06, *n* = 7-9). Multiple comparisons of individual timepoints can be found in Table S6. g, Mean *EF_stage_* score separated by “on” and “off-nest” behavior location across observed maternal timepoints (REML, behavior location effect, *P* = 0.0092, *n* = 7-8). Individual timepoints do not show significance following multiple comparisons (Table S7). Data shown are mean ± sem.; \*\**P* < 0.01, \*\*\**P* < 0.001, \*\*\*\**P* < 0.0001. All statistical analyses of mean *EF* scores were performed on fourth root transformed data. This was done to help stabilize variance and make the data more normally distributed. For the normalized *EF* trajectory data (b-c, f), the transformation was performed prior to calculating the percent change from baseline. All data shown are non-transformed. Statistical analyses include the one-way RM REML (b-c, f) and the two-way REML (g). If a significant effect of the repeated measure was found, an FDR correction for multiple comparisons was performed to specify which timepoints or measures were significantly different. PrL = Prelimbic Cortex, IL = Infralimbic Cortex, NAc = nucleus accumbens, BLA = basolateral amygdala, CeA = central amygdala, MeA = medial amygdala, VHipp = ventral hippocampus, VTA = ventral tegmental area, *EF* = *Electome Factor*, E = embryonic day, P = Postpartum Day, REML = restricted maximum likelihood, RM = repeated measures, FDR = False Discovery Rate correction for multiple comparisons. Panel a, Created in BioRender. Mitchell, SB. (2026) https://BioRender.com/60gmj72. Panel d, Created in BioRender. Mitchell, SB. (2026) https://BioRender.com/e7h1elo.

We found the *EF_on-nest_* network to increase in strength from the preconception timepoint until P3 (Fig. 2b, Table S4). The strength of the network then remained at a similar level at P8 before dropping throughout late postpartum. *EF_lick-groom_* network strength also changed across maternal stages, with an unexpected highest average strength observed in preconception as compared to gestation and postpartum (Fig. 2c, Table S5). These findings suggest that the time course of maternal plasticity is an important aspect of brain network activity underlying maternal behavior.

With this in mind, we sought to identify an additional *EF* specifically targeting maternal stage, independent of behavior. Our hypothesis was that a network trained to predict stage would have a unique set of selected contributing features, with some overlapping prediction of maternal behavior. Preconception neural data recorded in the homecage was set against early postpartum neural data to train a novel network highly discriminative of maternal stage (Fig. 2d-e; AUC (mean ± sem) = 0.80 ± 0.03, *n* = 8). This network was validated using neural data from all other maternal timepoints collected in concert with, but not included in, the training set (Fig 2a). For this novel “maternal stage” *EF* (*EF_stage_* Fig. 2d-e, Supplemental Fig. S13), 16 features were found to have significant network contributions, including intra-amygdala coherence contributions across all guided frequency bands. In particular, coherence between BLA and MeA and between CeA and MeA were identified as significant network contributors within all guided bands—contributing to both absolute strength and relative network specificity of the emergent *EF_stage_* (Fig. 2e). Similarly, MeA to NAc coherence was also an important network feature, with a contribution of network specificity found within the 14-23 Hz band (Fig. 2e). Additional 14-23 Hz regional coherence contributed to *EF_stage_*: the BLA with NAc and VHipp, the CeA with VHipp, and the IL with MeA and VHipp (Fig. 2e). Overall, six of the eight brain regions we recorded from were found to contribute to *EF_stage_* (all but PrL and VTA). This network revealed clear and unique differences in its contributing features as compared to the maternal behavior (“on-nest”, “licking”, and “lick-groom”) *EF*s.

Looking at network engagement across the maternal timecourse, the strength of *EF_stage_* rose sharply from preconception levels to peak at P1, before gradually reducing over the course of postpartum (Fig. 2f, Table S6). Unlike *EF_on-nest_*, the strength of the “maternal stage” network did not return close to preconceptual baseline levels by the time pups were of weaning age (Fig. 2f). Furthermore, while the overall strength of *EF_stage_* increased when postpartum dams were engaged in “on” vs. “off” the nest behavior (Fig. 2g), the effect was limited (Table S7) when compared to the dramatic shift observed during “on-nest” behavior for *EF_on-nest_* (Fig. 1e, Table S2). Like the other *EFs*, *EF_stage_*was associated with behavior (Fig. 2g, Supplemental Fig. S32d). However, *EF_stage_* had distinct network-specific contributing features from *EF_on-nest,_ EF_licking,_* and *EF_lick-groom,_* emphasizing its uniqueness.

### Stress-associated *EF*s and the maternal transition

Using similar computational strategies, previous work has established the identity of six stress-relevant brain-wide *EF*s in male C57BL/6 mice^15^. While we trained our novel maternal *EF*s to discriminate by maternal behavior and stage, the prior stress *EF*s were trained to discriminate different aspects of stress such as chronic stress exposure, susceptibility vs. resilience to chronic social stress, and acute negative affect response^15^. Because the present study recorded from brain regions underlying stress vulnerability as well as maternal care behavior, we were able to measure stress-relevant *EF* strength across the maternal timecourse for four of these networks (Fig. 3a). Together, these findings demonstrate broad applicability of stress-relevant *EF* strength to females and the CD1 mouse strain, and provide a basis to further test the impact of the maternal timecourse on stress-relevant *EF*s.

**Figure 3:**
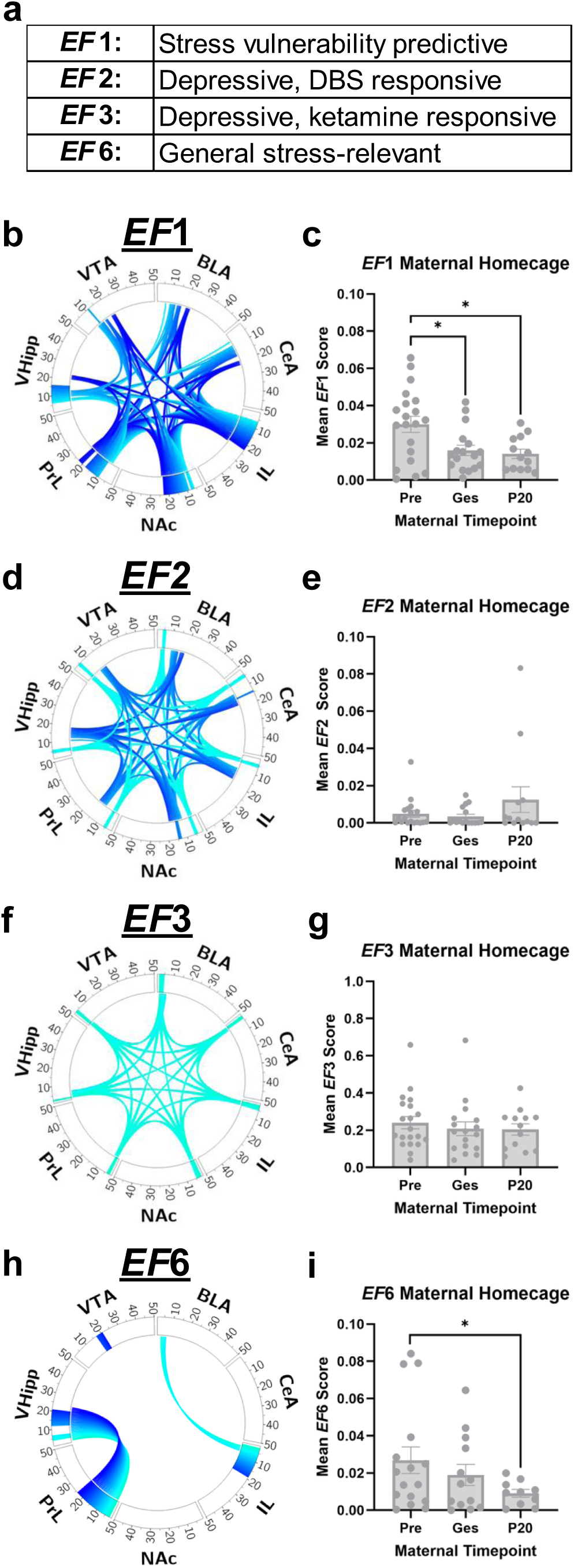
Stress-relevant *EF*s show altered engagement with maternal transition. a, Table listing known characterizations of the previously determined stress-relevant *EF*s. Only those for which data is presented are included. b,d,f,h, Circos plots detailing the network specific features for the stress-relevant *EF*s b, *EF*1, e, *EF*2, g, *EF*3, h, *EF*6. c,e,g,i, Mean stress *EF* scores across observed maternal-relevant timepoints and pooled behavior without pups present for c, *EF*1: (REML, timepoint effect, *P* = 0.0349, *n* = 13-20), with *EF*1 decreased during gestation and late postpartum (P20) as compared to preconception (Table S8), e, *EF*2, g, *EF*3, i, *EF*6: (REML, timepoint effect, *P* = 0.0263, *n* = 9-20), with *EF*6 decreased in late postpartum (P20) as compared to preconception (Table S9). Data are mean ± sem.; \**P*<0.05. Statistical analyses include the one-way REML (c, e, g, i). If a significant effect of the repeated measure was found, an FDR correction for multiple comparisons was performed to specify which timepoints or measures were significantly different, as indicated by the significance bars in the plots above. All statistical analyses of mean *EF* scores were performed on fourth root transformed data. This was done to help stabilize variance and make the data more normally distributed. All data shown are non-transformed. PrL = Prelimbic Cortex, IL = Infralimbic Cortex, NAc = nucleus accumbens, BLA = basolateral amygdala, CeA = central amygdala, MeA = medial amygdala, Vhipp = ventral hippocampus, VTA = ventral tegmental area, *EF* = *Electome Factor*, P = Postpartum Day, Pre = preconception, Ges = Gestation, REML = restricted maximum likelihood, RM = repeated measures, FDR = False Discovery Rate correction for multiple comparisons

*EF*1 (Fig. 3b) is the most highly characterized stress *EF* to date^15^ and is a well-validated indicator of stress vulnerability in C57BL/6 mice that increases in response to negative affective stimuli (restraint or aggressor) in both C57 and CD1 mice (Fig. 6d). When evaluated across the maternal transition, *EF*1 was substantially affected by maternal stage, with *EF*1 scores decreasing from preconception to later stages (Fig. 3c, Table S8). *EF*2 and *EF*3 represent two depressive-associated stress *EF*s. Looking at *EF*2 (Fig. 3d), we did not find a significant effect of maternal stage across homecage timepoints without pups (Fig. 3e). The same was true for *EF*3 over the maternal timecourse (Fig. 3f-g). *EF*6 (Fig. 3h) is a general stress-relevant network that has not been thoroughly characterized, which we hypothesized would be altered by maternal adaptation. When examining the dynamic strength of this network across three maternal timepoints (Fig. 3i), we found it significantly decreased across maternal stage (Table S9). This suggests a link between maternal experience and stress-relevant brain dynamics.

### Stress networks during postpartum behavior and influences of early life stress

Along with the dynamic nature of the plastic and adaptive maternal brain, we were also interested in exploring the maternal transition and subsequent postpartum period as one of increased neuropsychiatric risk^31^. Early life stress (ELS) induces latent brain changes that can emerge when unmasked by a subsequent stressor^52^ (such as pregnancy/postpartum), leading to depressive or other maladaptive states in adulthood ^30,53,54^ Exposure to ELS has been shown to alter brain gene expression, neural circuit activity, and social behavior long term^54^, including maternal behaviors^30^.

To test the impact of ELS on maternal brain network activity and associated care behavior, we recorded from a new set of animals exposed to a combination ELS approach that included maternal separation, early weaning, and limited nesting material (Fig. 4a). Exposure to ELS did not impact animal growth (Supplemental Figure S16). Surprisingly, ELS had no impact on the predictive performance nor strength of *EF*s trained to discriminate by maternal behavior or stage (Supplemental Figures S17-S20, Tables S10-S15). In fact, data collected from ELS dams provided a means of out-of-sample validation of our unstressed control-trained networks, showing similar discrimination AUCs (*EF_on-nest_* P1, P3, P8 pooled AUC = 0.70 ± 0.05; *EF_licking_* P3 on-nest pup licking vs. other AUC = 0.71 ± 0.08; *EF_lick-groom_* P3 on-nest pup licking vs. self-grooming AUC = 0.64 ± 0.08; *EF_stage_* preconception vs. P1, P3, P4 homecage pooled AUC = 0.77 ± 0.07). The similarity of network activity over time for control and ELS groups was matched by their similarity of network activity during maternal behavior on the nest (Supplemental Figs. S17-S20, Tables S10-S15). This suggests that these novel maternally-relevant networks are aligned with behaviors across distinctly different groups.

**Figure 4:**
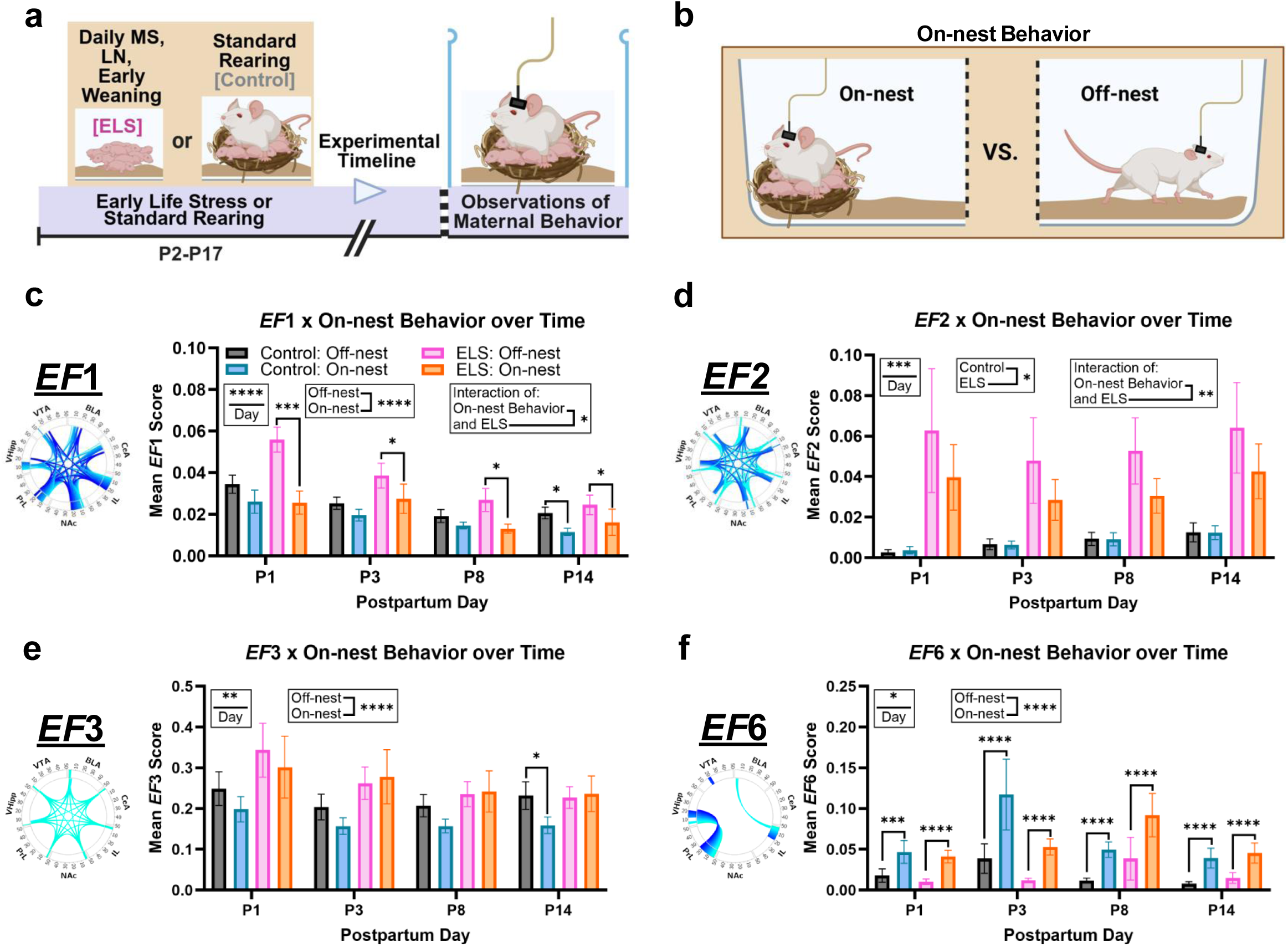
Stress-relevant *EF*s display dynamics in engagement with maternal behavior, which varies with prior ELS-exposure. a, Schematic presentation of the ELS paradigm experienced by a subset of experimental animals prior to their maturation, implantation, breeding, and recording. b, Schematic depiction of “on-nest behavior location”: indicating a dam’s presence “on” or “off” the nest. c,d,e,f, Circos plots and mean stress *EF* scores across observed postpartum timepoints taking into account potential differences by “on” vs. “off-nest” behavior location for c, *EF*1: (REML, day effect, *P* = 7.22E-10; location effect, *P* = 5.69E-11; location x ELS interaction, *P* = 0.0335 *n* = 11-17), d, *EF*2: (REML, day effect, *P =* 1.38E-04; ELS effect, *P* = 0.0251; location x ELS interaction, *P* = 0.0054, *n* = 9-16), e, *EF*3: (REML, day effect, *P* = 0.0032; location effect, *P* = 1.2E-05), f, *EF*6: (REML, day effect, *P* = 0.0107; location effect, *P =* 1.65E-29). Data shown are non-transformed mean ± sem.; \**P*<0.05, \*\**P*<0.01, \*\*\**P*<0.001, \*\*\*\**P*<0.0001. Statistical analyses include the three-way RM REML (c-f). If a significant effect of the repeated measure was found, an FDR correction for multiple comparisons was performed to specify which timepoints or measures were significantly different. All statistical analyses of mean *EF* scores were performed on fourth root transformed data. This was done to help stabilize variance and make the data more normally distributed. All data shown are non-transformed. MS = maternal separation, LN = limited nesting material, ELS = early life stress, PrL = Prelimbic Cortex, IL = Infralimbic Cortex, NAc = nucleus accumbens, BLA = basolateral amygdala, CeA = central amygdala, MeA = medial amygdala, Vhipp = ventral hippocampus, VTA = ventral tegmental area, *EF* = *Electome Factor*, P = Postpartum Day, REML = restricted maximum likelihood, RM = repeated measures, FDR = False Discovery Rate correction for multiple comparisons. Panel a, Created in BioRender. Mitchell, SB. (2026) https://BioRender.com/h436932. Panel b, Created in BioRender. Mitchell, SB. (2026) https://BioRender.com/87jivru.

When looking across the maternal timecourse, we did find that ELS substantially influenced some stress-relevant *EF*s (Fig. 4, Supplemental Fig. S21, Tables S16-S18), which became especially evident when evaluating their strength during postpartum “on-nest” behavior. Over the postpartum period (Fig. 4c, Table S19), we once again detected an effect of maternal timepoint on *EF*1, with a decrease in mean *EF*1 score observed over time. Interestingly, a significant effect of “on” vs. “off-nest” behavior was also found for *EF*1, with its strength notably higher when dams were off the nest (Fig. 4c, Table S20). Given prior work showing that *EF*1 increases with acute negative affect^15^ (Fig. 6c-d), engagement in maternal on-nest behavior appears to suppress neurophysiology of negative affect in this context. While there was no significant effect of ELS, there was an interaction of ELS with “on-nest” behavior across postpartum observations, whereby dams with prior ELS-exposure showed a greater discrepancy between their “off” and “on-nest” *EF*1 scores. For these ELS-exposed dams, down-regulation of *EF*1 activity while on-nest may therefore be an adaptive phenomena, especially when considering that ELS-exposure did not robustly alter on-nest behaviors (Supplemental Fig. S26-28, Tables S36-S37).

For *EF2*, a depressive-relevant network^15^, there was an effect of postpartum day detected, with a general elevation of *EF*2 over time (Fig. 4d, Table S21). There was also a robustly significant effect of ELS, with ELS-exposed dams having increased *EF*2 strength over postpartum observation days, and a significant interaction of ELS with “on-nest” behavior, wherein the ELS-exposed dams once again showed a greater change in *EF*2 strength according to being “on” or “off” the nest. There was no main effect of “on-nest” behavior on *EF*2 strength when considering both control and ELS-exposed dams together. For *EF3*, a second depressive-relevant network^15^, we identified a significant effect of postpartum day, this time with a decrease in strength moving across observation timepoints (Fig. 4e, Table S22). *EF3* network strength was also lower when dams were on the nest (Table S23), similar to what was seen with *EF*1. EF6, a largely uncharacterized general stress-relevant network^15^, had distinct patterns from *EF*s 1-3. While there was a significant timepoint effect detected across postpartum observations for *EF*6 (Fig. 4f, Table S24), the more striking and clear finding was the effect of “on” vs. “off-nest” behavior which diverged from the pattern found in *EF*1 and *EF*3, as the strength of *EF*6 during postpartum “on-nest” behavior increased (Table S25). This finding may help to characterize the relevance of *EF6* to different states.

### ELS-exposure, pup retrieval behavior, and stress-vulnerable network activity

Given the brain-network differences we identified in relation to naturalistic maternal behavior (Fig. 4, Supplemental Fig. S21) and emergent brain states (Supplemental Fig. S25), we next wanted to examine the responsiveness of network activity in the maternal brain during more controlled assays of active maternal care behavior at two experimental timepoints (Fig. 5a). We chose a standard pup retrieval paradigm (Fig 5b)^46^ to probe this question, as it presents a distinct demonstration of maternal behavior involving aspects of motivation and social interaction.

**Figure 5:**
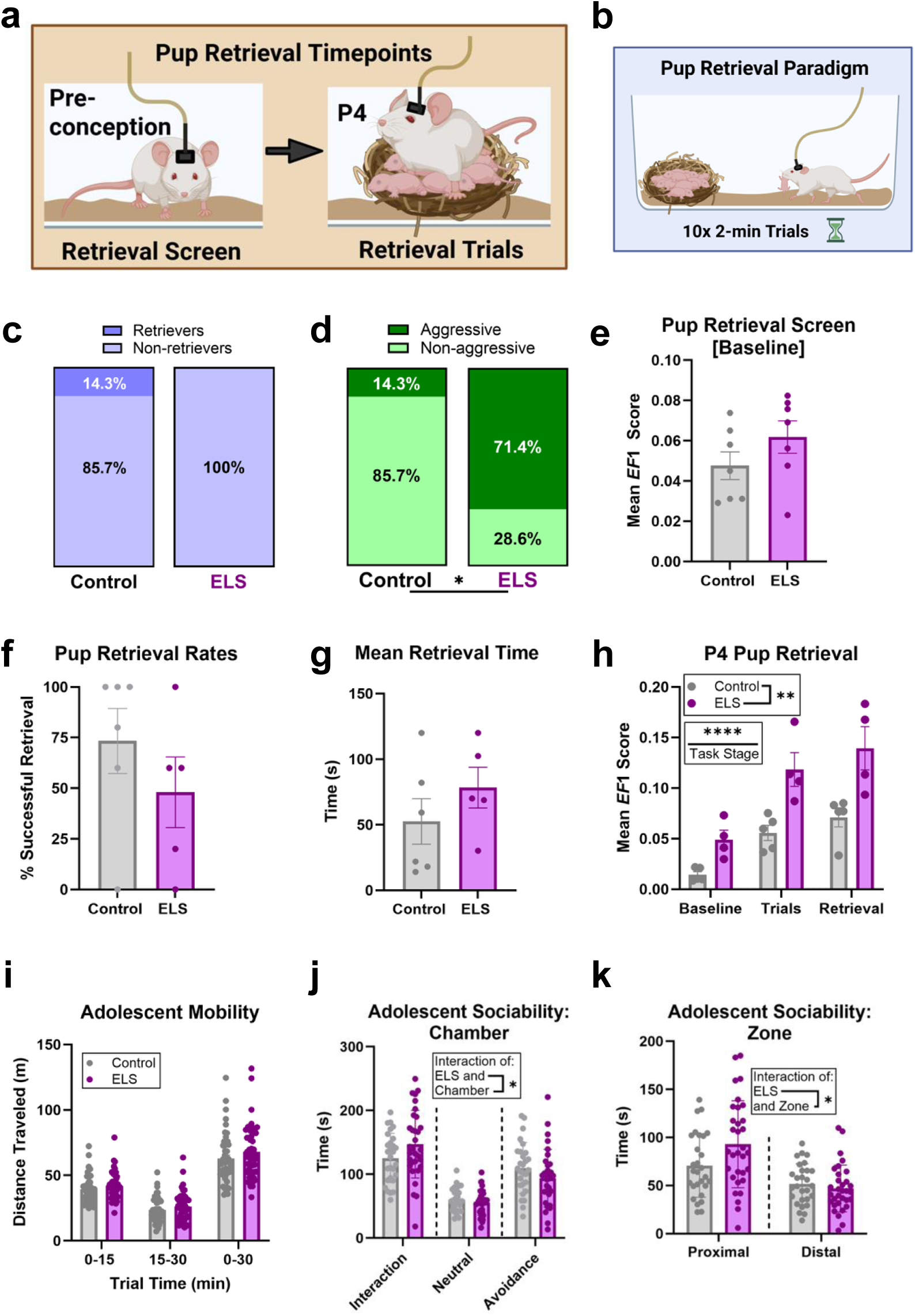
Prior ELS-exposure sensitizes the stress vulnerability network during pup retrieval while subtly impacting social behavior from adolescence. a, Schematic detailing the timepoints of the pup retrieval task. A pup retrieval screen was performed at a pre-conception (pup-naïve) timepoint as well at an early postpartum timepoint (P4) using a dam’s own pups. b, Schematic summarizing the pup retrieval paradigm. c-e) A preconception pup retrieval screen was performed with virgin, pup-naïve adult females (*n* = 7/group). c) Neither control nor ELS-exposed virgin animals reliably retrieved pups d) Some virgin animals displayed aggression towards pups, which was significantly higher in the ELS-exposed animals (*P*=0.0253). e) The mean baseline *EF*1 score for control and ELS-exposed animals prior to the start of the pup retrieval screen. f-h) A pup retrieval task using a dam’s own pups was performed at P4 (*n* = 5-6/group) f) Percentage of successful pup retrieval trials out of 10 total trials. g) Mean time taken to retrieve a pup back to the nest. Unsuccessful trials were included as 120 seconds. h) Mean *EF*1 score for control and ELS-exposed animals during baseline, retrieval trials, and active retrieval behavior (ANOVA, stress effect, *P* = 0.0031; task stage effect, *P* < 0.0001, *n*=4-5/group). The “baseline” stage refers to the habituation period prior to the start of the retrieval trials. The “trials” stage refers to the period of retrieval trials (10 x 1 pup removed from nest) over which no active retrieval behavior took place. The “retrieval” stage refers to periods of active retrieval behavior. i-k) Mobility and social behavioral assessment in experimental animals at adolescence. i) Distance traveled over a 30-minute open field task (*n* = 47-48/group). j) Time spent in each of the three chambers during the social interaction trial (ANOVA, ELS x chamber interaction, *P* = 0.0493, *n* = 28-32). k) Time spent in each of the two interaction zones during the social interaction trial (ANOVA, ELS x zone interaction, *p* = 0.0252, *n* = 28-32). Data are presented as parts-of-whole or mean ± sem. \**P*<0.05, \*\**P*<0.01, \*\*\*\**P*<0.0001. Statistical analyses include the likelihood ratio test (d), unpaired t-test (e, g, i), Mann Whitney test (f), and two-way RM ANOVA (h, j, k). If a significant effect of the repeated measure was found, an FDR correction for multiple comparisons was performed to specify which timepoints or measures were significantly different. All statistical analyses of mean *EF* scores were performed on fourth root transformed data. This was done to help stabilize variance and make the data more normally distributed. All data shown are non-transformed. P= postpartum day, ELS = early life stress, RM = repeated measures, ANOVA = analysis of variance, FDR = False Discovery Rate correction for multiple comparisons. Panel a, Created in BioRender. Mitchell, SB. (2026) https://BioRender.com/9nmbevo. Panel b, Created in BioRender. Mitchell, SB. (2026) https://BioRender.com/iqg3cwz.

First, animals without any maternal experience were assessed and, as expected, did not reliably retrieve pups back to the nest (Fig 5c). They also showed an unanticipated high rate of aggression towards pups (Fig. 5d), which was significantly higher for animals with prior ELS-exposure. Because these pup-naïve animals did not retrieve and often showed aggression to pups, only baseline neural activity was assessed at this preconception timepoint (Fig. 5e). We decided to evaluate *EF*1 in this assay, as it is the stress-vulnerable predictive network having prior association to aversion and negative affect response^15^. During the baseline period prior to the pup retrieval screen, pup-naïve animals did not display a significant difference in *EF*1 score based on prior ELS-exposure (Fig. 5e). However, at P1, ELS-exposed dams showed higher *EF*1 strength when less time was spent on the nest (Supplemental Fig. S29). This suggests greater negative affect network activity when spending less time with pups in early postpartum (Supplemental Fig. S29).

A subsequent pup retrieval assay was performed with postpartum dams at P4 in the homecage. Equipped with maternal experience, no animals demonstrated pup-directed aggression throughout the course of the task, and ELS-exposure did not affect successful pup retrieval rate (Fig. 5f) or time to retrieve (Fig. 5g). Yet, ELS-exposure increased *EF1* strength during all task stages (Fig 5h, Tables S38). There was also a significant effect of task stage on *EF*1, with *EF*1 continually increasing from baseline to active retrieval in the homecage. ELS-exposure did not affect retrieval performance or *EF*1 strength during an additional open field pup retrieval assay at P4 (Supplemental Fig. S30). During pup retrieval, we also evaluated *EF*6, a network with strength that differed substantially between “on- and off-nest” behavior. *EF*6 strength increased across pup retrieval task stages during the homecage retrieval assay, but was unaffected by ELS and during the open field retrieval assay (Supplemental Fig. S31, Table S39). Despite their relevance to the maternal brain, the strength of *EF_on-nest_* and *EF_stage_* was not affected by pup-retrieval, pup-licking, nor dam self-grooming behaviors. Similarly, the strength of these *EFs* during pup-licking or self-grooming was not affected by ELS-exposure (Supplemental Fig. S32).

To better understand the global impact of ELS on brain and behavior, social behavior of experimental animals and same-sex siblings was also assessed prior to implantation and recording, during adolescence. ELS-exposure did not affect locomotor activity (Fig. 5i). However, on the three-chamber social approach task, ELS-exposed mice spent more time in the chamber with a novel mouse than a nonsocial chamber (Fig. 5j) and more time in close social approach (Fig. 5k). While these animals had elevated social approach in adolescence, this did not align with observations of pup-directed maternal behavior in postpartum, indicating there may be brain network activity that stabilizes offspring care behaviors as a part of the maternal transition.

### A task of negative affect before and after the maternal transition

The stress-relevant *EFs* were originally trained and characterized in male C57BL/6 mice using a task designed to induce a negative affect response^15^, the Forced Interaction Test (FIT; Fig. 6b), based loosely on human tasks evoking emotional responses to negatively valenced faces. Since then, *EF*1 has been further validated in both male and female C57s^41^ but no stress *EF*s had been characterized in CD1s nor, for the most part, in female animals. The interesting relationship between stress networks and the maternal experience led us to more deeply investigate the relationship between negative affect and these networks in the context of CD1 female mice. To better characterize these *EF*s in female CD1s, we conducted FITs at both preconception and late postpartum/post-weaning timepoints (Fig. 6a-b)^15^. For this analysis, we evaluated the two stress networks that had revealed baseline differences between stress-resilient and stress-susceptible animals during the FIT in prior work, *EF1* and *EF2*^15^.

**Figure 6:**
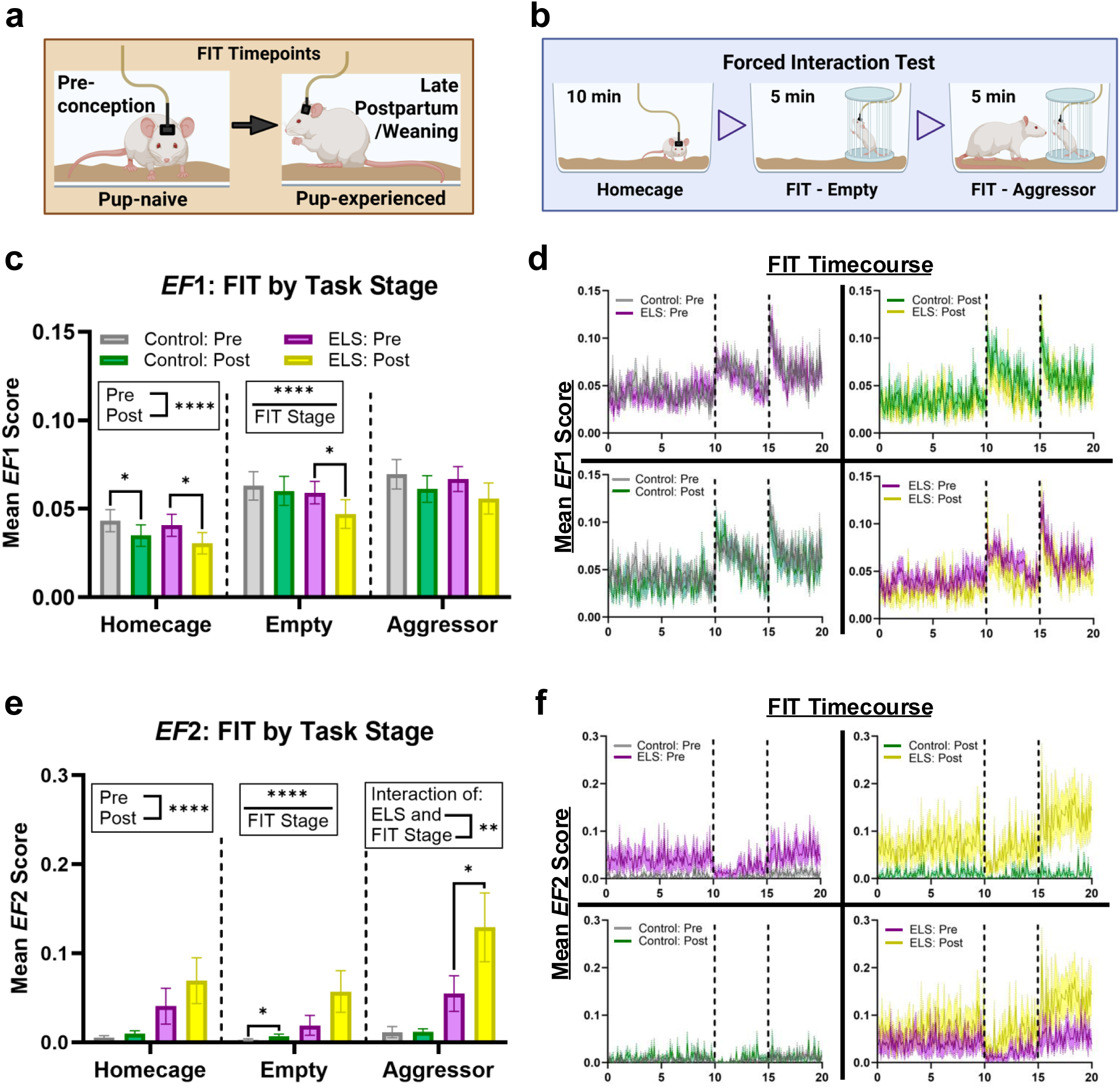
ELS-specific reactivity to negative affect is unmasked following maternal experience in mice. a, Schematic highlighting the two timepoints the FIT was performed in implanted experimental mice. b, Schematic illustration of the three stages of the Forced Interaction Test (FIT). The task is 20-mins in total. c and e, Summary data for all stages of the FIT at both preconception and postpartum timepoints for both control and ELS-exposed dams for c, *EF*1: (REML, FIT stage effect, *P =* 9.17E-14; timepoint effect, *P* = 7.46E-06, *n*=12-20), and e, *EF*2: (REML, FIT stage effect, *P =* 6.94E-11; timepoint effect, *P =* 1.84E-06, ELS x FIT stage interaction, *P* = 0.008212, *n*=11-20). d, Mean *EF*1 and f, *EF*2 scores across the FIT timecourse for all comparisons of interest. The dotted lines denote changes in the stage of the FIT. Data are mean ± sem.; \**P*<0.05, \*\**P*<0.01, \*\*\*\**P*<0.0001. All statistical analyses of mean *EF* scores were performed on fourth root transformed data. This was done to help stabilize variance and make the data more normally distributed. All data shown are non-transformed.Statistical analyses include the three-way RM Mixed Effects Model [REML](c, e). If a significant effect of the repeated measure was found, a False Discovery Rate (FDR) correction for multiple comparisons was performed to specify which timepoints or measures were significantly different. FIT = forced interaction test, *EF* = *Electome Factor*, Pre = preconception, Post = postpartum, ELS = early life stress, REML = restricted maximum likelihood, RM = repeated measures, FDR = False Discovery Rate correction for multiple comparisons Panel a, Created in BioRender. Mitchell, SB. (2026) https://BioRender.com/cuezcgy. Panel b, Created in BioRender. Mitchell, SB. (2026) https://BioRender.com/xdlfq9t.

At both preconception and late postpartum timepoints (Fig. 6c-d), we identified a significant effect of FIT stage on network activity strength, by which the mean *EF*1 score increased from the homecage, to the empty, and then aggressor task stages (Table S40). This aligns with previous findings of FIT *EF*1 patterns in C57BL/6 mice^15,42^. Although no ELS effect was identified, *EF*1 during the FIT was lower postpartum as compared to preconception (Table S41). *EF*2 also increased over FIT stages (Fig. 6e-f, Table S42) but instead was higher in postpartum than preconception overall (Table S43). Additionally, *EF*2 strength was observed to increase with ELS-exposure and significantly interact with FIT stage, as *EF*2 elevation was greatest in the aggressor stage of the FIT. The increased engagement of *EF*2 during a negative affect task in ELS-exposed dams supports the idea of substantial changes stemming from the maternal transition for stress-relevant brain activity. In addition, this unique response to an aggressor in ELS females suggests *EF2* may have specific relevance to early stress exposure in CD1 mice.

An additional version of this analysis was carried out using imputed data for missing or misplaced regions (Supplemental Fig. S33, Tables S44-S47). Once again, this secondary version of the analysis generated results very similar to the dataset presented above, with the same main effects being identified for both *EF*1 and *EF*2. Together, these findings further support the concept of potential unmasked vulnerability in postpartum for ELS-exposed dams.

## Discussion

The best performing maternal-relevant networks identified in our study were those that involved either all or most of the regions we recorded from, showcasing the importance of taking a brain-wide network approach when investigating the maternal transition. These results, in addition to our analyses of *EF* strength across different contexts (e.g. on/off nest) and timepoints, support the identification of unique, discriminative networks that are independently predictive of either maternal care behavior or maternal stage. We found that dynamic networks specifically discriminative of maternal behavior (*EF_on-nest_* and *EF_lick-groom_*) also had a stage-dependent relationship with maternal experience, identifying specific patterns of change even during gestation, before the onset of pup-directed care (Fig. 2b-c). Interestingly, the trajectory for *EF_on-nest_* strength revealed a similar progression as longitudinal *structural* brain changes observed in murine dams ^8^, with levels returning close to baseline by the time of weaning. In particular, two distinct regions of high consequence for *EF_on-nest,_* Vhipp and AMY, followed this structural timecourse ^8^. These two regions have implications for coordinated maternal behaviors^18^ and goal-directed pup attentiveness^27^. In contrast, the trajectory of *EF_stage_* did not return to baseline levels by weaning. In other words, the network most predictive of whether animals are preconceptual or postpartum (i.e. are parents) was not observed to return to preconception levels of network activity. This finding parallels some human studies, which demonstrate that the transition to parenting can result in prolonged changes to both relevant brain structure^1,2^ and function ^3^, persisting past immediate offspring need and even impacting cognitive capacity throughout life ^55^. Although *EF_stage_* was highly amygdala-dependent, it also involved the IL, a neuroanatomical correlate of the human mPFC, which is one region that demonstrates such lasting maternal change ^56^.

Importantly, the activity of these maternal *EF* networks was robust to disruption by ELS (Supplemental Figs. S17-S20, S32). However, ELS did impact neural activity in our study, with changes appearing specifically in networks trained on social stress-related conditions (Figs. 4-6). Impacts of ELS on the stress *EF*s were influenced both by postpartum day and whether dams were engaged in maternal behaviors on the nest, in addition to the interrelationship between these factors. For example, *EF*2 strength was found to be substantially increased for postpartum dams with prior ELS-exposure both in the homecage (Supplemental Fig. S21, Fig. 4d) and during the FIT (Fig. 6e-f). These findings hint at an apparent ELS-induced unmasking of vulnerability with maternal transition as well as an ELS-specific association of lower *EF*2 with “on-nest” behavior. Moreover, the observed increased variability in *EF*2 under the ELS condition could indicate a susceptible sub-population within this group.

Similarly profound differences in network activity of other stress *EF*s (1,3, 6) were identified regarding ELS and maternal experience (Fig. 4c, e, f). After the transition to parenthood, *EF*1 activity in ELS-exposed dams was explicitly stronger “off-nest” (Fig. 4c), while their *EF*1 reactivity during the FIT at the end of this same period was not different from controls (Fig. 6c-d). Conversely, ELS also impacts *EF*6 activity during “on” and “off” the nest behavior, but not in a linear way (*EF*6 is always higher on-nest than off nest, but at P3, EF6 is higher for controls and at P8, EF6 is higher for the ELS group; Fig. 4f, Table S24, Supplemental Fig. S24l, Table S34). ELS-exposed animals also showed heightened *EF*1, but not *EF*6, activity during pup retrieval (Fig. 5h, Supplemental Figure S31). This is expected to be a more stressful context of maternal care, both because it is cognitively and physically demanding^46^, and because having the pups out of the nest represents a degree of threat.

The stress *EF*s, originally developed in C57BL/6 males, had not been previously tested in the CD1 mouse strain^15, 42^. We found that *EF*1 behaves similarly during the FIT in female CD1s, demonstrating an increase in *EF*1 activity (albeit with different dynamics than C57BL/6 male and female mice) with each stage of increasing negative affect. This finding is an important cross-strain validation. Similarly, the substantial increase in *EF*2 activity (Fig. 4d and 6e) observed with our combination ELS strategy also provides an important validation of these prior stress *EF*s for another stress condition across strain. While the “replication” of stress-relevant network activity changes with different strains, stressors, and sex could be interpreted as incremental progress, this work adds important new dimensions to our understanding of these networks. Specifically, here we link these networks to a substantially different type of developmental stress and a conserved mammalian behavioral transition, a frequent and widely applicable mammalian experience (parenthood). This suggests that these networks may have a more universal relevance under different contexts (much like the default mode network has in human fMRI literature). At the same time, the demonstration that maternal experience and behavior alter these neural underpinnings (Fig. 4; *EF*1,2,3,6) of unrelated behavioral states sheds light on how maternal care arises.^15,42^

Additionally, the maternal transition itself also altered the activity of these stress-relevant networks (Fig. 3c, i; Supplemental Fig. S21), and their activity was distinct with maternal care behavioral epochs (Fig. 4). How these neural signatures relate to this distinct, dynamic form of stress is therefore complex. *EF*1 and *EF*6 display remarkably similar but not identical, activity in different contexts: both *EF*1 and *EF*6 go down over time with maternal stage (Fig. 3), both increase across the phases of pup retrieval (Fig. 5, Supplemental Fig. S31), and yet *EF*1 and *EF*6 are again divergent in how they respond to being “on” or “off” the nest (Fig. 4). Curiously, network strength of *EF*1 and *EF*3 displayed divergent patterns over maternal stage (Fig. 3), with a general decrease in strength observed for *EF*1 and no change in *EF3* (controls), while both displayed stronger activity when dams were off the nest (Fig. 4). Given our substantial findings with regard to a role for stress networks in maternal behavior and experience within the context of this study, the peripartal period could be viewed as one of substantial and multifaceted stress. Future work will expand upon these findings by incorporating additional brain regions of known maternal significance while moving toward animal models with greater translational impact.

Our identification of unique networks that change with maternal experience and of those that are resistant to change in the face of stress suggests maternal-specific neural activity that is alterable and specific networks that are essential to species survival. It may be that multiple networks arise simultaneously with differing functions and timecourses to serve multiple adaptive purposes. This overall picture of multiple networks coming on and offline in differing contexts and life stages suggests the possibility of leveraging such network information to inform strategies for preserving the most essential network activity of parenting while modifying others. Such approaches may be desirable to mitigate past stress effects or improve current functioning.

## Acknowledgements

We would like to thank our colleagues Brian Trainor, Catherine Marler, and Bianca Jones-Marlin [insert all who rise to this from our author acknowledgements list] for helpful discussions and feedback. We would like to thank Radha Velamuri and Kayla’hania Williams for technical support. We also acknowledge Biorender, which was used for making figures. We would like to thank our funding agencies who supported this work: NIH Director’s New Innovator 1DP2MH126377-01 (RH), Roy J. Carver Charitable Trust (RH, HES), NINDS T32NS007124 (SBM), Transformative Research Award R01OD039332 (RH and HES).

