## Supplement for "Coordinated Brain-Network Dynamics of Maternal Adaptation"

[Table S2 – *EF_on-nest_* scores by “off-nest” and “on-nest” behavior across postpartum observation days [Controls]	57](#_Toc240022382)

[Table S3 – *EF_on-nest_* scores by on-nest behavior and “on-nest” subtyped behavior (nursing) at P3 [Controls]	57](#_Toc240022383)

[Table S4 – *EF_on-nest_* percent change from baseline: Multiple comparisons by maternal timepoint [Controls]	58](#_Toc240022384)

[Table S5 – *EF_lick-groom_* percent change from baseline: Multiple comparisons by maternal timepoint [Controls]	58](#_Toc240022385)

[Table S6 – *EF_stage_* percent change from baseline: Multiple comparisons by maternal timepoint [Controls]	59](#_Toc240022386)

[Table S7 – *EF_stage_* scores by “off-nest” and “on-nest” behavior across postpartum observation days [Controls]	60](#_Toc240022387)

[Table S8 – Homecage *EF*1 scores: Multiple comparisons by maternal timepoint [Controls]	60](#_Toc240022388)

[Table S9 – Homecage *EF*6 scores: Multiple comparisons by maternal timepoint [Controls]	60](#_Toc240022389)

[Table S10 – *EF_on-nest_* scores by “off-nest” and “on-nest” behavior across postpartum observation days [ELS]	60](#_Toc240022390)

[Table S14 – *EF_stage_* scores by on-nest behavior: Multiple comparisons by postpartum observation day [ELS]	63](#_Toc240022394)

[Table S27 – *EF*1 Maternal homecage: Multiple comparisons by maternal timepoint [Imputed]	67](#_Toc240022407)

[Table S28 – *EF*1 maternal homecage observations: Multiple comparisons by postpartum day [Imputed]	68](#_Toc240022408)

[Table S29 – *EF*1 maternal homecage observations: Multiple comparisons by “off-nest” and “on-nest” behavior [Imputed]	68](#_Toc240022409)

[Table S30 – *EF*2 maternal homecage observations: Multiple comparisons by postpartum day [Imputed]	68](#_Toc240022410)

[Table S31 – *EF*3 Maternal homecage: Multiple comparisons by maternal timepoint [Imputed]	69](#_Toc240022411)

[Table S32 – *EF*3 maternal homecage observations: Multiple comparisons by postpartum day [Imputed]	69](#_Toc240022412)

[Table S33 – *EF*6 maternal homecage: Multiple comparisons by maternal timepoint [Imputed]	69](#_Toc240022413)

[Table S34 – *EF*6 maternal homecage observations: Multiple comparisons by postpartum day [Imputed]	69](#_Toc240022414)

[Table S35 – *EF*6 maternal homecage observations: Multiple comparisons by “off-nest” and “on-nest” behavior [Imputed]	70](#_Toc240022415)

[Table S40 – *EF*1 FIT: Multiple comparisons by FIT stage [Cleaned and Complete]	71](#_Toc240022420)

[Table S41 – *EF*1 FIT: Multiple comparisons by maternal timepoint [Cleaned and Complete]	71](#_Toc240022421)

[Table S42 – *EF*2 FIT: Multiple comparisons by FIT stage [Cleaned and Complete]	71](#_Toc240022422)

[Table S43 – *EF*2 FIT: Multiple comparisons by maternal timepoint [Cleaned and Complete]	72](#_Toc240022423)

[Table S44 – *EF*1 FIT: Multiple comparisons by FIT stage [Imputed]	72](#_Toc240022424)

[Table S45 – *EF*1 FIT: Multiple comparisons by maternal timepoint [Imputed]	72](#_Toc240022425)

[Table S46 – *EF*2 FIT: Multiple comparisons by FIT stage [Imputed]	72](#_Toc240022426)

[Table S47 – *EF*2 FIT: Multiple comparisons by maternal timepoint [Imputed]	73](#_Toc240022427)

### Supplemental Experimental Methods

**Model development detailed overview (SAE-NMF)**

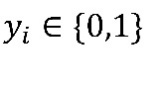
We modeled *Electome Factors* (*EF*s) of maternal behavior or stages using a supervised autoencoder (SAE) with a Non-negative Matrix Factorization (NMF) decoder ^1^. For each time window $i$, let $x_{i}$ denote the spectral feature vector and the corresponding behavioral label (task-dependent; e.g., on-nest vs off-nest). The encoder $f_{\theta}(\cdot)$ maps input features to $K$ latent factors $s_{i}=f_{\theta}\left( x_{i} \right)\in\mathbb{R}_{+}^{K}$, which we refer to as *EF* scores, where $s_{ik}$ quantifies the expression strength of factor $k$ in window $i$. These scores are used in two ways. First, they are decoded back to the feature space via an NMF decoder $\hat{x}_{i}=g_{W}\left( s_{i} \right)=s_{i}W$, where each factor’s loading vector $w_{k}$ is normalized to have fixed $\mathcal{l}_{2}$ norm $\left\| w_{k} \right\|_{2}=\sqrt{p}$ for identifiability and biological justification. Second, supervised prediction is performed using logistic regression on a designated subset of the latent factor scores, producing a predicted probability $\hat{y}_{i}$. The encoder was implemented as a single-hidden-layer feedforward network with 64 units (for prespecified frequency bands) or 128 units (for 1-Hz frequency steps), batch normalization, leaky ReLU activation, and softplus output. Model parameters are learned by minimizing an MSE reconstruction loss and a prediction loss (binary cross-entropy) weighted by $\mu$ :

$$\mathcal{L=}\mathcal{L}_{\text{rec }}\left( x_{i},\hat{x}_{i} \right)+\mu\mathcal{L}_{\text{pred }}\left( y_{i},\hat{y}_{i} \right),$$

where supervision weight $\mu$ controls the trade-off between reconstruction fidelity and task relevance.

To obtain *EF*s tied to distinct maternal-relevant states while maintaining interpretability, we trained four separate SAE-NMF models. Three models were trained by behavioral discriminator (“on” vs. “off-nest”, pup licking vs. all other behavior on the nest, pup licking vs. self-grooming on the nest), and one model was trained by maternal stage (preconception vs. early postpartum). Each model used $10$ factors for feature reconstruction, with $1$ of them designated for prediction of the task/stage-specific label. This design avoids identifiability issues when labels are nested or overlapping (e.g., licking occurs within on-nest windows), ensuring each predictive *EF* corresponds to a single well-defined discriminator. All models were trained using control animals only, with Early Life Stress (ELS) animals held out as an external test set.

**Training, validation, and calculation of *Electome Factors***

Models were optimized using Stochastic Gradient Descent (SGD: learning rate 1 x 10^-3^ or 2 x 10^-3^ (task-dependent), momentum 0.9) with mini-batches sampled to balance animal contributions by inversely weighting each animal by its number of windows. To pretrain the model, decoder weights were first set using an unsupervised NMF decomposition; the resulting latent components were ranked by their discriminative ability for the behavioral label, and the leading component was oriented to have a positive correlation with the label. With the decoder fixed at these NMF-derived weights, the encoder was pretrained to reproduce the corresponding NMF latent activations. The supervision weight $\mu$ (Table S48) was selected based on the relative magnitudes of the reconstruction and prediction losses after pretraining and adjusted, if necessary, so that neither loss exceeded the other by more than a factor of 2.

We employed a simplified nested leave-one-animal-out cross-validation approach to maximize training data utilization in our small sample setting. First, leave-one-animal-out cross-validation was performed with further internal splits: for each fold, training data were split 9:1 into training and validation sets, models were trained with early stopping (patience = 20 epochs, minimum validation loss improvement = 10^-4^), and the best epoch from each fold was recorded. A fixed training duration was then determined based on these best-epoch values and inspection of the validation loss curves. Using this fixed duration, we re-ran leave-one-animal-out cross-validation with all training data in each fold, testing each held-out animal once. This yielded fold-wise independent test AUCs, which were summarized as mean ± sem. Per animal AUCs were compared with chance (0.5) using one-sided exact Wilcoxon signed-rank tests. Finally, a model was trained on all animals using the selected duration to obtain the complete model.

After training, the encoder and decoder weights were frozen and used for projection to independent data. For any new recordings, window-level *EF* scores were obtained directly via the encoder forward pass, $s=f_{\hat{\theta}}(x)$, without additional optimization.

**Feature Thresholding**

To facilitate biological interpretation of each *EF*, we derived a core set of elements capturing the largest contributions to the *EF* loading vector. For a given *EF*, we ranked elements by squared loading magnitude and computed the cumulative proportion of total squared loadings. Elements were selected sequentially from largest to smallest until the accumulated sum exceeded a scree-based threshold chosen to ensure all the most important information was retained (70% threshold for the main analysis, see Supplemental Figs. S3-4, S6, S9, S13-14 for other thresholds). Elements within this core set may still differ in how specific they are to the *EF*, since an element may carry comparable weight in the remaining factors, which serve only to reconstruct the spectral activities. As a complementary annotation, we therefore computed a relative specificity score for each selected element: the supervised-factor loading divided by the sum of loadings across all factors. We report this as an annotation of supervised *EF* specificity (highlighted above 0.3; see Figures 1f, 2e, and Supplemental Figs. S5, S7, S10, S15).

**Determination of required numbers of brain regions for Stress *EF* projections**

When *EF*s defining stress-related brain states were originally trained ^2^, it was discovered (likely due to the generative nature of these *EF*s) that at least two brain regions could be removed and still retain the same overall network activity. However, all this work was done in male C57 animals and only for *EF*s 1-3. We tested (in CD1 female animals) the impact of missing regional LFP data on the calculation of network strength of all stress-associated *EF* network scores. A subset of experimental animals that had all regions correctly targeted were used for this test and specifically, the preconception FIT recording was used. We sequentially removed regions from each dataset and re-calculated *EF* strength scores for all combinations of missing regions, starting with removing one region at a time and moving up to three regions total, against the results calculated using all regions. Comparisons were performed by assessing mean timecourse data (see Supplemental Results and Supplemental Fig. S22).

**Imputation of missing LFP regional data**

Imputation was employed to better utilize LFP data from animals with mistargeted (excluded) implant regions. We modeled the 808-dimensional vector of retained Fourier coefficients (stacked across regions and frequencies) within each subset as complex multivariate normal. Specifically, if $x_{mt}$ denotes the coefficients for time window $t$ from mouse $m$ from a particular group, then we assume that $x_{mt}$ follows the multivariate complex normal distribution with mean $\mu_{t}$ and covariance $\Sigma_{t}$, where we impose a low-rank factor model structure (i.e., low-rank-plus-diagonal matrix) on $\Sigma_{t}$ . The groups consisted of two (mouse group: control and ELS), four (mouse group: control and ELS; behavior: “on” and “off-nest”) or six factor combinations (mouse group: control and ELS; FIT stage: homecage, FIT-empty, and FIT-aggressor). For each group, we estimated the two parameters within each time window using data from 10 mice with complete recordings and a rank-100 low-rank structure (i.e., $m=1, \ldots, 10$; the $t$ range varies by group; and the number of latent factors $=$ 100). We then used these estimates to impute the group-, behavior-, and/or experimental stage-specific missing recordings, taking the conditional mean of the missing recordings given the observed recordings from the mice with complete data. Specifics of the empirical evaluation are provided below. See Hultman, I. et al. (2026, arXiv)^3^ for details of the estimation algorithm.

Imputation was developed and tested as a method for making the most use of animals with regions mistargeted (i.e. missing) that impeded their calculation of *EF* network strength. We performed the same LFP preprocessing steps here as in the training models and reiterate them. For each of the nine datasets (obtained across the maternal timecourse and detailed below), we segmented the LFP recordings into non-overlapping 1-second windows. Because the signals were sampled at 1000 Hz, each window contained 1000 time points. For each window and each of the eight brain regions, we computed the Fourier transform using the fast Fourier transform (FFT) and retained only the coefficients corresponding to 0-100 Hz. Given the sampling rate and window length, the frequency resolution was 1 Hz, yielding 101 Fourier coefficients per brain region per window. These coefficients served as the features in our model.

After windowing the data and extracting Fourier coefficients for each 1-second segment, we assigned labels to each time window. First, every window was labeled by group as either control or ELS. In addition, we assigned maternal stage-specific, timecourse-dependent labels based on the recording context. For maternal homecage recordings, all windows were labeled as a single stage (homecage). For FIT recordings, windows were labeled as belonging to one of three task stages: homecage, FIT 1 (empty), or FIT 2 (aggressor). For the recordings involving maternal care observations, windows were additionally labeled according to periods of behavior location, either “on-nest” or “off-nest”.

Initially, our imputation strategy was tested in animals with all brain regions successfully targeted as described in Hultman, I. et al.^3^. Using ten animals that had all 8 regions correctly targeted gave us 360 = 10 × (8+28) tests corresponding to the eight regions and 28 pairs of regions for each of the ten “good” mice. To see how close our predictions were to the actual data, we inverse Fourier transformed the imputed FFT coefficients to obtain imputed LFP data in the time domain and computed the log relative norm error:

$$\text{logerr}_{\left( m,r \right)}=\log\frac{\left| \text{X}_{\text{true}\left( m,r \right)}-\text{X}_{\text{pred}\left( m,r \right)} \right|}{\left| \text{X}_{\text{true}\left( m,r \right)} \right|},$$

where *m* = 1*,...,*10 indexes the mouse, *r* = 1*,...,*36 indexes either the single region or region pair that was left out and imputed, **X***_true_*_(_*_m,r_*_)_ denotes the actual observed LFP data for the *m*th mouse at the *r*th region or pair of regions, and **X***_pred_*_(_*_m,r_*_)_ denotes the imputed LFP data for the *m*th mouse at the *r*th region or pair of regions.

We used three methods to impute the missing data. For each method, we grouped the data based on whether the mice were control or ELS, and to which of the three task stages the data corresponded: homecage, FIT-empty, or FIT-aggressor. This resulted in the data being split into six subsets. For two of the imputation methods, we modeled the vector of 101 Fourier coefficients for each of the eight brain regions pertaining to each time window as coming from the same subset-specific complex multivariate normal distribution, i.e.:

**x**(*s,t,m*) ∼ $\mathbb{C}N$*p* (*µs,***Σ***s*)*,*	(1)

where *s* = 1*,...,*6 indexes the subset of the data based on experimental group to which each mouse belonged (ELS or control) and stage of the FIT task, *t* = 1*,...,T_s_* indexes the time window of the *s*th subset of data for the *m*th mouse, *T_s_* denotes the total number of time windows in the *s*th subset, *p* = 808 = 101×8 for the 101 Fourier coefficients for each of the eight brain regions, *µ_s_* ^∈^ C*^p^* denotes the mean parameter feature vector for the *s*th subset, and **Σ***_s_* ∈ C*^p^*^×^*^p^* denotes the complex covariance matrix between the 808 features for the *s*th subset. For each subset, we estimated *µ_s_*, denoted as *µ*ˆ*_s_*, by just computing the mean feature vector with NA removal across all time windows for all mice in the *s*th subset.

For the first imputation method we used our SSFA model which assumes the following structure on the covariance matrix from (1):

$\boldsymbol{\Sigma}_{s}=\boldsymbol{\Lambda}_{\mathbf{s}}{\boldsymbol{\Lambda}_{\boldsymbol{s}}}^{\boldsymbol{\star}}+\boldsymbol{\Psi}_{\mathbf{s}}$,	(2)

where **Λ***_s_* ∈ C*^p^*^×^*^k^* denotes the matrix of factor loadings and **Ψ***_s_* ∈ R*^p^*^×^*^p^* denotes the diagonal feature noise matrix. We set the number of factors to *k* = 100 for each subset of the data and we imposed no sparsity on the factors. We subtracted ${\hat{\boldsymbol{\mu}}}_{s}$ from each observation **x**_(_*_s,t,m_*_)_ and used the de-meaned dataset to estimate **Λ***_s_* and **Ψ***_s_*. While we could use all the mice pertaining to the *s*th subset to obtain ${\hat{\boldsymbol{\mu}}}_{s}$ via NA removal based on the elements of the mice data that had NA values where their electrodes were found to have been incorrectly implanted, we currently can only use the mice with complete data in our model when estimating (2). This means that for the three subsets involving the eight control mice, for each mouse held out in order to create one of the test sets, we only used the seven remaining “good” control mice with complete data for estimating (2), and for the three subsets involving the two ELS mice, for each mouse held out in order to create one of the test sets, we only used the single other remaining “good” ELS mouse for estimating (2).

For the second imputation method we used the traditional complex-variate covariance computation with NA removal to estimate **Σ***_s_* from (1). Let *x*˜_(_*_i,s,t,m_*_)_ = *x*_(_*_i,s,t,m_*_)_ − ${\hat{\boldsymbol{\mu}}}_{(i,s)}$ denote the de-meaned observation pertaining to the *i*th feature for the *m*th mouse at the *t*th time window in the *s*th subset of the data, then we estimated the covariance between the *i*th and *j*th features in the *s*th subset as:

$\hat{\sigma}_{\left( i, j, s \right)}=\frac{1}{a_{(i,j,s)}}\sum_{m=1}^{M_{s}} \sum_{t=1}^{T_{s}} \tilde{x}_{(i,s,t,m)}{\overline{\tilde{x}}}_{(j,s,t,m)}\boldsymbol{1}_{(i,j,m)}$,

where ${\overline{\tilde{x}}}_{(j,s,t,m)}$ denotes the complex conjugate of $\tilde{x}_{(i,s,t,m)}S$, 1(*i,j,m*) denotes the indicator that neither *i*th nor *j*th features of the *m*th mouse’s data are set to NA, *M_s_* denotes the total number of mice in the *s*th subset, and $a_{(i,j,s)}=\sum_{m=1}^{M_{s}} \sum_{t=1}^{T_{s}} \boldsymbol{1}_{(i,j,m)}$ denotes the total number of non-NA pairs ${(\tilde{x}}_{(i,s,t,m)},\tilde{x}_{(j,s,t,m)})$. We set the (*i,j*)th element of ${\hat{\boldsymbol{\Sigma}}}_{s}$ to $\hat{\sigma}_{\left( i, j, s \right)}$.

Once we estimated ${\hat{\boldsymbol{\Sigma}}}_{s}$ for *s* = 1*,...,*6 for the first two imputation methods, we used it to predict the missing values for each mouse’s data in the test sets. Let **x**_(mis_*_,s,t,m_*_)_ and **x**_(obs_*_,s,t,m_*_)_ denote the vector of features with NA values and the vector of observed non-NA features respectively for the *m*th mouse at the *t*th time window in the *s*th subset. We assume these vectors are from the following complex multivariate normal distribution, which is just a re-expression of (1) above:

$\left[ \begin{matrix} \mathbf{x}_{(\mathrm{mis},s,t,m)} \\ \mathbf{x}_{(\mathrm{obs},s,t,m)} \end{matrix} \right]\mathbb{\sim C}N\left( \left[ \begin{matrix} \boldsymbol{\mu}_{(\mathrm{mis},s)} \\ \boldsymbol{\mu}_{(\mathrm{obs},s)} \end{matrix} \right],\left[ \begin{matrix} \boldsymbol{\Sigma}_{(\mathrm{mis},s)} & \boldsymbol{\Sigma}_{(\mathrm{cov},s)} \\ \boldsymbol{\Sigma}_{(\mathrm{cov},s)}^{\star} & \boldsymbol{\Sigma}_{(\mathrm{obs},s)} \end{matrix} \right] \right)$,

where *µ*_(mis_*_,s_*_)_ and *µ*_(obs_*_,s_*_)_ denote the mean vectors for the missing and observed features respectively in the *s*th subset, **Σ**_(mis_*_,s_*_)_ and **Σ**_(obs_*_,s_*_)_ denote the covariance matrices for the missing and observed features respectively in the *s*th subset, and **Σ**_(cov_*_,s_*_)_ denotes the matrix of covariance terms between the elements of the missing features and the elements of the observed features. We split and reordered our estimates ${\hat{\boldsymbol{\mu}}}_{s}$ and ${\hat{\boldsymbol{\Sigma}}}_{s}$ into sets $\left( {\hat{\boldsymbol{\mu}}}_{(\text{mis, s})},{\hat{\boldsymbol{\mu}}}_{(\text{obs, s})} \right)$ and $\left( {\hat{\boldsymbol{\Sigma}}}_{(\text{mis, s})},{\hat{\boldsymbol{\Sigma}}}_{(\text{obs, s})},{\hat{\boldsymbol{\Sigma}}}_{(\text{cov, s})} \right)$ which we then used to predict **x**_(mis_*_,s,t,m_*_)_ by setting its values to the conditional expectation:

${\hat{\boldsymbol{x}}}_{(\mathrm{mis},s,t,m)}={\hat{\boldsymbol{\mu}}}_{(\text{mis, s})}+{\hat{\boldsymbol{\Sigma}}}_{(\text{cov, s})}{\hat{\boldsymbol{\Sigma}}}_{(\text{obs, s})}^{-1}(\mathbf{x}_{(\mathrm{obs},s,t,m)}-{\hat{\boldsymbol{\mu}}}_{\left( \text{obs, s} \right)})$.

For the third imputation method we simply set the missing values for the *i*th feature of every observation to be the NA-removed mean estimated across all mice in the *s*th subset, i.e. we set *x*_(_*_i,s,t,m_*_)_ = ${\hat{\boldsymbol{\mu}}}_{(\text{i,s})}$ for every time window *t* = 1*,...,T_s_* for every mouse *m* = 1*,...,M_s_* that had missing values at the *i*th feature.

Our empirical comparisons show that SSFA-based imputation outperforms both covariance-based and mean imputation. We therefore use SSFA with $k=100$ for the imputation described below. Detailed comparisons in a similar setting are provided in Hultman, I. et al.^3^

**Using imputations on *EF* data**

For each animal with any brain region identified via histology as incorrectly implanted, we removed the incorrect LFP data and set the corresponding Fourier coefficients to NA (“missing” or “not available”) and then imputed them using our model. Imputation was performed within strata defined by shared labeling information, so that windows were only imputed using other windows recorded under the same experimental conditions. For example, in an experiment involving labeled observations of maternal care, each time window carried (i) a group label (control vs ELS) and (ii) a behavior label (“on-nest” vs. “off-nest”). This yields four strata: control/”off-nest”, control/”on-nest”, ELS/”off-nest”, and ELS/”on-nest”. We fit the SSFA model separately within each stratum, producing a condition-specific model that we then used to impute NA values for windows belonging to that same stratum.

Below we summarize the sample sizes for each experimental timepoint, along with the corresponding labeling structure (group, task stage, maternal timepoint, behavior) and the number of mice with at least one brain region requiring imputation:

- **Preconception-FIT:** We analyzed 41 mice (20 control, 21 ELS). Among the control mice, 10 required imputation in one or more brain regions; among the ELS mice, 13 required imputation. Recordings spanned three task stages: homecage, FIT-empty, and FIT-aggressor.
- **Preconception Homecage:** We analyzed 41 mice (20 control, 21 ELS). Among the control mice, 10 had one or more brain regions imputed; among the ELS mice, 13 required imputation. Homecage recordings comprised a single maternal timepoint (preconception).
- **Gestational Homecage:** We analyzed 34 mice (18 control, 16 ELS). Among the control mice, 8 required imputation in one or more brain regions; among the ELS mice, 10 required imputation. Homecage recordings comprised a single maternal timepoint (E17/gestation).
- **P1:** We analyzed 22 mice (11 control, 11 ELS). Among the control mice, 5 required imputation in one or more brain regions; among the ELS mice, 6 required imputation. Maternal homecage recordings were divided into periods of “on” or “off-nest” behavior.
- **P3:** We analyzed 32 mice (17 control, 15 ELS). Among the control mice, 7 required imputation in one or more brain regions; among the ELS mice, 9 required imputation. Maternal homecage recordings were divided into periods of “on” or “off-nest” behavior.
- **P8:** We analyzed 32 mice (15 control, 17 ELS). Among the control mice, 6 required imputation in one or more brain regions; among the ELS mice, 11 required imputation. Maternal homecage recordings were divided into periods of “on” or “off-nest” behavior.
- **P14:** We analyzed 28 mice (13 control, 15 ELS). Among the control mice, 4 required imputation in one or more brain regions; among the ELS mice, 10 required imputation. Maternal homecage recordings were divided into periods of “on” or “off-nest” behavior.
- **P20 Homecage:** We analyzed 28 mice (13 control, 15 ELS). Among the control mice, 4 required imputation in one or more brain regions; among the ELS mice, 10 required imputation. Homecage recordings comprised a single maternal timepoint (late postpartum).
- **Postpartum-FIT:** We analyzed 25 mice (12 control, 13 ELS). Among the control mice, 4 required imputation in one or more brain regions; among the ELS mice, 11 required imputation. Recordings spanned three task stages: homecage, FIT-empty, and FIT-aggressor.

We describe the SSFA method again for imputation. To impute missing values, we treated each 1-second window as an 8-block complex-valued feature vector, with 101 Fourier coefficients per brain region. Within each subset defined by the relevant labels (for example, control/ELS and task stage or “off-nest” and “on-nest” behavior), we modeled these window-level vectors as independent draws from a subset-specific complex multivariate normal distribution; that is,

$\boldsymbol{x}_{\left( s,t,m \right)}\mathbb{\sim C}N_{p}\left( \boldsymbol{\mu}_{s},\boldsymbol{\Sigma}_{s} \right),$	(1)

where $s=1,\ldots,S_{e}$ indexes the subset of the data based on experimental group (ELS/control) and stage/behavioral label of the experiment, $S_{e}$ denotes the total number of subsets in the *e*th experimental setting, $t=1,\ldots,T_{s\left( m \right)}$ indexes the time window of the *s*th subset of data for the *m*th mouse, $T_{s\left( m \right)}$ denotes the total number of time windows in the *s*th subset for the *m*th mouse, $p=808=101\times8$ for the 101 Fourier coefficients for each of the eight brain regions, $\mu_{s}\in\mathbb{C}^{p}$ denotes the mean parameter feature vector for the *s*th subset of the *e*th experimental setting, and $\Sigma_{s}\in\mathbb{C}^{p\times p}$ denotes the complex covariance matrix between the 808 features for the *s*th subset of the *e*th experimental setting. For each subset, we estimated $\mu_{s}$, denoted as $\hat{\mu_{s}}$, by just computing the mean feature vector with NA removal across all time windows for all mice in the *s*th subset of the *e*th experimental setting.

Our SSFA model assumes the following structure on the covariance matrix from (1):

$\boldsymbol{\Sigma}_{\boldsymbol{s}}=\boldsymbol{\Lambda}_{\boldsymbol{s}}\boldsymbol{\Lambda}_{\boldsymbol{s}}^{\text{*}}+\boldsymbol{\Psi}_{\boldsymbol{s}},$	(2)

where $\boldsymbol{\Lambda}_{\boldsymbol{s}}\in\mathbb{C}^{p\times k}$ denotes the matrix of factor loadings and $\boldsymbol{\Psi}_{\boldsymbol{s}}\in\mathbb{R}^{p\times p}$ denotes the diagonal feature noise matrix. We set the number of factors to $k=100$ for each subset of the data. We subtracted $\hat{\mu_{s}}$ from each observation $x_{\left( s,t,m \right)}$ and used the de-meaned dataset to estimate $\Lambda_{s}$ and $\Psi_{s}$. We currently use mice with complete data, having all regions correctly placed, in our model when estimating (2).

Once we estimated ${\hat{\boldsymbol{\Sigma}}}_{s}$ for $s=1,\ldots,S_{e}$, we used it to predict the missing values for each mouse that had data set to NA based on the histology analysis. Let $x_{\left( \text{mis},s,t,m \right)}$ and $x_{\left( \text{obs},s,t,m \right)}$ denote, respectively, the subvectors of NA and observed features for mouse (m) at time window (t), within subset (s) of experimental setting (e). We assume that the full feature vector follows the complex multivariate normal distribution and express (1) in the block form as

$$\left[ \begin{matrix} x_{\left( mis,s,t,m \right)} \\ x_{\left( obs,s,t,m \right)} \end{matrix} \right]\mathbb{\sim C}N_{p}\left( \left[ \begin{matrix} \boldsymbol{\mu}_{\left( mis,s \right)} \\ \boldsymbol{\mu}_{\left( obs,s \right)} \end{matrix} \right],\left[ \begin{matrix} \boldsymbol{\Sigma}_{\left( mis,s \right)} & \boldsymbol{\Sigma}_{\left( cov,s \right)} \\ \boldsymbol{\Sigma}_{\left( cov,s \right)}^{\text{*}} & \boldsymbol{\Sigma}_{\left( obs,s \right)} \end{matrix} \right] \right),$$

where $\mu_{\left( mis,s \right)}$ and $\mu_{\left( obs,s \right)}$ denote the mean vectors for the missing and observed features respectively in the *s*th subset, $\Sigma_{\left( mis,s \right)}$ and $\Sigma_{\left( obs,s \right)}$ denote the covariance matrices for the missing and observed features respectively in the *s*th subset, and $\Sigma_{\left( cov,s \right)}$ denotes the matrix of covariance terms between the elements of the missing features and the elements of the observed features. We partitioned and reordered our estimates $\hat{\mu_{s}}$ and $\hat{\Sigma_{s}}$ into sets $(\hat{\mu}_{mis,s}, \hat{\mu}_{obs,s})$ and $(\hat{\Sigma}_{mis,s}, \hat{\Sigma}_{obs,s}, \hat{\Sigma}_{cov,s})$. We then imputed $x_{\left( \text{mis},s,t,m \right)}$ by setting it equal to its conditional expectation given the observed entries $x_{\left( \text{obs},s,t,m \right)}$:

$\hat{x}_{(\mathrm{mis},s,t,m)}=\hat{\mu}_{mis,s}+\hat{\Sigma}_{cov,s}{\hat{\Sigma}_{obs,s}}^{-1}\left( x_{\left( \mathrm{obs},s,t,m \right)}-\hat{\mu}_{obs,s} \right)$.

After imputing missing values as needed, we converted the Fourier-domain data back to the time domain using the inverse fast Fourier transform (iFFT). We then used the resulting imputed LFP signals, together with the original (non-imputed) LFP signals, in the backprojection analyses to obtain stress *EF* scores.

**Open field pup retrieval at P4**

Following the completion of the home cage pup retrieval assay (described in the main methods), implanted dams were removed from their home cage and placed in an open field chamber with white, opaque walls (18” long x 18” wide x 16” tall). The dams were left alone in the chamber for a duration of 10 minutes for habituation. Dams were then briefly returned to their home cage while the open field chamber was cleaned (Super Sani Cloth®, PDI Inc., NJ) and dried. Four pups belonging to the dam were chosen at random and placed one per corner in the open field chamber. The dams were then returned to the open field chamber and observed for an additional 10 minutes. At the end of the 10-minutes, if no pups had been grouped together, the dam received a score of zero. If two pups were grouped together, then the dam received a score of one. If two sets of two pups were grouped together, then the dam received a score of three. If all four pups were found grouped together, then the dam received a score of four ^4^. In addition to experimenter observation and scoring of the pup-grouping result, trials were also video recorded, and neurophysiological data were simultaneously gathered throughout the trial (Supplementary Figs. S30-S31, Tables S38-S39).

### Supplemental Experimental Results

**Additional *EF_on-nest_* network validations, including different behavioral contexts**

To ensure that our AUCs were meaningful above chance, we evaluated group AUC values for both the early postpartum (P1, P3) and P8 data sets and found them to be systematically and significantly higher than chance prediction for both control (Wilcoxon *P* = 0.003906) and ELS (Wilcoxon *P* = 0.031250) animals. We also evaluated *EF_on-nest_* network strength with respect to non-nursing sub-typed behaviors (licking/grooming of pups, self-grooming, and pup retrieval) at P3-P4 and found that *EF_on-nest_* was significantly less engaged during licking of pups as compared to all other behaviors (Supplement Fig. S32). *EF_on-nest_* was unaffected by self-grooming and pup retrieval behaviors.

***EF_on-nest-1-Hz_* network not substantially improved over guided bands**

To evaluate whether additional frequencies contribute to maternal brain state, a maternal brain network of “on-nest” behavior was further trained (LOO cross validation) using a more computationally costly approach of 1-Hz frequency steps (Supplemental Fig. S4-5). The “on-nest-1-Hz” *EF* (*EF_on-nest_*_-1-Hz_) resulted in the inclusion of more frequencies while having similar overall performance (leave-one-animal-out cross-validation AUC mean ± sem = 0.72 ± 0.03 , *n* = 8) to *EF_on-nest_* with the early postpartum timepoint data used in training. *EF_on-nest_*_-1-Hz_ had an improved AUC moving into mid-postpartum (P8 test AUC mean ± sem = 0.79 ± 0.03, *n* = 8), suggesting that this network may be more engaged at later maternal timepoints. This trend was also reflected by external testing on ELS animals (P1, P3 AUC mean ± sem = 0.67 ± 0.06, *n* = 6; P8 AUC mean ± sem = 0.74 ± 0.06, *n* = 6). Despite having similar overall features as the network trained with guided frequency bands, a notable difference of *EF_on-nest_*_-1-Hz_ was the contribution of power for all recorded regions within the lower frequencies, which was not seen in *EF_on-nest_*. However, more striking was the shared feature of VHipp coherence with multiple regions being an important network contributor. Overall, this *EF_on-nest_*_-1-Hz_ network was highly consistent with our more efficient guided method. Thus, we continued using the original method for the remainder of our maternal behavioral investigations.

**Additional characterization of the *EF_licking_* network**

Because *EF_on-nest_* discriminates neural activity from when mice are “on” vs. “off” the nest, it discriminates conditions that not only have a difference in maternal condition (pups or no pups) but also in other environmental features such as temperature, texture, scent, etc. The majority of a dam’s time spent on the nest is engaged in nursing behavior, an activity that can span significant lengths of time and include passive pup engagement. Therefore, to look more directly at maternal sub-behavior requiring direct and active social interaction with pups, we trained a model of maternal licking behavior (Supplemental Figs. S6-S7). To examine active maternal behaviors within the same environmental context and thus rule out the possibility that our networks are merely encoding environmental differences, we examined licking behavior solely within the context of the nest. For *EF_licking_* (licking vs non-licking while on-nest), P3 on-nest windows were labeled licking (1) or non-licking (0); cross-validation achieved AUC = 0.78 ± 0.05 (*n* = 8), and external testing on ELS animals yielded AUC = 0.71 ± 0.08 (*n* = 6). Group AUC values were found to be systematically and significantly higher than chance prediction for controls (Wilcoxon *P* = 0.007812) and trending for ELS (Wilcoxon *P* = 0.078125) animals. The 2-7 Hz oscillatory frequency band was observed to be essential, with all regional coherence pairs except those involving VHipp found to have absolute strength contributions to the *EF_licking_* network in this range (Supplemental Fig. S7). Approximately half of the selected features within the 2-7 Hz band were also found to have significant relative specificity with respect to the discriminative *EF_licking_*. These contributing coherence pairs include PrL with BLA, CeA, MeA, and VTA; IL with BLA, MeA, and VTA; VTA with BLA and NAc, and MeA with NAc. Absolute strength contributions were also detected for IL-PrL coherence for both the 8-12 and 14-23 Hz bands, with IL-NAc and BLA-CeA coherence additionally contributing at 14-23 Hz. In particular, prefrontal coherence with amygdala regions seems especially distinctive and strong in this network (Supplemental Fig. S7). Finally, over time, the *EF_licking_* network does not change much from baseline, except at P8. As pups are becoming more active, it becomes less engaged in both control and ELS mice (Supplemental Figs S12, S18, Table S12). By quantifying *EF_licking_* scores by pup-licking and self-grooming behaviors on the nest at P3 (Supplemental Fig. S8), we found that the strength of the network increased with both licking and self-grooming behaviors. We modeled the “lick-groom” network detailed below, with the goal of discriminating between these two active on-nest behaviors.

**Additional characterization of the *EF_lick-groom_* network**

As described in the main text, we trained a model that discriminates times when the mice are licking pups from those when they are licking to groom themselves on the nest (*EF_lick-groom_*). This way, the motor functions and environment are expected to be engaged in the same way with the main difference being pup-directed (1) vs. self-directed behavior (0), reflecting the mutual exclusivity of these behaviors (Supplemental Figs. S9-S10). We describe this approach and analysis more thoroughly here. P3 on-nest windows were labeled; two control animals were not included as they lacked sufficient data for both behaviors at P3 (one had no on-nest self-grooming windows and the other had only two), and cross-validation achieved AUC $=0.64\pm0.04$ ($n=6$), with external testing on ELS animals yielding AUC $=0.64\pm0.08$ ($n=4$). Group AUC values were found to be systematically and significantly higher than chance prediction for controls (Wilcoxon *P* = 0.015625), but not for ELS (Wilcoxon *P* = 0.125000) animals. While this prediction is more modest than our other EFs, the *n* is small and the exclusivity of the 8-12 Hz oscillatory frequency band is worth noting. Absolute strength contributions were found for coherence of MeA with BLA, CeA, IL, PrL, and NAc; CeA with BLA, IL, PrL, and NAc; and BLA with IL, PrL, and NAc. VTA coherence, especially with CeA, seems especially distinctive and strong in this network (Supplemental Fig. S10). Overall, this *EF* network seemed strongest at baseline. After declining during gestation, a partial increase in *EF_lick-groom_* network strength was observed in early postpartum (P1), followed by a gradual decline, returning to approximately gestation levels (Fig. 2c, Supplemental Fig. S19, Tables S5, S13). These fluctuations could be reflective of the shifting balance between offspring and self-care during this time. We quantified *EF_lick-groom_* scores by pup-licking and self-grooming behaviors on the nest at P3 and found that the strength of the network increased with pup-licking (when compared to all other behaviors on the nest; Fig. 1h) and decreased with self-grooming (when compared to pup-licking; Fig. 1i). These results further support the discriminative power of the network for maternal licking behavior.

**Additional *EF_stage_* network validations, including different behavioral contexts**

To ensure that our AUCs were meaningful above chance, we evaluated group AUC values and found them to be systematically and significantly higher than chance prediction for both control (Wilcoxon *P* = 0.003906) and ELS (Wilcoxon *P* = 0.015625) animals. We also evaluated *EF_stage_* network strength with respect to non-nursing sub-typed behaviors (licking/grooming of pups, self-grooming, and pup retrieval) at P3 and found that (similar to *EF_on-nest_*) the *EF_stage_* network was significantly less engaged during licking of pups as compared to all other behaviors (Supplement Fig. S32). *EF_stage_* was likewise unaffected by self-grooming and pup retrieval behaviors.

**Detailed results of *EF_stage-1-Hz_* network**

Like *EF_on-nest_*, a corresponding model was trained using 1-Hz frequency-steps (Supplementary Figs. S14-S15). Cross-validation achieved an AUC of 0.74 ± 0.05 (*n* = 8), and external validation in ELS animals yielded an AUC of 0.71 ± 0.09 (*n* = 6). Overall, the *EF_stage-1-Hz_* model was not as predictive as the guided-frequency band network (*EF_stage_*). There were minor differences and overarching similarities observed in the specific contributing features of the two “maternal stage” networks. Differences again highlighted the contribution of power within regions, this time for VHipp at 2-11Hz and MeA at 3 and 21-55Hz. There was also the striking likeness between the two approaches in the prominence of the BLA-MeA, CeA-MeA, and MeA-NAc coherence pairs. For the 1-Hz frequency iteration, these strong network contributions were found in frequencies above 24, 27, and 33Hz, respectively.

**Assessing the impact of missing regions on stress *EF* scores in animals with on-target electrode placement**

A challenge of implanting multiple brain regions in outbred animals is that the morphological heterogeneity can result in missed targeting, sometimes of a single or multiple regions in a given animal. Quality control measures can also pick up anomalies in the neural data, sometimes requiring regional data to be removed from analysis. Previous work in male C57 mice identified that *EF*1 scores can still be calculated from animals missing up to two regions and provide reliable network scores^2,5^. We sought to evaluate this for up to three missing regions and amongst the other stress *EF*s, for the female CD1 mice presented here. Our test set included experimental animals having perfect electrode targeting. We evaluated averaged network activity strength for all versions of the missing vs. complete preconception FIT data. This experimental condition was chosen, as it exhibits expected, timecourse-specific impacts on the stress *EF* scores, thereby providing landmarks for comparison. When a region or combination of missing regions did not impact the stress *EF* FIT timecourse, the corresponding data was included in the maternal and FIT stress *EF* analyses. We refer to this inclusive and verified dataset as the “cleaned and complete” dataset. See Supplemental Fig. S22 for example traces of averaged timecourse data assessed for missing recording region impact, and Table S26 for timepoint and stress-*EF* specific animal numbers. The “cleaned and complete” dataset was used for data analysis presented in main Figs. 3-6.

**Calculating *EF* scores with missing regions**

In our prior studies using multi-site in vivo neurophysiological approaches in inbred mouse strains, we achieved a >97% rate of correctly targeting electrode placements^2,6^. However, in the CD1 female animals used for this study, post-mortem histological analyses revealed a high rate of morphological diversity that resulted in a high mis-target rate. As such, we implemented several strategies for assessing network strength in the absence of some regions. As an alternative to the “cleaned and complete” datasets, we developed a method for imputing missing data from animals with mostly correct implants with only one to three missing regions. Specifically, our histology revealed that some mice had misplaced implants in specific brain regions, so the corresponding electrophysiology recordings were treated as missing and required either performance validation (see: “Assessing the impact of missing regions on stress EF scores in animals with perfect electrode targeting” above) or imputation (results below) in order make the most of our collected data.

**Proof of imputation success in animals with perfect electrode targeting:**

Prior to implementing an imputation strategy on the data in this paper, we first tested the success of this approach using “perfect” data from animals with all regions correctly targeted. Thus, we used LFP data recorded from seven different brain regions in ten mice that had perfect placement of all regional electrodes, eight of which were control mice and two of which were ELS mice. We created test data by leaving out one region at a time from each of the ten “good” mice and imputing the missing FFT values and then leaving out each pair of regions at a time and imputing the missing FFT values. We tested three different methods for imputing the data (see Supplemental Methods): SSFA, Cov, and Mean. We show the results in Supplemental Fig. S23, where ’SSFA’ identifies the results for our factor model, ’Cov’ identifies the results for the traditional covariance estimate with NA removal, and ’Mean’ identifies the results for the simple mean with NA removal. From all three figure panels, we see that SSFA generally performs the best in terms of log relative prediction error. Panel **c** shows that even in the cases where the traditional covariance method outperformed SSFA, the SSFA results were similar, whereas several of the traditional covariance method’s predictions were significantly worse than the SSFA predictions.

**Use of imputation to analyze stress *EF* data:**

For this analysis, we used local field potential (LFP) recordings from 41 mice across eight brain regions, thus increasing our *n* by up to 23 animals over the dataset in which recordings with missing regions were excluded. The increase in *n* for each experimental condition was more modest when compared to the “cleaned and complete” dataset but theoretically reflected activity from all regions. See Table S26 for animal number comparisons across all conditions. We initially imputed these data for the forced interaction test. Data were collected under nine experimental conditions, with different subsets of the 41 mice contributing to each condition: preconception homecage, pre-FIT, post-FIT, gestational homecage, and P1, P3, P8, P14, and P20 homecage.

**Imputation results for the “Maternal Homecage”, “Postpartum Day”, and “On” vs. “Off-nest” behavior analyses of *EF1, EF2, EF3*, and *EF6***

Consistent with the “cleaned and complete” dataset for *EF*1 across the maternal homecage timepoints (Fig. 3c, Supplemental Fig. S21a, Table S16), we found a significant effect of maternal timepoint (REML, *P* = 1.89E-08), with the preconception stage revealed to have a higher *EF*1 score than the gestational and P20 timepoints for the imputed dataset (Supplemental Fig. S24b, Table S27). Looking across the postpartum observations, we again found a strong effect of postpartum day (REML, *P* = 3.15E-13) (Supplemental Fig. S24c, Table S28) on network strength, as well as an effect of whether dams were “on-nest” (*P* = 1.34E-18), with *EF*1 being higher when dams were off the nest (Table S29). For the imputed dataset, there was additionally a significant interaction of postpartum day and “on-nest” behavior (*P* = 0.0293), as the discrepancy between “on” and “off-nest” *EF*1 scores diminished over the postpartum period.

For *EF*2, unlike the “cleaned and complete” dataset (Supplemental Fig. S21b), using the imputed data we did not observe any effects of, nor interaction between, maternal timepoint and ELS (Supplemental Fig. S24e). However, over the postpartum observation points, we did find a significant effect of postpartum day (REML, *P* = 1.34E-05), with network scores observed to increase across the postpartum period (Supplemental Fig. 24f, Table S30). While the “cleaned and complete” dataset revealed a significant interaction of network activity by “on-nest” behavior and ELS (*P* = 0.0054) (Fig. 4d), there was no such effect of “on-nest” behavior nor any interactions observed for the imputed *EF*2 dataset.

For *EF*3, the imputed dataset revealed a significant effect of maternal timepoint (REML, *P* = 0.0420) (Supplemental Fig. S24h, Table S31), which was also observed in the “cleaned and complete” version of the analysis (REML, *P* = 0.0045) (Supplemental Fig. S21c, Table S17). Furthermore, while the evaluation of observations by postpartum day once again found a significant effect of maternal timepoint for *EF*3 (REML, *P* = 7.23E-05) (Supplemental Fig. S24i, Table S32), the effect of “on” vs. “off-nest” behavior observed in the “cleaned and complete” dataset (REML, *P* = 1.2E-05) (Fig. 4e, Table S23) was not found for the imputed dataset. *EF*3 scores showed an overall slight decrease over the postpartum observations.

For *EF*6, like the cleaned and complete dataset (Supplemental Fig. S21d), there was a significant effect of maternal timepoint detected (REML, *P* = 0.0111) (Supplemental Fig. S24k, Table S33). Additionally, there was a significant interaction of maternal timepoint and stress condition (*P* = 0.0378), which was not present in the “cleaned and complete” version of the analysis (Supplemental Fig. S21d, Table S18). When analyzing the postpartum observations, *EF*6 revealed a significant effect of postpartum day (REML, *P* = 0.0102) (Supplemental Fig. S24l, Table S34), and a highly significant effect of “on” vs. “off-nest” behavior (REML, *P* = 1.77E-34) (Table S35). These findings indicate a clear increase in *EF*6 score when dams are on the nest across the postpartum period, agreeing with the “cleaned and complete” dataset (Fig. 4f, Tables S24-S25). The imputed data also revealed a significant interaction of ELS and postpartum day (REML, *P* = 0.01188) that had trend significance in the “cleaned and complete” set (REML, *P* = 0.0542). These findings support the notion that the strength of stress-relevant *EF*s can change with life experience and also vary by engagement in naturalistic maternal behavior. See Table S26 for timepoint and analysis-specific animal numbers. In general, imputed findings for *EF1, EF2, EF3*, and *EF6* agree with those of the “cleaned and complete” findings, with the most noticeable differences being the lack of ELS-effect for increased *EF*2 across the maternal timecourse, and the loss of “on” vs. “off-nest” behavior effect for *EF*3 across postpartum observations.

**Imputation results for the analysis of *EF1* and *EF2* from repeated FIT**

When examining the FIT *EF*1 scores across both preconception and late postpartum maternal timepoints (Supplemental Fig. S33), we found a significant effect of FIT stage (REML, *P* = 2.23E-19) (Table S44), with scores generally increasing over the course of the task, as well as a significant effect of maternal timepoint (*P* = 1.37E-04) (Table S45). These findings align with the “cleaned and complete” version of the analysis (Fig. 6c, Tables S40-S41). Evaluating *EF*2 strength during the FIT and across maternal timepoints yielded several significant findings. Once again, there were significant effects of FIT task stage (REML, *P* = 9.43E-13) (Table S46) and maternal timepoint (*P* = 0.0022) (Table S47) detected, with a significant effect of ELS (*P* = 0.0366) also found, which was trend significant for the “cleaned and complete” dataset (*P* = 0.0633). ELS-exposed dams hint at higher *EF*2 engagement overall as compared to control dams. Agreeing with the “cleaned and complete” dataset, a significant interaction was found for the *EF*2 imputed dataset between FIT stage and ELS (*P* = 0.0043), with an additional interaction of FIT stage and maternal timepoint (*P* = 0.0431) also popping up. This last interaction was unique to the imputed *EF*2 dataset. These results highlight the striking postpartum increase in *EF*2 for ELS-exposed dams, particularly during the aggressor stage of the FIT. See Table S26 for timepoint and analysis-specific animal numbers.

**Prior early life stress exposure did not impact maternal networks**

We obtained measures of baseline-normalized *EF_on-nest_* strength across all timepoints and compared the trajectories for control and ELS-exposed dams (Fig. 2b, Supplemental Fig. S17). There was no significant effect of stress condition, but the clear changes in network strength across maternal timepoints (Tables S4, S11) were similar for both control and ELS-exposed dams. The *EF* strength trajectories of the “licking” and “lick-groom” networks (*EF_licking_* and *EF_lick-groom_* ) were also evaluated for both control and ELS-exposed dams (Fig. 2c, Supplementary Figs. S12, S18-S19) with no significant effect of or timepoint interaction with stress exposure detected. In the case of *EF_licking_*, there was a slightly significant effect of maternal timepoint (Supplemental Fig. S18); however, multiple comparisons revealed few specific timepoints to be significantly different by network strength (Table S12). We also identified a significant effect of maternal timepoint across both control and ELS-exposed dams for *EF_lick-groom_* (Supplemental Fig. S19), with a decrease in network strength observed as the maternal timecourse progressed following from preconception (Table S13).

As with *EF_on-nest_*, the normalized *EF_stage_* network trajectories were compared for both control and ELS-exposed dams over the maternal timecourse (Fig. 2f, Supplemental Fig. S20). Once again, there was no significant effect of stress, but there was an effect of maternal timepoint (Tables S6, S15). While not significant, the ELS group showed a more rapid decrease in postpartum network strength following its peak in early postpartum (P1) as compared to the control group.

*EF_on-nest_* network strength of ELS-exposed dams was greater during “on-nest” as compared to “off-nest” behavior across postpartum observation days (Supplemental Fig. S17, Table S10), while the *EF_stage_* network was not affected by this behavioral state (Supplemental Fig. S20). This mirrors what was observed in the control dams (Fig. 1d, Fig. 2g), confirming the distinction between the two novel electrical brain networks.

No effects of stress condition were found when evaluating *EF_on-nest_* and *EF_stage_* network strength during pup-licking and self-grooming behaviors at P3 or during pup retrieval at P4 (Supplemental Fig. S32). Taking a closer look at the correlation between “on-nest” behavior and network activity for individual dams, we tested whether ELS-exposure altered this relationship. Control dams showed a positive relationship between behavior and network activity at P8, but this differed significantly for ELS dams, which did not show this relationship (Supplemental Figs. S11, S25).

Independent of maternal network assessment across stages, there were no differences in maternal care behavior between control and ELS-exposed dams for percentage of total time spent on the nest (Supplemental Fig. S26b) or number of visits to the nest (Supplemental Fig. S26c, Table S36). However, we did find ELS exposure to interact with the duration of “on-nest” care visits across postpartum days (Supplemental Fig. S26d, Table S37), suggesting P3 as a potentially vulnerable timepoint for maternal care. This led us to further explore more specific P3 maternal behaviors, finding disrupted maternal care patterning and nursing behavior following ELS-exposure (Supplemental Fig. S27c-d). No differences were found in pup-directed licking/grooming or self-grooming behavior at this timepoint (Supplemental Fig. S27). To ensure that our non-blinded behavioral scoring was unbiased, an additional scoring method employing fixed observation times was performed in parallel at P3 by a blinded experimenter, revealing comparable data: ELS did not induce broad behavioral changes (Supplemental Fig. S28).

**Open Field Pup Retrieval Results at P4**

Dams with prior ELS-exposure did not show a significant elevation in the engagement of *EF*1 (Supplemental Fig. S30) or *EF*6 (Supplemental Fig. S31) during either the open field (no pups) or pup retrieval (pups) portions of the open field pup retrieval task at P4. Neither control nor ELS-exposed dams performed above a score of one (grouping 2 pups together) on the open field pup retrieval task, and there was no difference in performance by experimental group (Supplemental Fig. S30).

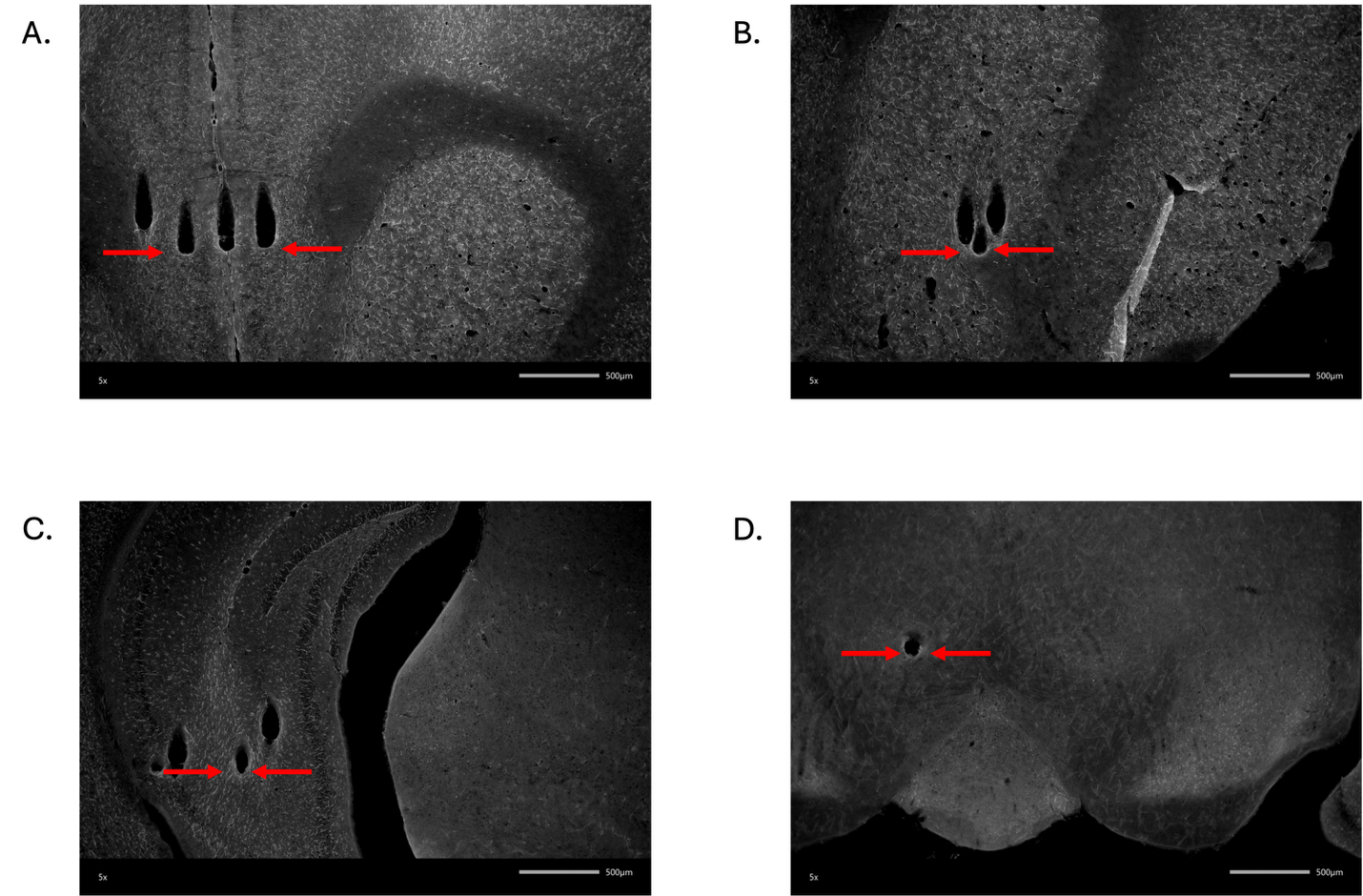

**b**

**a**

**d\**

**c**

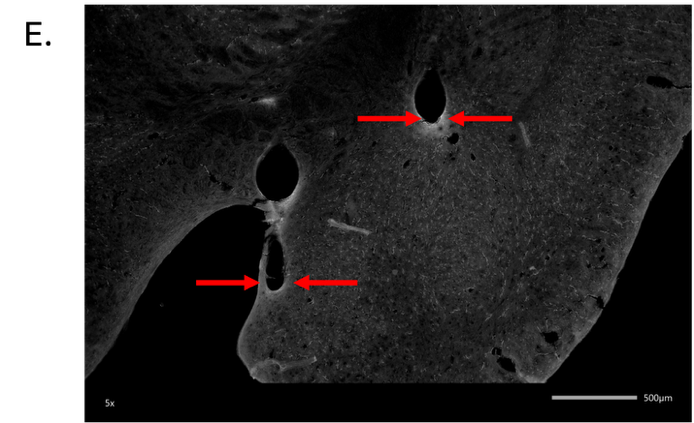

**e\**

### Supplemental Figure S1: Histological examination of electrode placement

Electrode wires were verified in the **a)** PrL, **b)** NAc, **c)** VHipp, **d)** VTA, and **e)** AMY (MeA/CeA). Red arrows indicate the ends of electrode wire tips.

PrL = prelimbic cortex, NAc = nucleus accumbens, VHipp = ventral hippocampus, VTA = ventral tegmental area, AMY = amygdala, MeA = medial amygdala, CeA = central amygdala
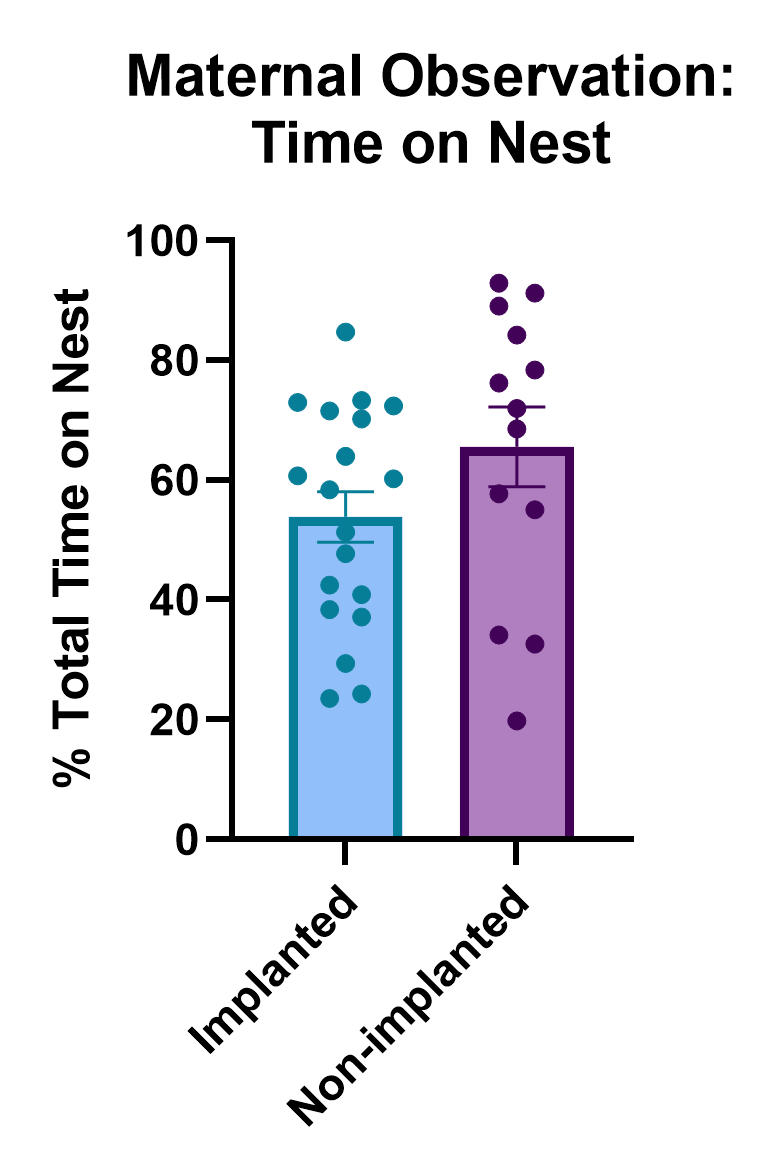

**a**

### Supplemental Figure S2: General maternal behavior is not impacted by neural implant

**a)** Total percentage of time spent on the nest for implanted and non-implanted control dams at P3 (unpaired t-test, *P* = 0.1307, *n* = 13-19/group).

Data are mean ± sem.

P = postpartum day.

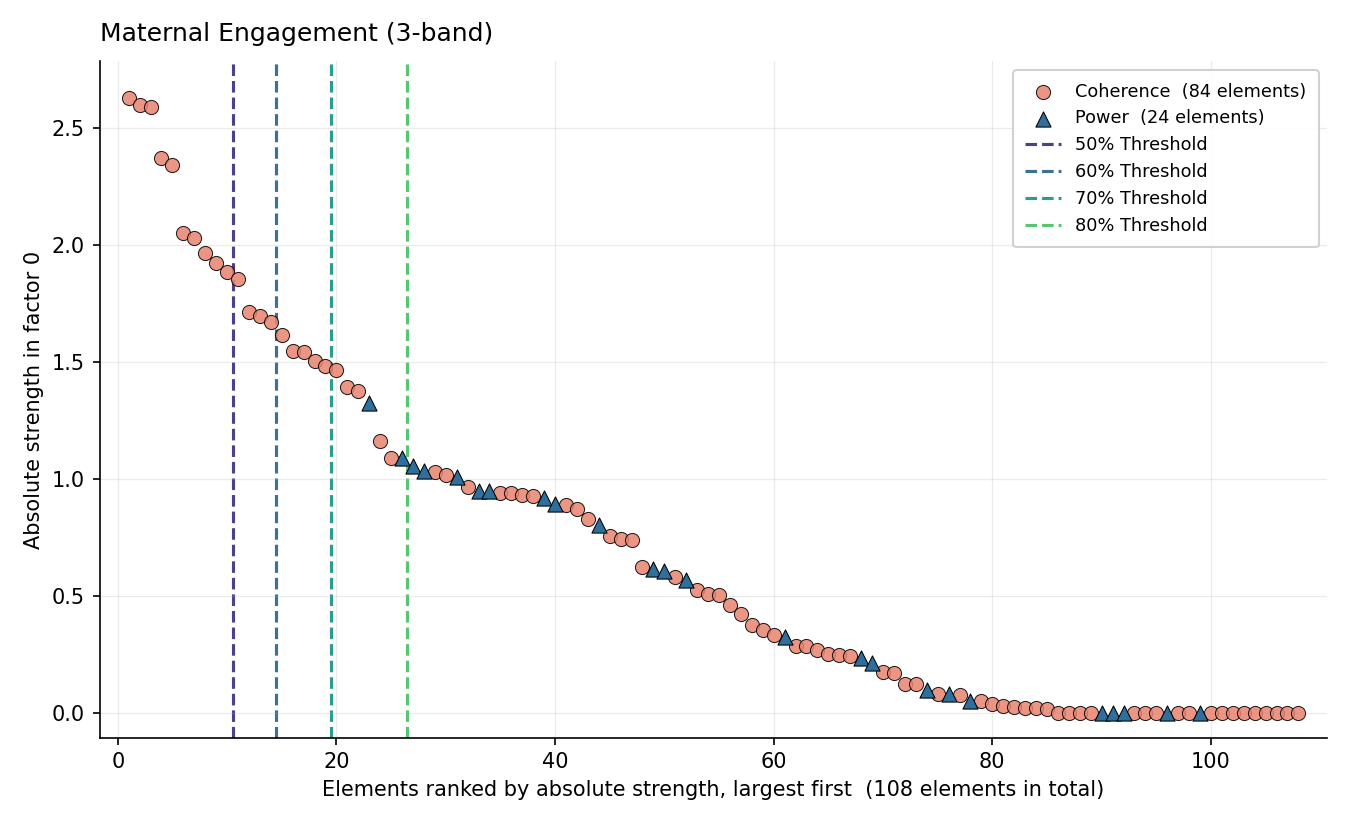

**a**

**b**

50% Threshold

80% Threshold

70% Threshold

60% Threshold

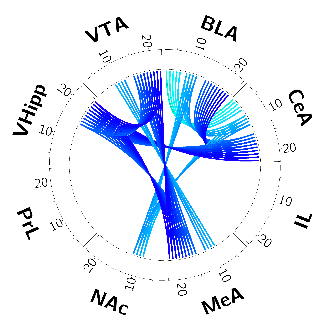

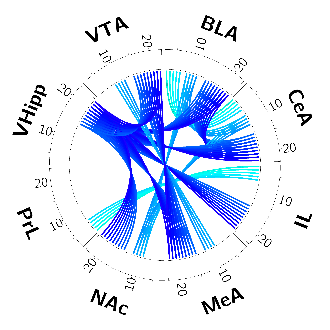

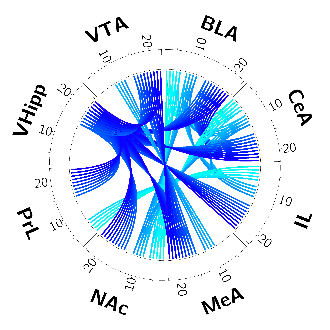

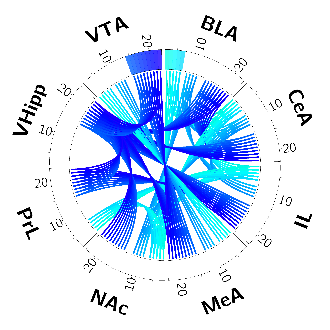

### Supplemental Figure S3: Determination of features threshold *EF_on-nest_* – guided frequency bands

**a)** Scree plot and novel *EF_on-nest_* specific elements when using guided frequency bands during training (2-7, 8-12, 14-23 Hz). Dotted lines indicate the element cut-off points for different thresholds as labeled. **b)** Circos plots visualizing the network specific *EF_on-nest_* contributing features at four different thresholds: 50%, 60%, 70%, 80%.

PrL = Prelimbic cortex, IL = infralimbic cortex, NAc = nucleus accumbens, BLA = basolateral amygdala, CeA = central amygdala, MeA = medial amygdala, VHipp = ventral hippocampus, VTA = ventral tegmental area, *EF* = *Electome Factor*

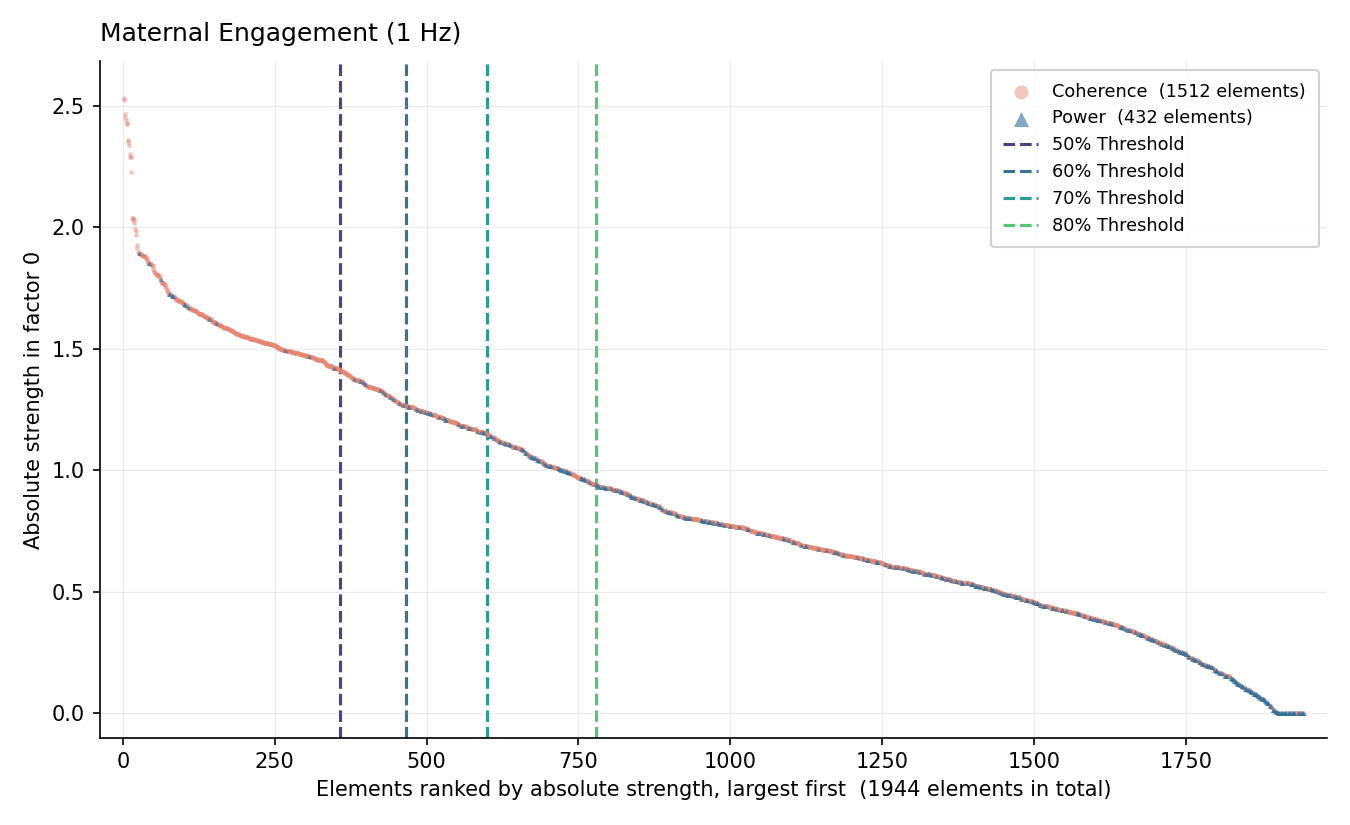

**a**

80% Threshold

70% Threshold

60% Threshold

**b**

50% Threshold

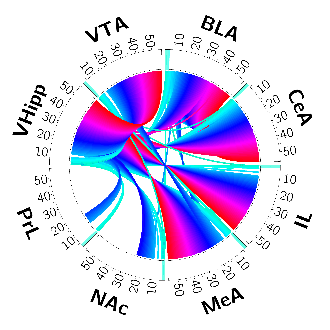

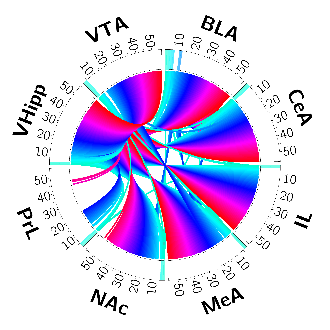

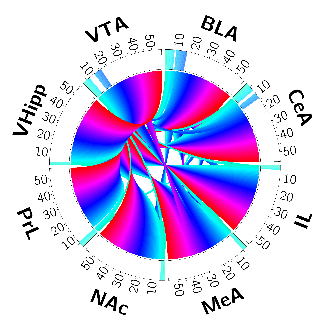

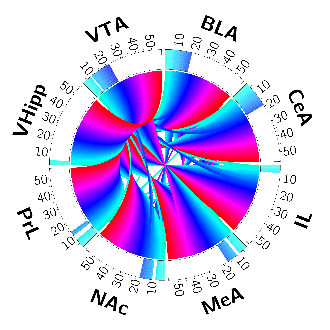

### Supplemental Figure S4: Determination of features threshold for *EF_on-nest_* – 1 Hz frequency bands (*EF_on-nest-1-Hz_)*

**a)** Scree plot and novel *EF_on-nest-1-Hz_* specific elements when using 1 Hz frequency bands during training. Dotted lines indicate the element cut-off points for different thresholds as labeled. **b)** Circos plots visualizing the network specific *EF_on-nest-1-Hz_* contributing features at four different thresholds: 50%, 60%, 70%, and 80%.

PrL = Prelimbic cortex, IL = infralimbic cortex, NAc = nucleus accumbens, BLA = basolateral amygdala, CeA = central amygdala, MeA = medial amygdala, VHipp = ventral hippocampus, VTA = ventral tegmental area, *EF* = *Electome Factor*

**Network Specific Feature Visualization**

**Dot Size = Absolute Strength | Dot Color = Relative Specificity**

**a**

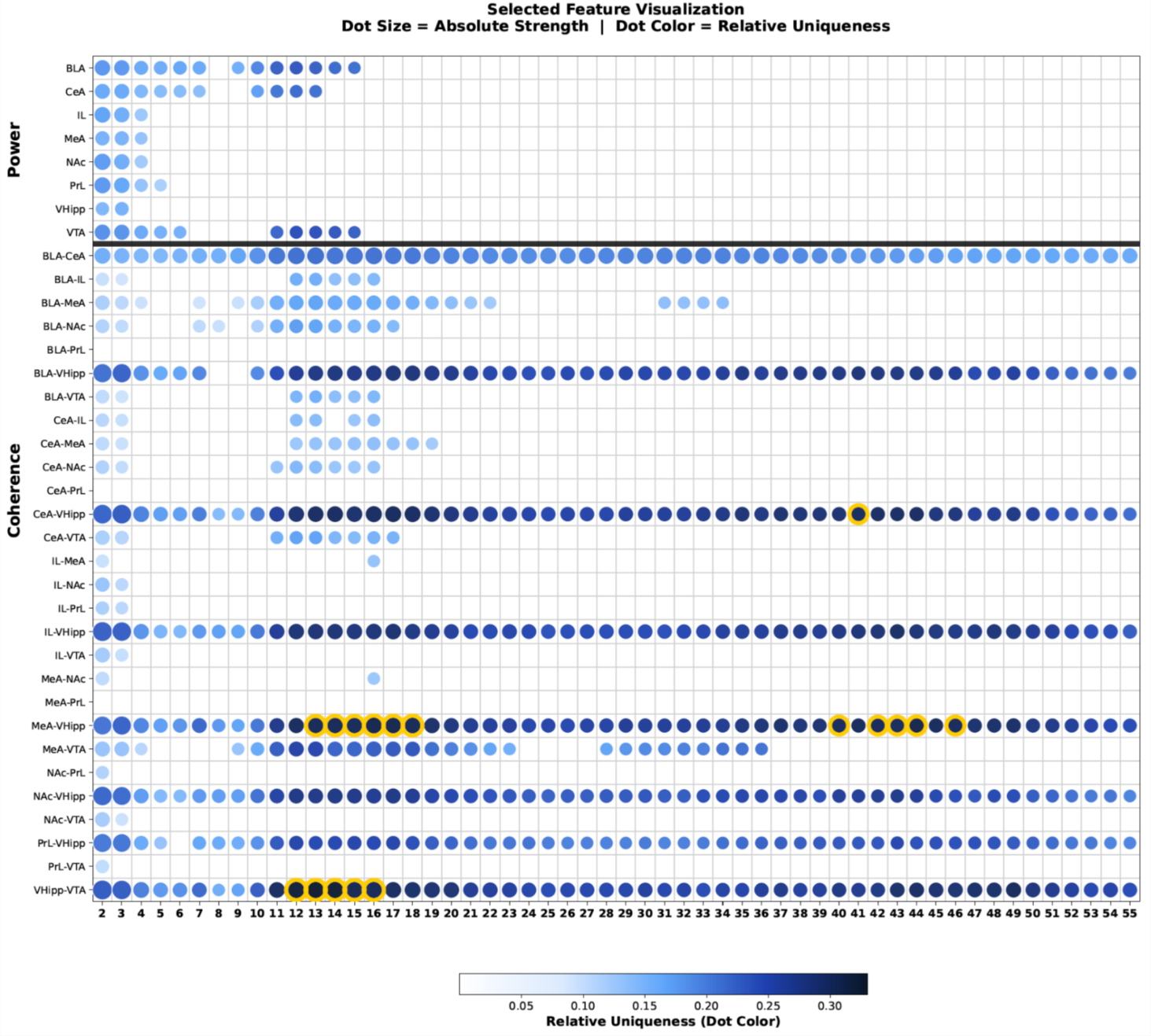

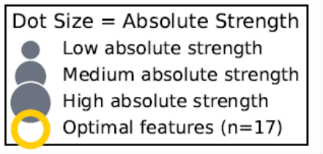

**Relative Specificity (Dot Color)**

### Supplemental Figure S5: Network-specific feature contribution for *EF_on-nest_* – 1 Hz frequency bands (*EF_on-nest-1-Hz_)*

**a)** Network-specific feature contribution for *EF_on-nest-1-Hz_* at 70% threshold. Absolute strength of feature contribution is indicated by dot size, and *EF_on-nest-1-Hz_* network specificity is indicated by darkness of color. Gold circles highlight features that are particularly important contributors to the network, reaching threshold levels for both absolute strength and relative specificity contributions.

PrL = Prelimbic cortex, IL = infralimbic cortex, NAc = nucleus accumbens, BLA = basolateral amygdala, CeA = central amygdala, MeA = medial amygdala, VHipp = ventral hippocampus, VTA = ventral tegmental area, *EF* = *Electome Factor*

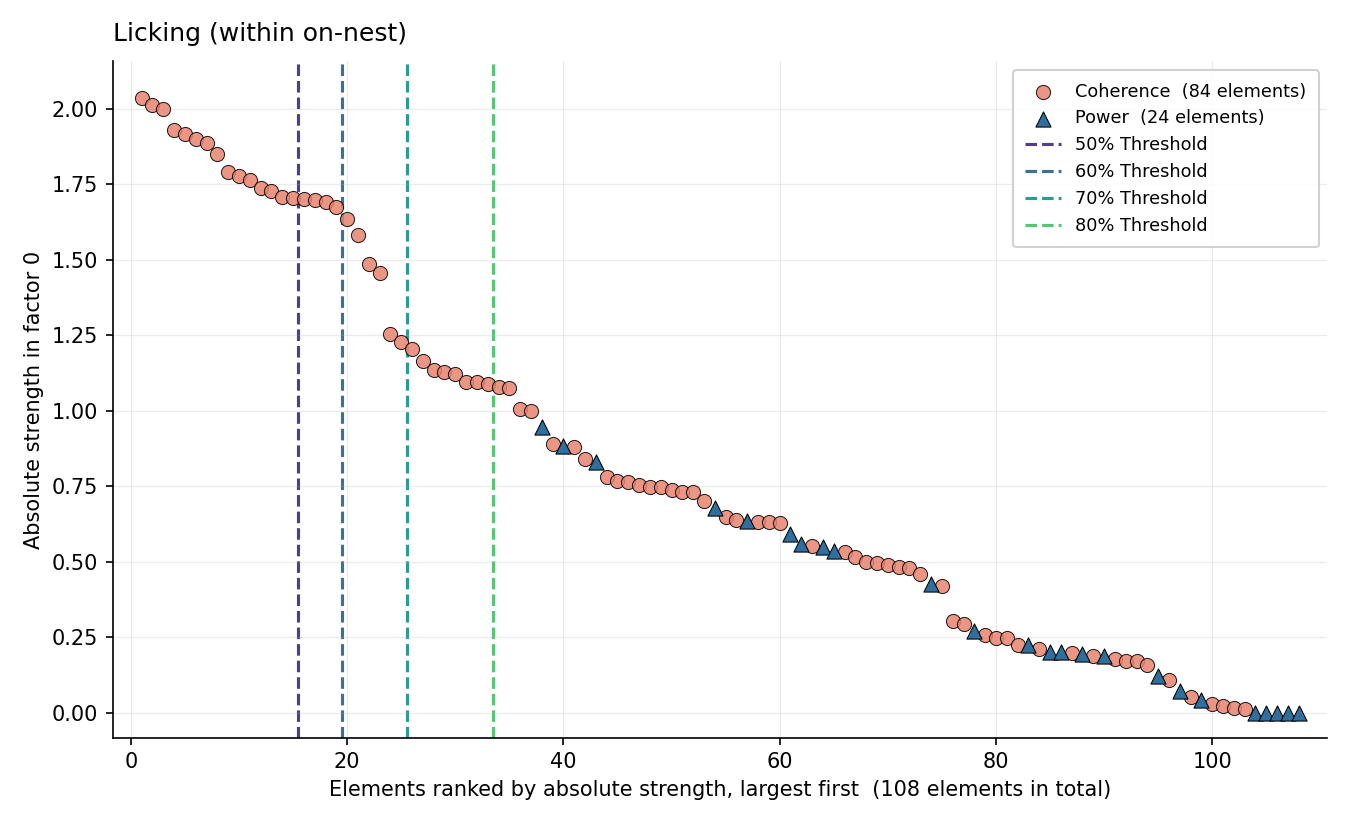

**a**

**b**

70% Threshold

80% Threshold

60% Threshold

50% Threshold

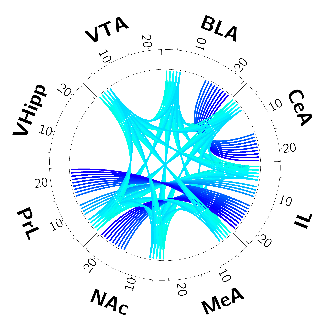

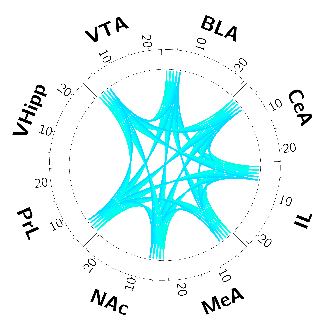

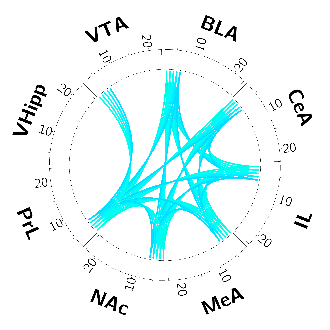

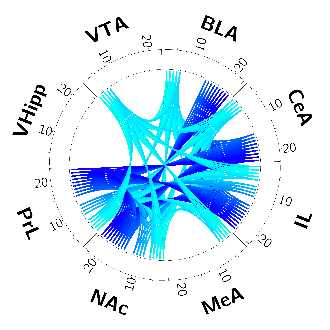

### Supplemental Figure S6: Determination of features threshold for *EF_licking_* – guided frequency bands

**a)** Scree plot and novel *EF_licking_* specific elements when using guided frequency bands during training (2-7, 8-12, 14-23 Hz). Dotted lines indicate the element cut-off points for different thresholds as labeled. **b)** Circos plots visualizing the network specific *EF_licking_* contributing features at four different thresholds: 50%, 60%, 70%, and 80%.

PrL = Prelimbic cortex, IL = infralimbic cortex, NAc = nucleus accumbens, BLA = basolateral amygdala, CeA = central amygdala, MeA = medial amygdala, VHipp = ventral hippocampus, VTA = ventral tegmental area, *EF* = *Electome Factor*

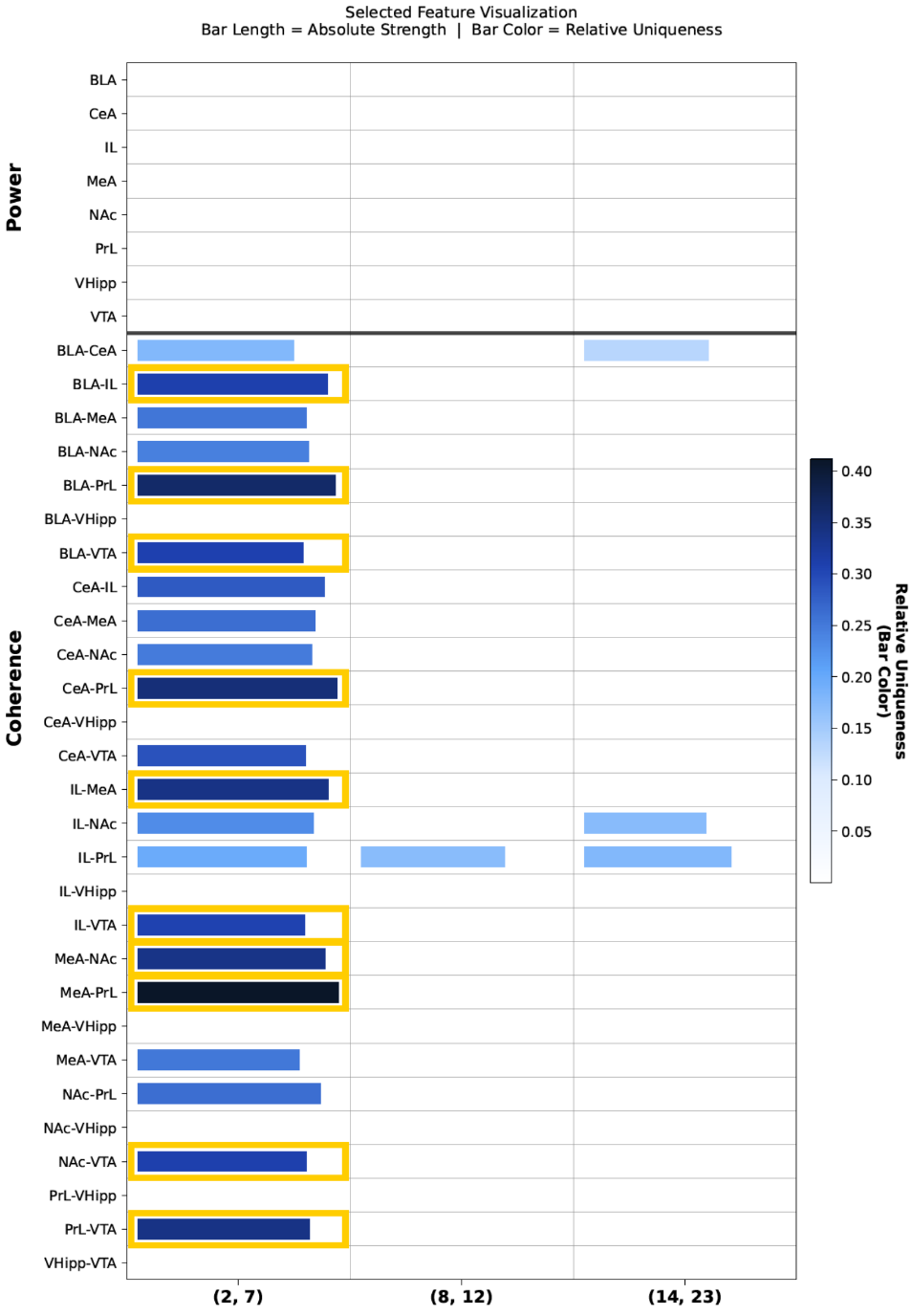

**Network Specific Feature Visualization**

**Bar Length = Absolute Strength | Bar Color = Relative Specificity**

**a**

**Relative Specificity (Bar Color)**

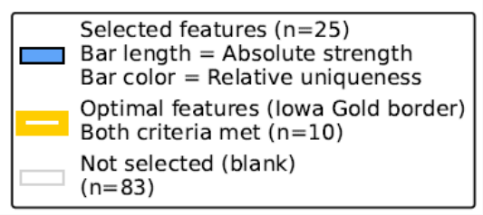

### Supplemental Figure S7: Network-specific feature contribution for *EF_licking_* – guided frequency bands

**a)** Network-specific feature contribution for *EF_licking_* using guided frequency bands during training (2-7, 8-12, 14-23 Hz) at 70% threshold. Absolute strength of feature contribution is indicated by bar length, and *EF_licking_* network specificity is indicated by darkness of color. Gold borders highlight features that are particularly important contributors to the network, reaching threshold levels for both absolute strength and specificity contributions.

PrL = Prelimbic cortex, IL = infralimbic cortex, NAc = nucleus accumbens, BLA = basolateral amygdala, CeA = central amygdala, MeA = medial amygdala, VHipp = ventral hippocampus, VTA = ventral tegmental area, *EF* = *Electome Factor*

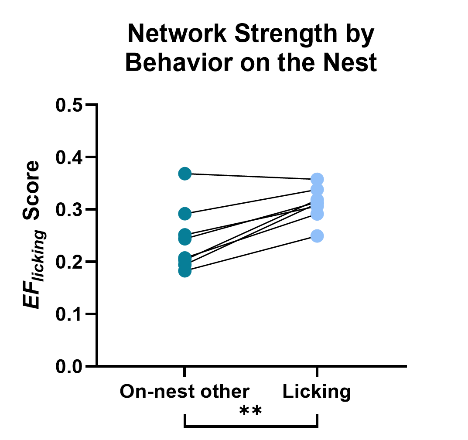

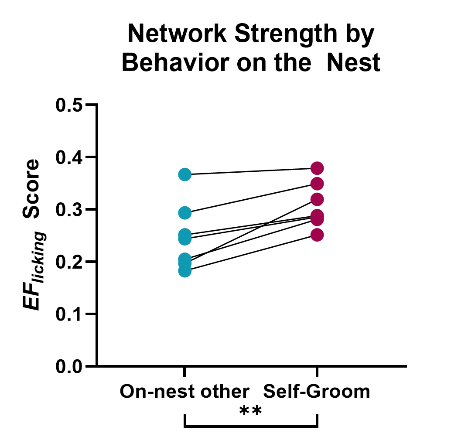

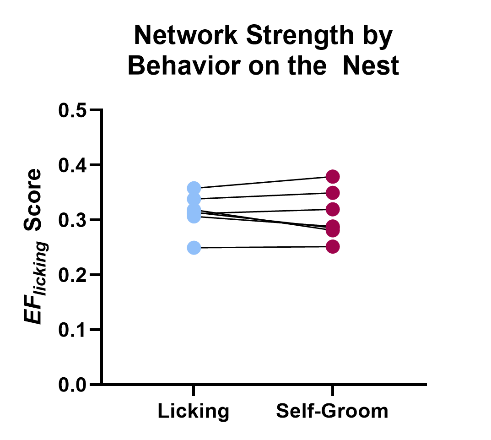

**c**

**b**

**a**

### Supplemental Figure S8: *EF_licking_* discriminates licking/grooming of pups from other on-nest behaviors but not from self-grooming behavior on the nest

**a)** Strength of *EF_licking_* during licking (of pups) on the nest (light blue) vs. all other behavior on the nest (teal) to assess behavioral discrimination of the network (paired t-test, *P* = 0.0024, *n* = 8). **b)** Strength of *EF_licking_* during dam self-grooming on the nest (red) vs. all other behavior on the nest (teal, paired t-test, *P* = 0.0060, *n* = 7) and **c)** during licking (of pups) on the nest (light blue) vs. self-grooming on the nest (red, paired t-test, *P* = 0.4476, *n* = 7).

Data shown are raw mean ± sem. ***P* < 0.01.

Statistical analyses were performed on fourth root equivalent transformed data. This was done to help stabilize variance and make the data more normally distributed.

*EF* = *Electome Factor*

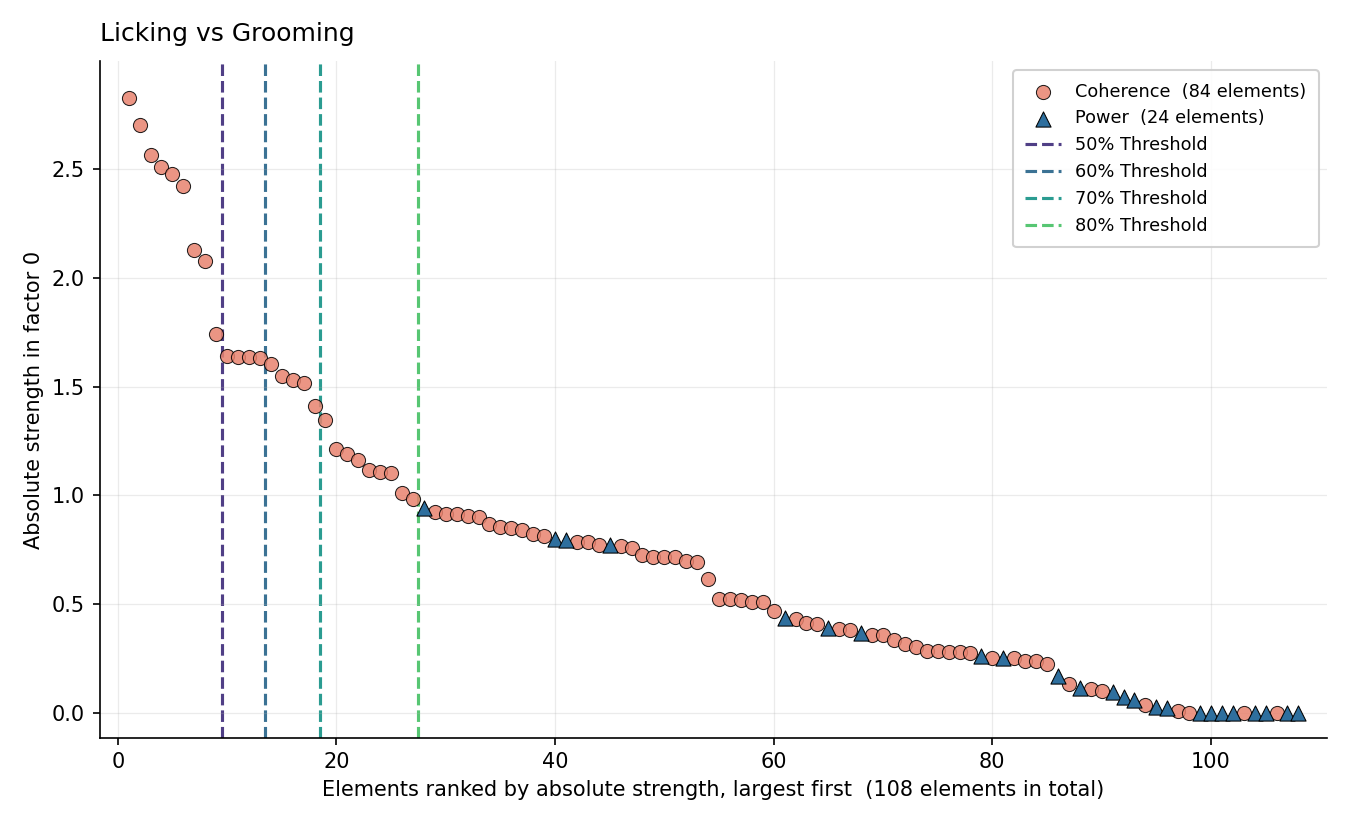

**a**

**b**

50% Threshold

60% Threshold

70% Threshold

80% Threshold

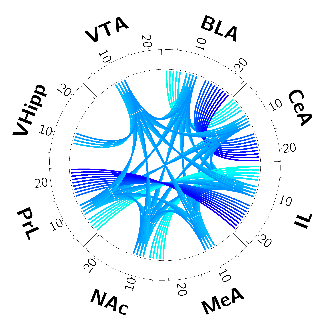

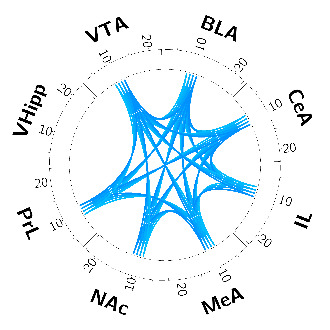

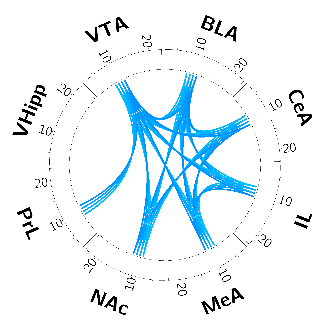

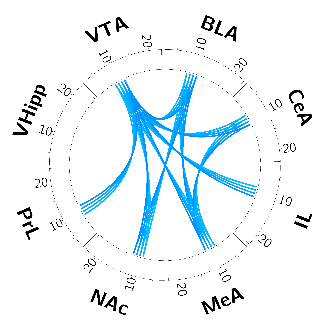

### Supplemental Figure S9: Determination of features threshold for *EF_lick-groom_* – guided frequency bands

**a)** Scree plot and novel *EF_lick-groom_* specific elements when using guided frequency bands during training (2-7, 8-12, 14-23 Hz). Dotted lines indicate the element cut-off points for different thresholds as labeled. **b)** Circos plots visualizing the network specific *EF_lick-groom_* contributing features at four different thresholds: 50%, 60%, 70%, and 80%.

PrL = Prelimbic cortex, IL = infralimbic cortex, NAc = nucleus accumbens, BLA = basolateral amygdala, CeA = central amygdala, MeA = medial amygdala, VHipp = ventral hippocampus, VTA = ventral tegmental area, *EF* = *Electome Factor*

**a**

**Network Specific Feature Visualization**

**Bar Length = Absolute Strength | Bar Color = Relative Specificity**

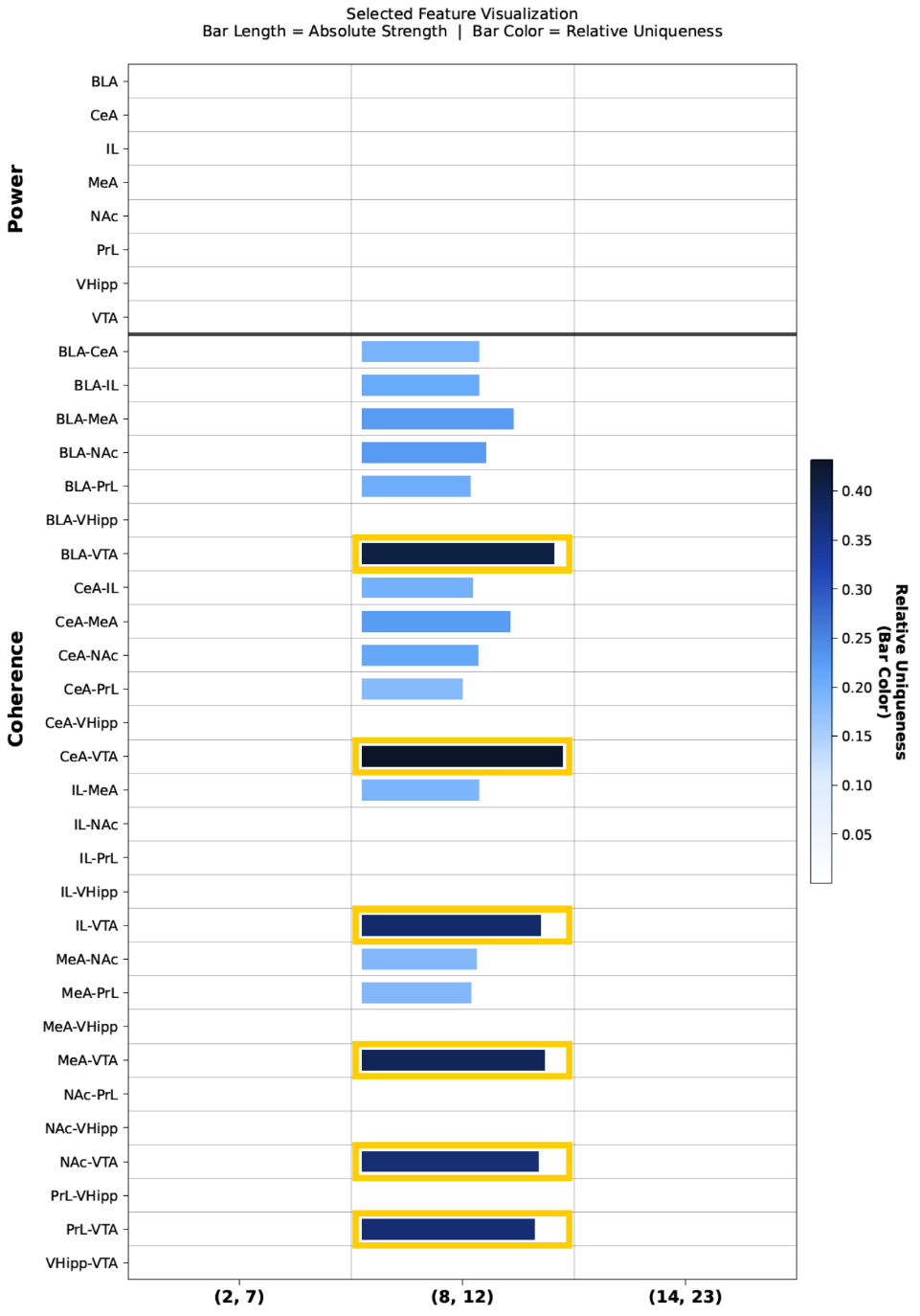

**Relative Specificity (Bar Color)**

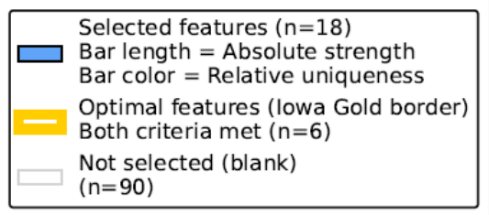

### Supplemental Figure S10: Network-specific feature contribution for *EF_lick-groom_* – guided frequency bands

**a)** Network-specific feature contribution for *EF_lick-groom_* using the guided frequency bands at 70% threshold. Absolute strength of feature contribution is indicated by bar length and *EF_lick-groom_* network specificity is indicated by darkness of color. Gold borders highlight features that are particularly important contributors to the network, reaching threshold levels for both absolute strength and specificity contributions. PrL = Prelimbic cortex, IL = infralimbic cortex, NAc = nucleus accumbens, BLA = basolateral amygdala, CeA = central amygdala, MeA = medial amygdala, VHipp = ventral hippocampus, VTA = ventral tegmental area, *EF* = *Electome Factor*

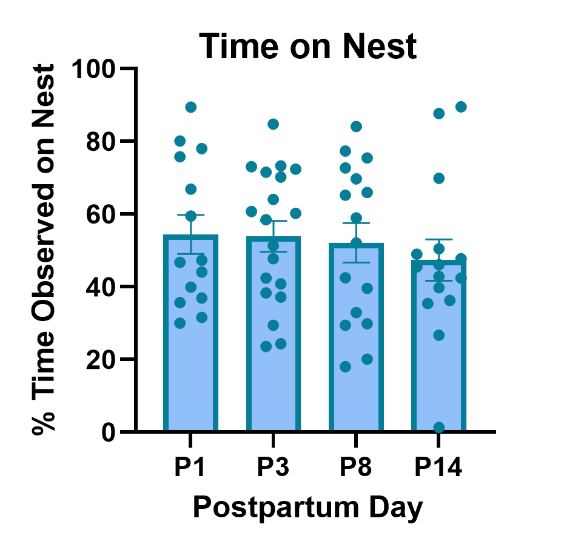

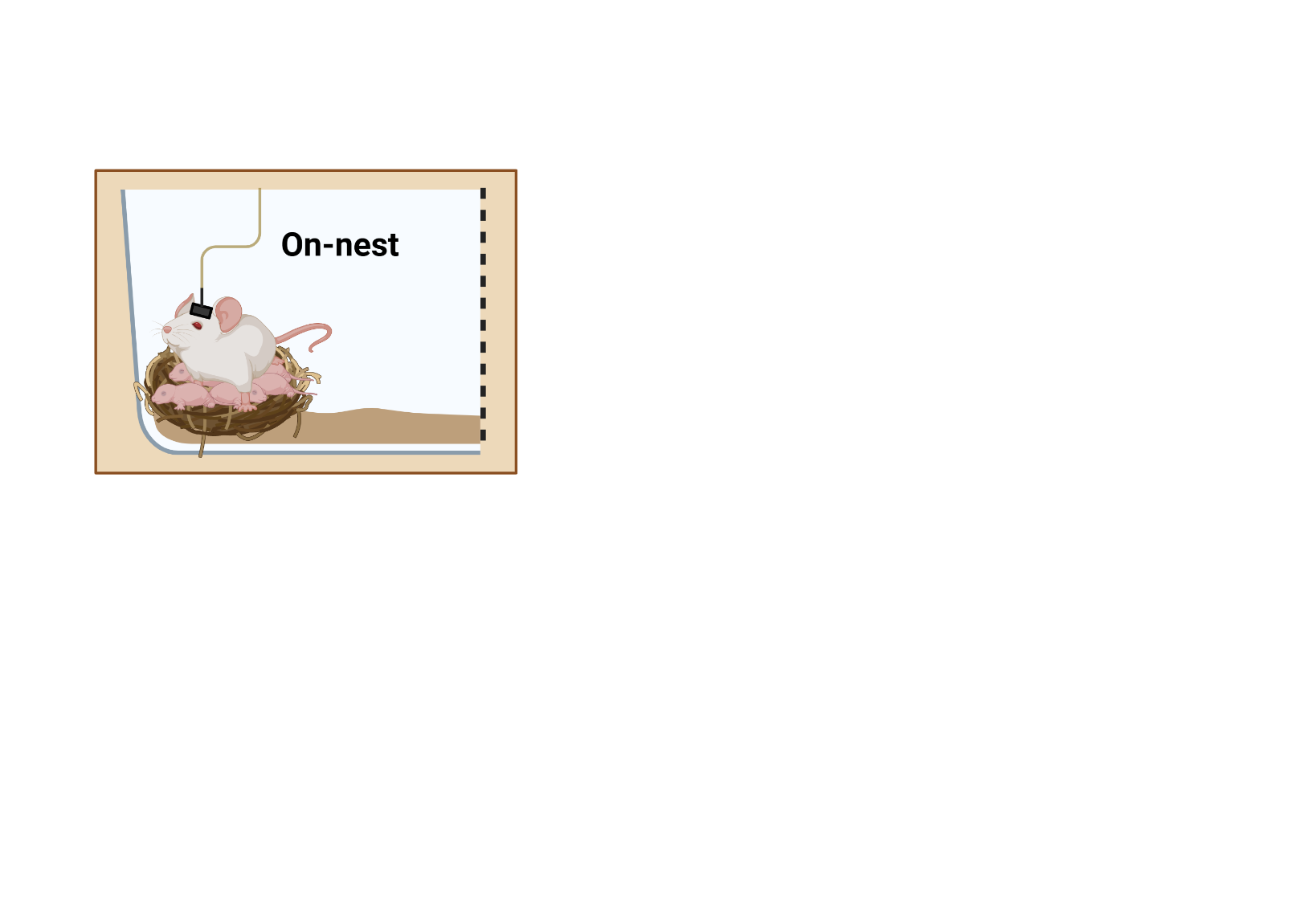

**b**

**a**

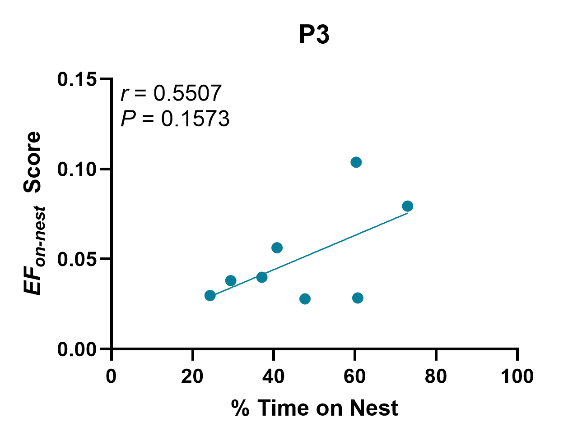

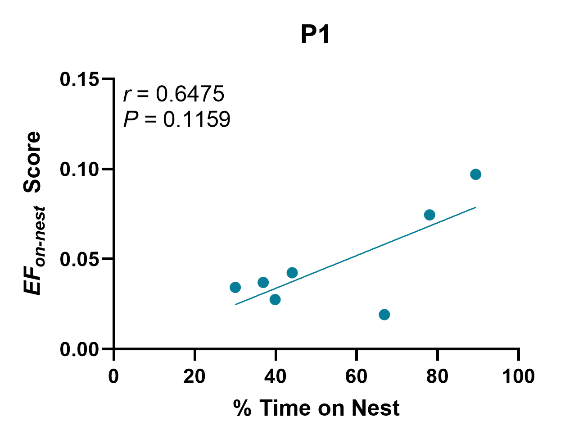

**d**

**c**

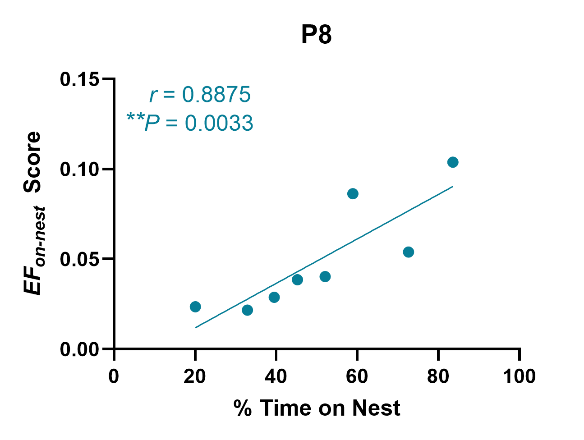

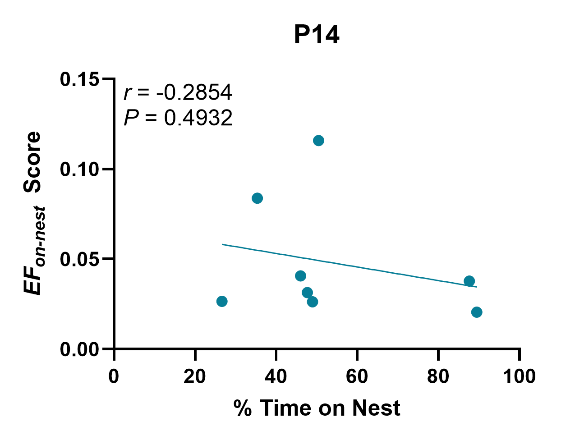

**f**

**e**

### Supplemental Figure S11: Time on nest across postpartum observations and correlations with *EF_on-nest_* score

**a)** Illustration of a dam spending time on the nest. **b**) Percent total time “on nest” across four postpartum observation timepoints**. c-f)** Correlations of individual percent time on the nest and average *EF_on-nest_* score during behavior on the nest across four postpartum timepoints: **c)** P1, **d)** P3, **e)** P8 (*r* = 0.8875, *P* = 0.0033, *n* = 8), **f)** P14. Panel **a** was created in BioRender. Mitchell, SB. (2026) https://BioRender.com/x9540la.

Data shown are raw mean ± sem. ***P* < 0.01. Statistical analysis was completed using Pearson’s correlation (**c-f**) on fourth root transformed data. This was done to help stabilize variance and make the data more normally distributed.*EF* = *Electome Factor*, P = postpartum day, *r* = Pearson’s correlation coefficient

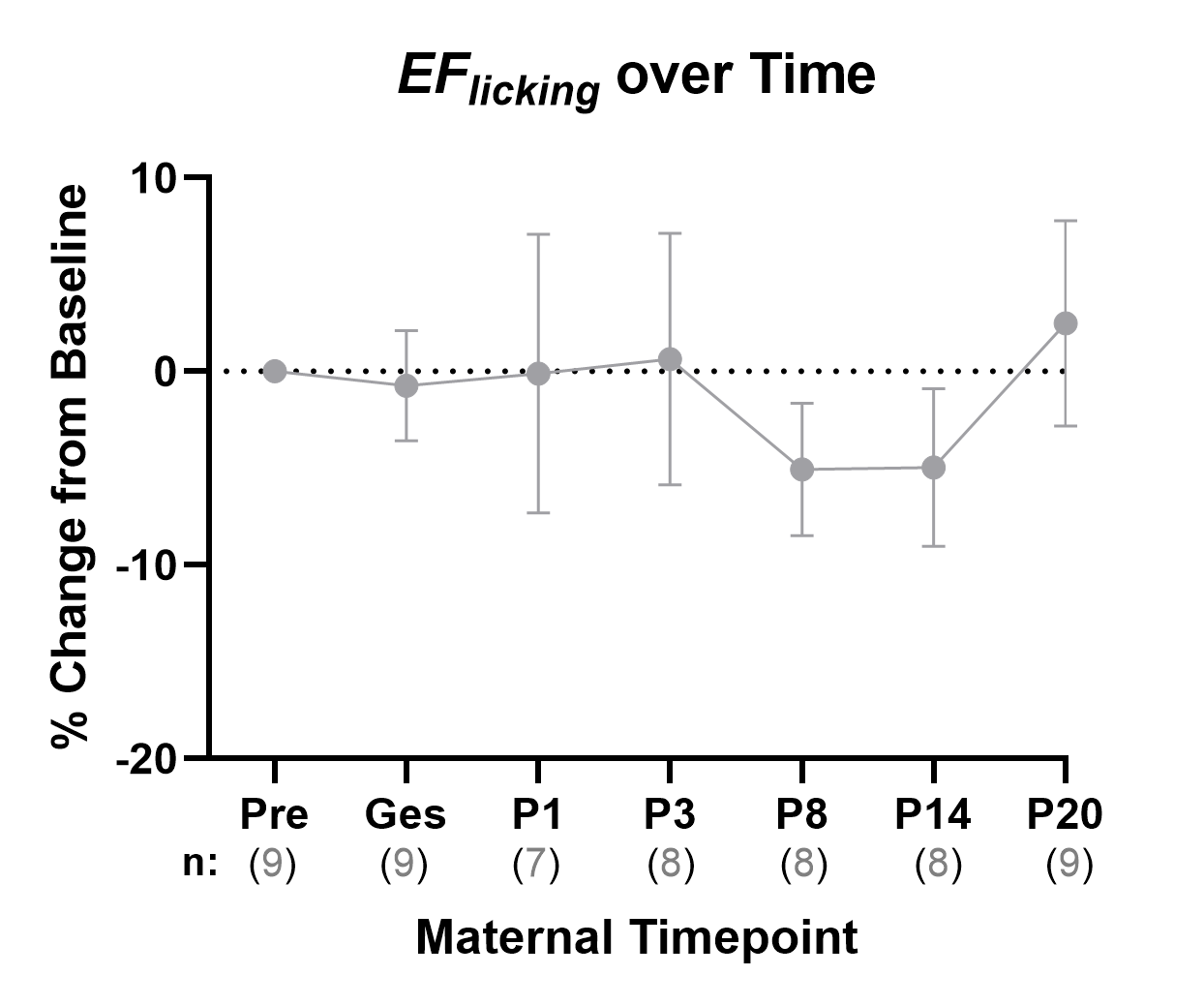

**a**

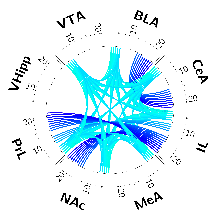

### Supplemental Figure S12: Trajectory of *EF_licking_* across all maternal timepoints

**a)** The strength of *EF_licking_* across observation timepoints calculated as the percent change from baseline (preconception). Timepoint specific *n* can be found below each respective observation.

Data shown are raw mean ± sem.

Statistical analysis (one-way RM REML) of mean *EF* scores was performed on fourth root transformed data. This was done to help stabilize variance and make the data more normally distributed. For the normalized *EF* trajectory data**,** the transformation was performed prior to calculating the percent change from baseline. All data shown are non-transformed.

Pre = preconception, Ges = gestation, P = postpartum day, RM = repeated measures, REML = restricted maximum likelihood, *EF* = *Electome Factor*

**a**

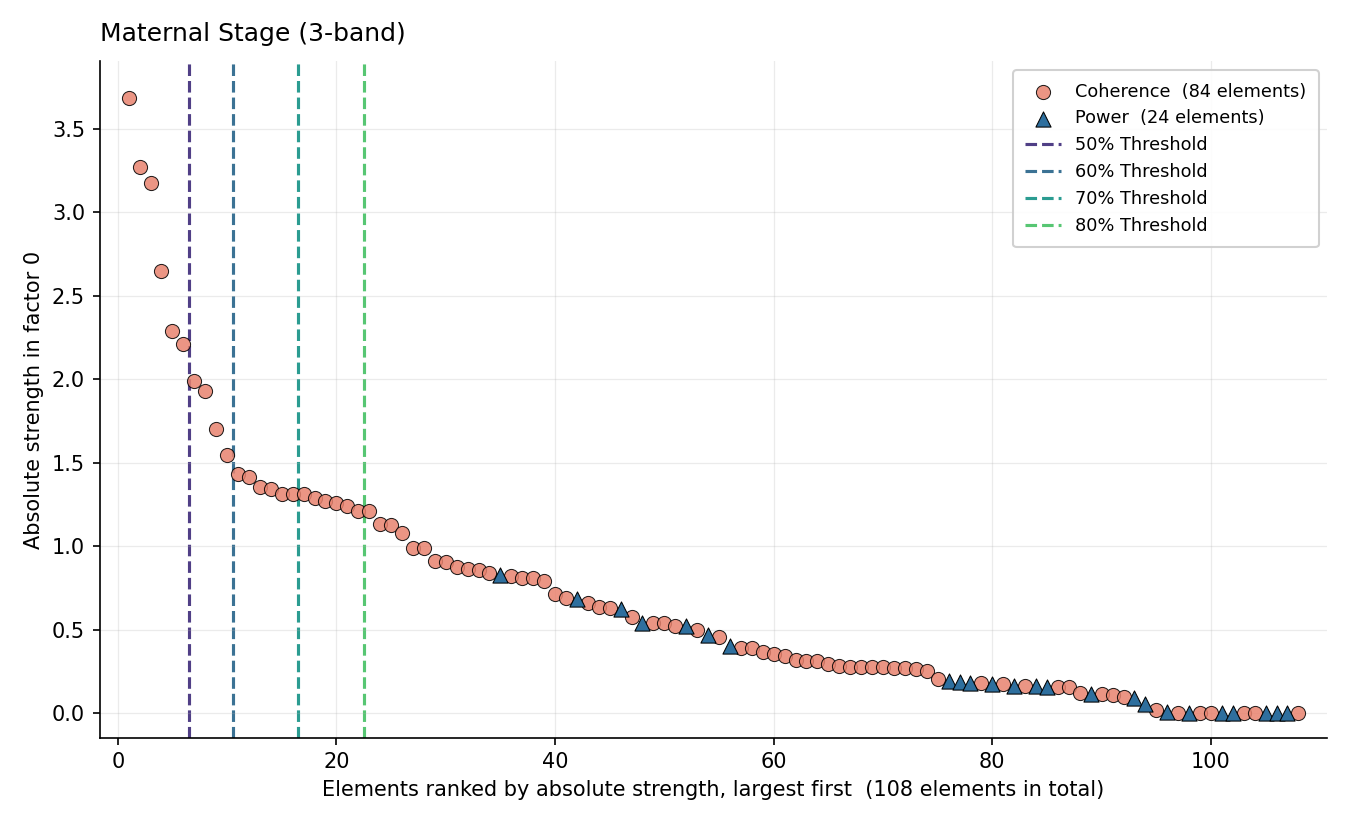

**b**

80% Threshold

70% Threshold

60% Threshold

50% Threshold

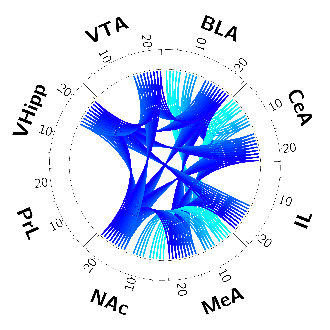

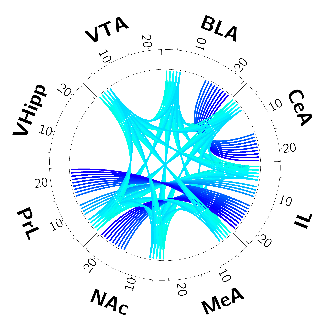

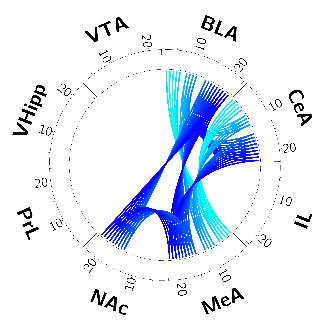

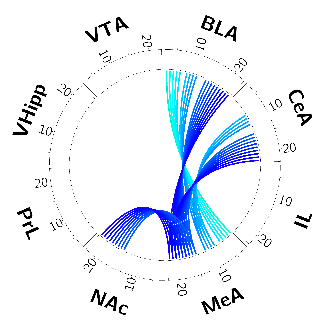

### Supplemental Figure S13: Determination of selected features threshold for *EF_stage_* – guided frequency bands

**a)** Scree plot and novel *EF_stage_* specific elements when using guided frequency bands during training (2-7, 8-12, 14-23 Hz). Dotted lines indicate the element cut-off points for different thresholds as labeled. **b)** Circos plots visualizing the network specific *EF_stage_* contributing features at four different thresholds: 50%, 60%, 70%, and 80%.

PrL = Prelimbic cortex, IL = infralimbic cortex, NAc = nucleus accumbens, BLA = basolateral amygdala, CeA = central amygdala, MeA = medial amygdala, VHipp = ventral hippocampus, VTA = ventral tegmental area, *EF* = *Electome Factor*

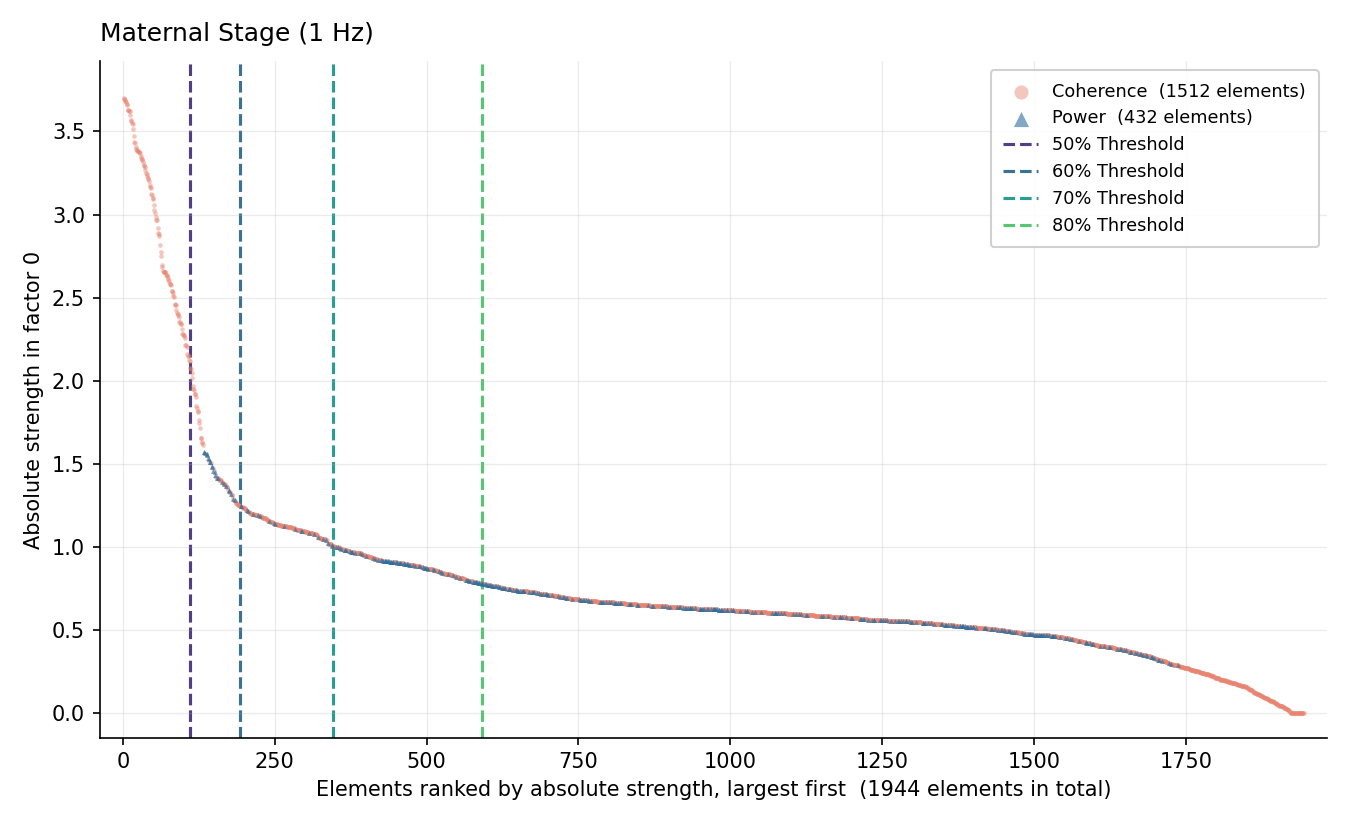

**a**

80% Threshold

70% Threshold

60% Threshold

50% Threshold

**b**

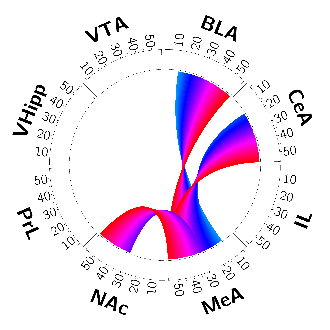

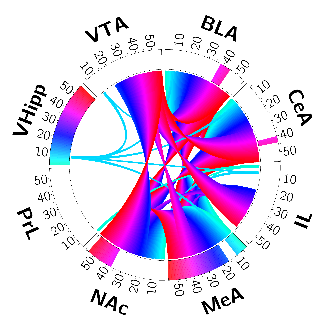

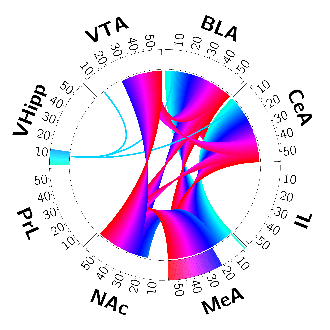

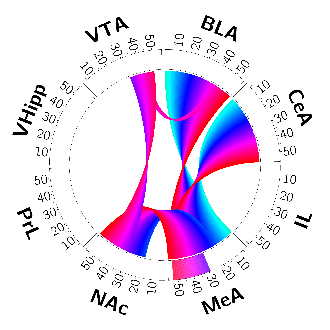

### Supplemental Figure S14: Determination of selected features threshold for *EF_stage_* – 1 Hz frequency bands (*EF_stage-1-Hz_)*

**a)** Scree plot and novel *EF_stage-1-Hz_* specific elements when using 1 Hz frequency bands during training. Dotted lines indicate the element cut-off points for different thresholds as labeled. **b)** Circos plots visualizing the network specific *EF_stage-1-Hz_* contributing features at four different thresholds: 50%, 60%, 70% and 80%.

PrL = Prelimbic cortex, IL = infralimbic cortex, NAc = nucleus accumbens, BLA = basolateral amygdala, CeA = central amygdala, MeA = medial amygdala, VHipp = ventral hippocampus, VTA = ventral tegmental area, *EF* = *Electome Factor*

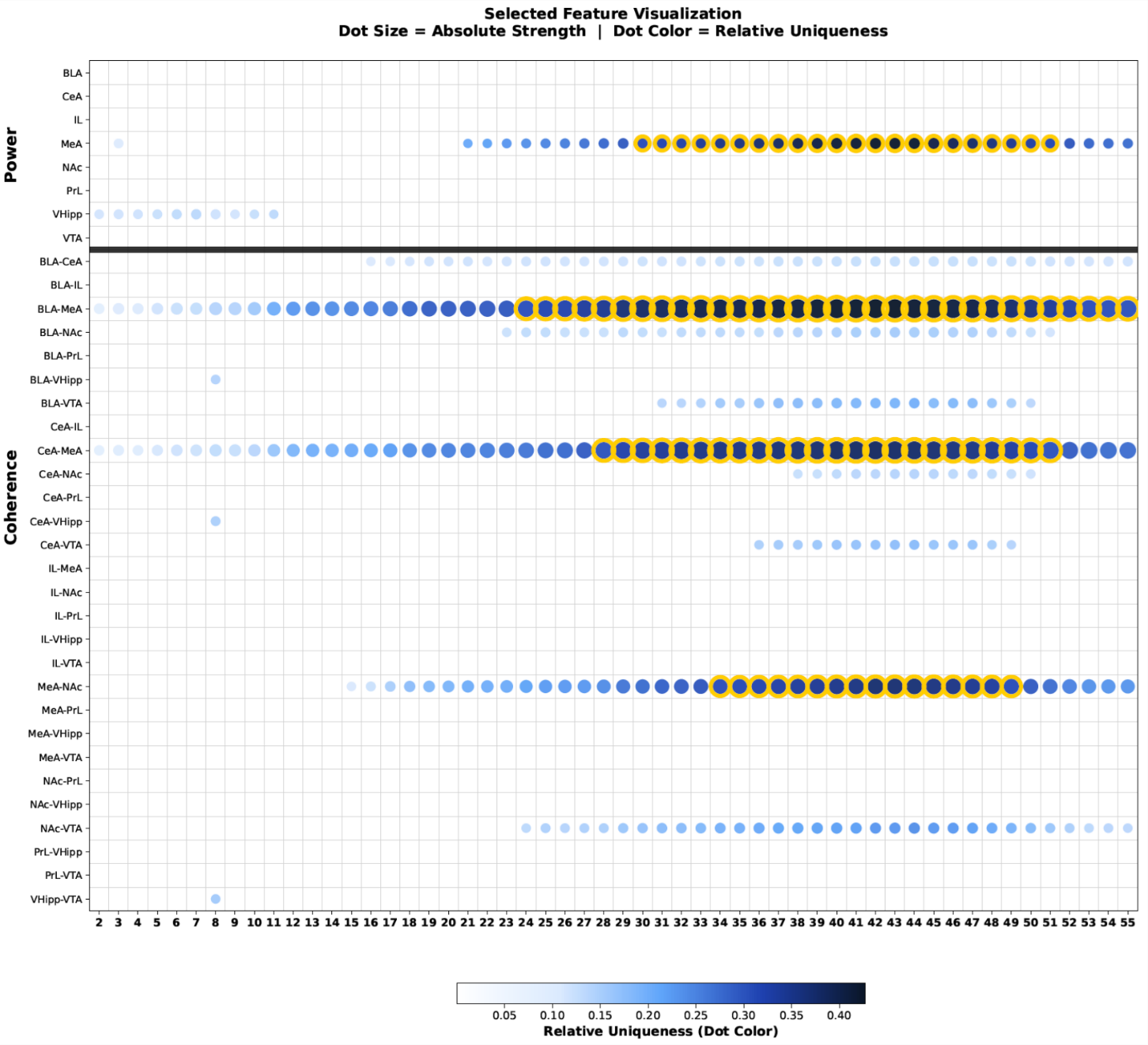

**a**

**Network Specific Feature Visualization**

**Dot Size = Absolute Strength | Dot Color = Relative Specificity**

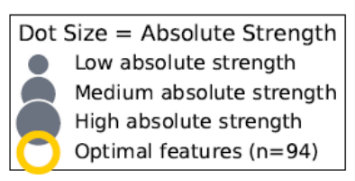

**Relative Specificity (Dot Color)**

### Supplemental Figure S15: Network-specific feature contribution for *EF_stage_* – 1 Hz frequency bands (*EF_stage-1-Hz_)*

**a)** Network-specific feature contribution for *EF_stage-1-Hz_* at 70% threshold. Absolute strength of feature contribution is indicated by dot size and *EF_stage-1-Hz_* network specificity is indicated by darkness of color. Gold circles highlight features that are particularly important contributors to the network, reaching threshold levels for both absolute strength and specificity contributions.

PrL = Prelimbic cortex, IL = infralimbic cortex, NAc = nucleus accumbens, BLA = basolateral amygdala, CeA = central amygdala, MeA = medial amygdala, VHipp = ventral hippocampus, VTA = ventral tegmental area, *EF* = *Electome Factor*

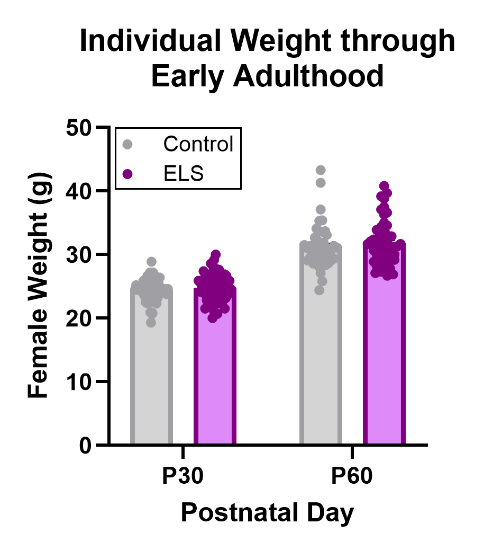

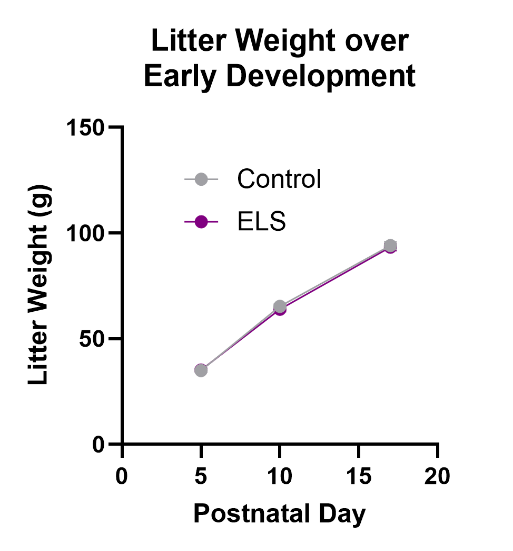

**b**

**a**

### Supplemental Figure S16: Developmental animal weight is unaffected by early life stress

**a)** Control and ELS-exposed litter weights over early development. Litters were culled to 8 pups each (*n* = 19 litters/group). **b)** Individual weights of control and ELS-exposed female mice at an adolescent (P30) and early adult (P60) timepoint (*n* = 45-79 individuals/group from 19 total litters/group).

Data are mean ± sem.

Statistics were performed using the two-way RM ANOVA **(a)** or unpaired t-test **(b)**.

ELS = early life stress, P = postnatal day

**b**

**a**

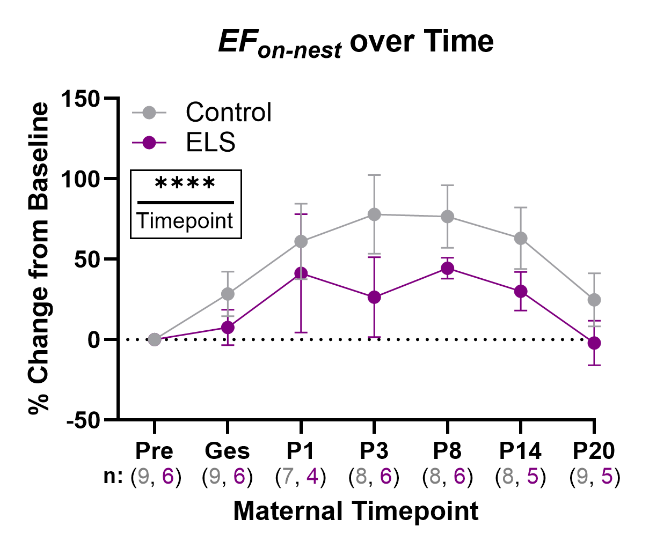

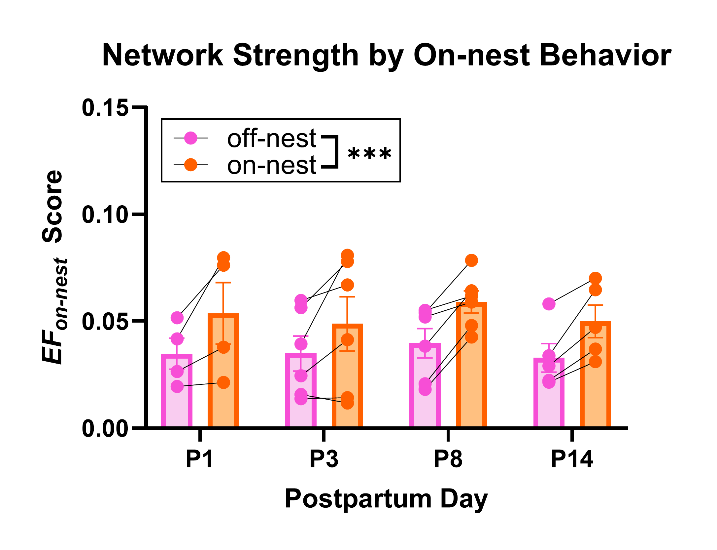

### Supplemental Figure S17: ELS-exposure does not impact *EF_on-nest_* performance or maternal trajectory

**a)** Mean *EF_on-nest_* score separated by “on” and “off-nest” behavior across observed maternal timepoints (REML, behavior effect, *P* = 5.27E-04, *n*=4-6) for ELS-exposed dams. Multiple comparisons by postpartum day can be found in Table S10. **b)** Mean *EF_on-nest_* (REML, stage effect, *P* = 1.89E-05) scores across observed maternal timepoints normalized to the preconception baseline level for both control and ELS-exposed dams. Timepoint specific *n*/group noted beneath each stage. Multiple comparisons of individual timepoints can be found in Table S11.

Data are mean ± sem. ****P* < 0.001, *****P* < 0.0001

Statistical analyses were completed using two-way RM REML. All statistical analyses of mean *EF* scores were performed on fourth root transformed data. For the normalized *EF* trajectory data**,** the transformation was performed prior to calculating the percent change from baseline. All data shown are non-transformed. If a significant effect of the repeated measure was found, a False Discovery Rate (FDR) correction for multiple comparisons was performed to specify which timepoints or measures were significantly different.

*EF* = *Electome Factor*, P = postpartum day, Pre = preconception, Ges = gestation, RM = repeated measures, REML = Restricted Maximum Likelihood, FDR = False Discovery Rate correction for multiple comparisons

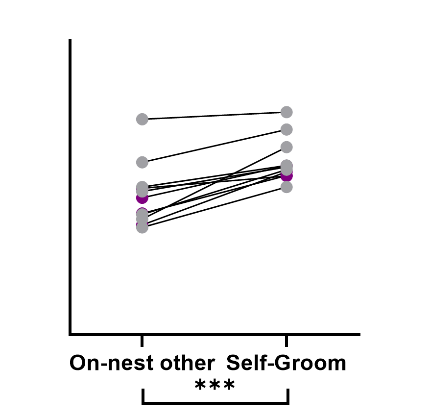

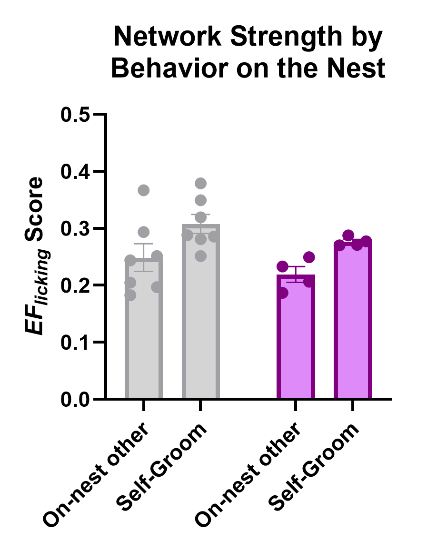

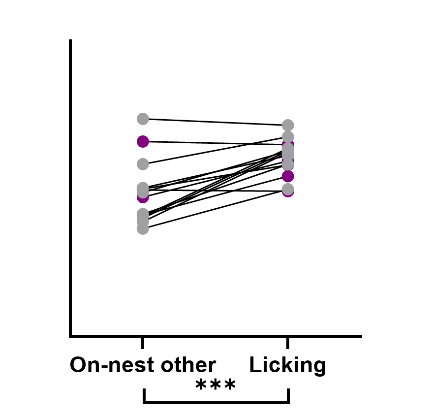

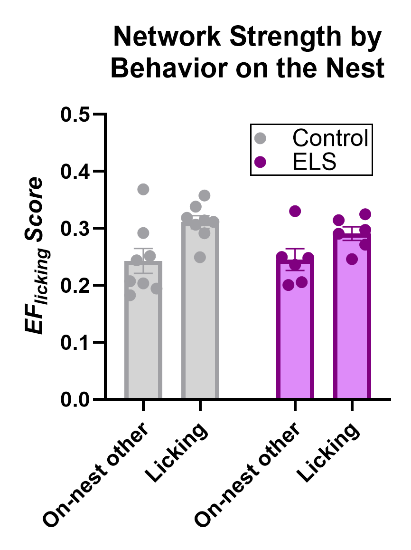

**b**

**a**

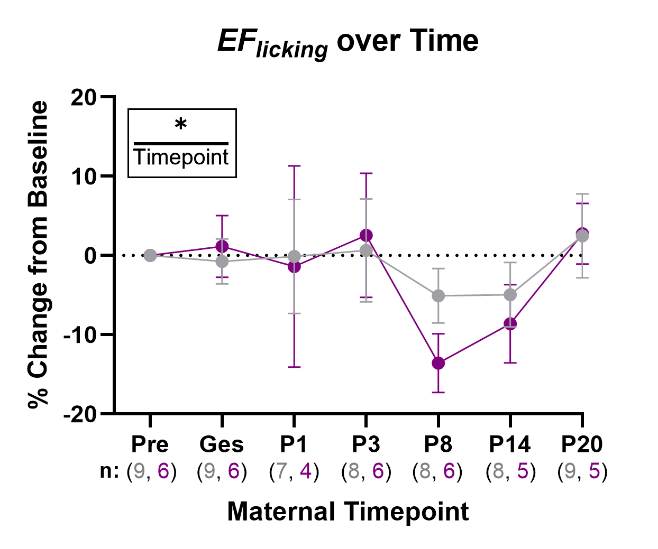

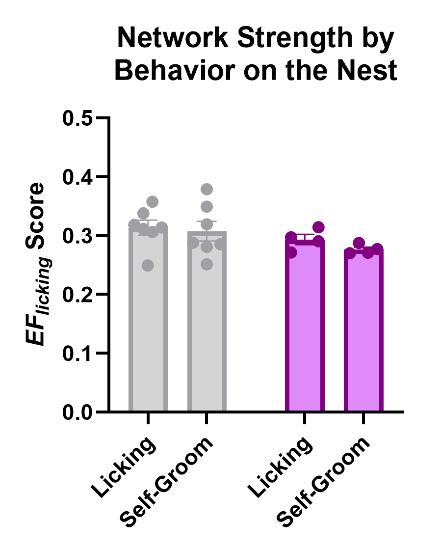

**d**

**c**

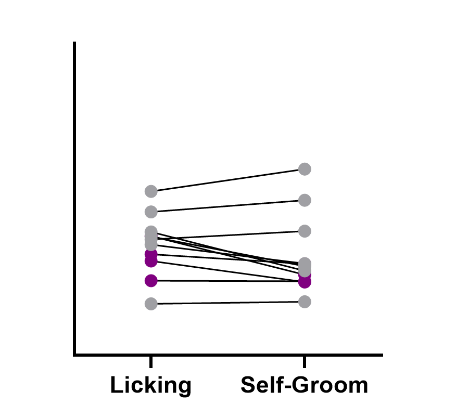

### Supplemental Figure S18: Licking (of pups), self-grooming behaviors, and the maternal trajectory for *EF_licking_*

The strength of *EF_licking_* during **a)** licking/grooming of pups by dams vs. all other behaviors on the nest (ANOVA, behavior effect, *P* = 0.0004, *n* = 6-8), **b)** during self-grooming behavior by dams vs. all other behaviors on the nest (ANOVA, behavior effect, *P* = 0.0005, *n* = 4-7), and **c)** licking/grooming of pups by dams vs. self-grooming by dams on the nest at P3. Both individual and paired representations of the data are shown. **d)** The strength of *EF_licking_* across observation timepoints calculated as the percent change from preconception baseline (REML, timepoint effect, *P* = 0.0420). The group specific *n* can be found below each respective observation timepoint. Multiple comparisons of individual timepoints can be found in Table S12.

Data are mean ± sem. **P* < 0.05, ****P* < 0.001. Statistics were performed using the two-way RM ANOVA **(a-c)** or two-way RM REML **(d)**. All statistical analyses of mean *EF* scores were performed on fourth root transformed data. This was done to help stabilize variance and make the data more normally distributed. For the normalized *EF* trajectory data**,** the transformation was performed prior to calculating the percent change from baseline.

*EF* = *Electome Factor*, Pre = preconception, Ges = gestation, P = postpartum day, RM = repeated measures, ANOVA = analysis of variance, REML = restricted maximum likelihood, FDR = false discovery rate

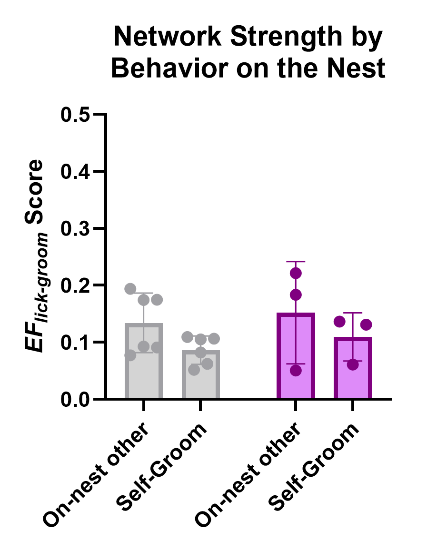

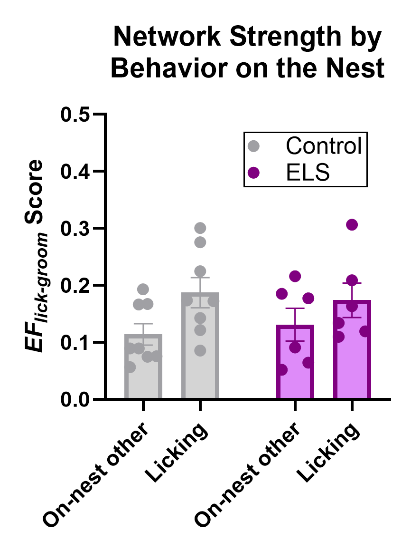

**a**

**b**

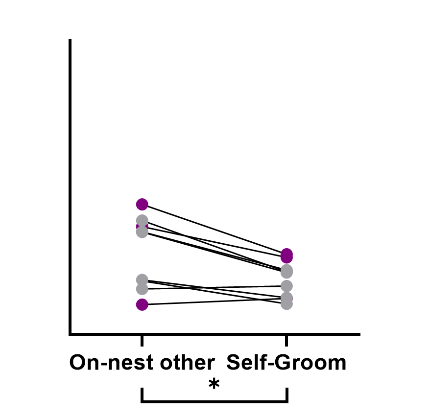

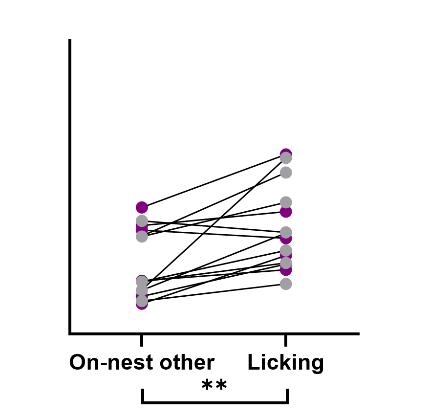

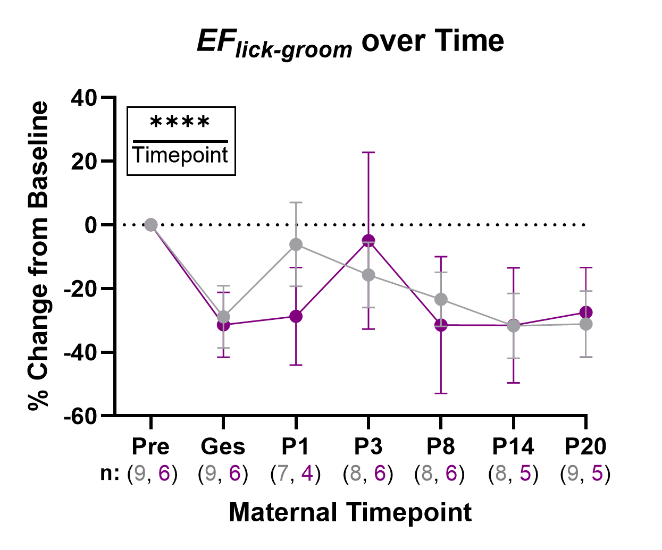

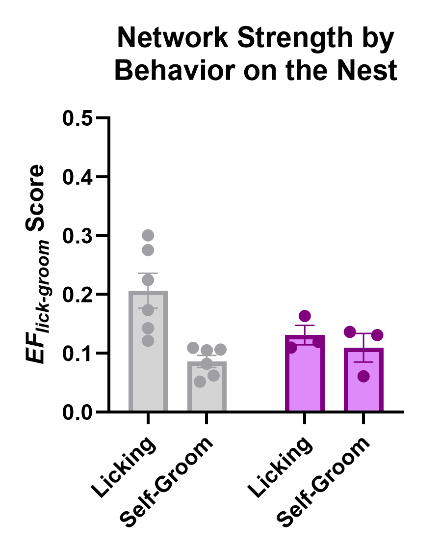

**c**

**d**

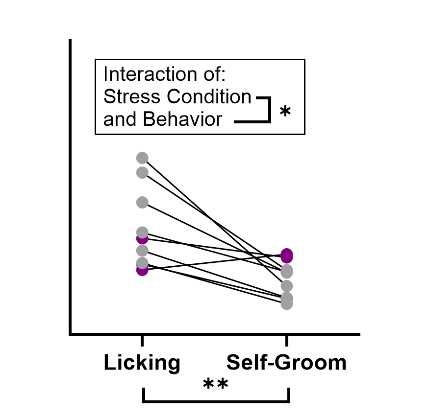

### Supplemental Figure S19: Licking (of pups), self-grooming behaviors, and the maternal trajectory for *EF_lick-groom_*

The strength of *EF_lick-groom_* during **a)** licking/grooming of pups by dams vs. all other behaviors on the nest (ANOVA, behavior effect, *P* = 0.0022, *n* = 6-8), **b)** during self-grooming behavior by dams vs. all other behaviors on the nest (ANOVA, behavior effect, *P* = 0.0130, *n* = 3-6), **c)** and during licking vs. self-grooming behaviors on the nest (ANOVA, behavior effect, *P* = 0.0020, ELS x behavior interaction, *P* = 0.0270, *n* = 3-6) at P3. Animals that spent less than 30-seconds self-grooming on the nest were excluded from the analysis. **d)** The strength of *EF_lick-groom_* across observation timepoints calculated as the percent change from preconception baseline (REML, timepoint effect, *P* = 1.87E-05). The group specific *n* can be found below each respective timepoint. Multiple comparisons of individual timepoints can be found in Table S13.

Data are mean ± sem. **P* < 0.05, ***P* < 0.01, *****P* < 0.0001. Statistics were performed using the two-way RM ANOVA **(a-c)** or REML **(d)**. All statistical analyses of mean *EF* scores were performed on fourth root transformed data. For the normalized *EF* trajectory data**,** the transformation was performed prior to calculating the percent change from baseline.

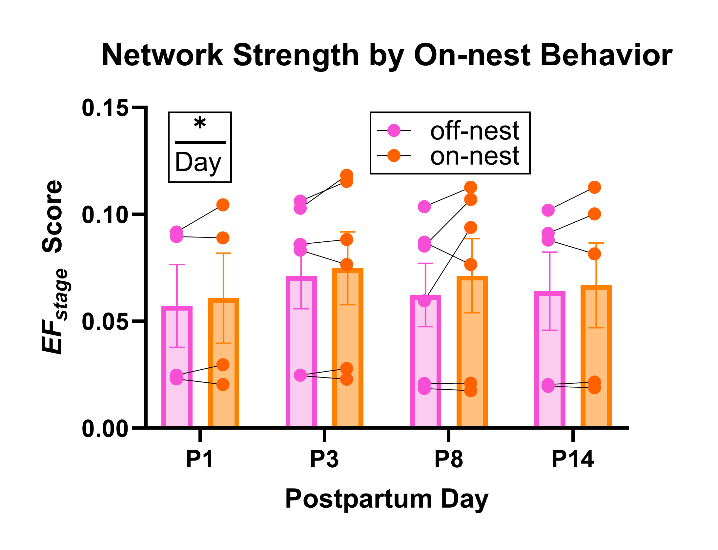

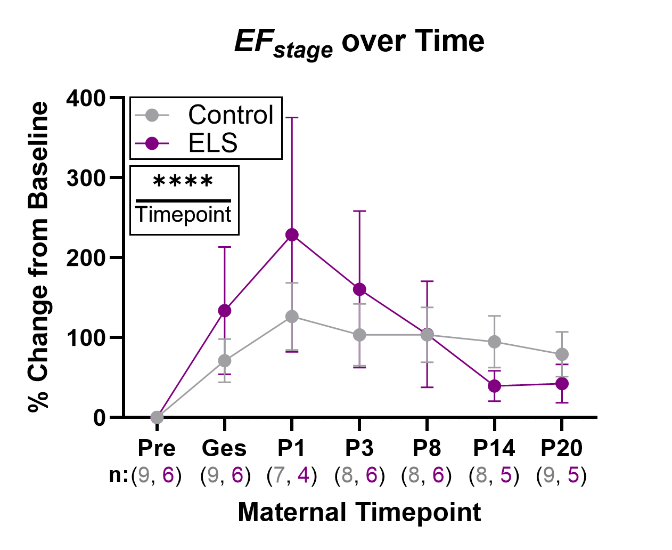

**b**

**a**

### Supplemental Figure S20: ELS-exposure does not impact *EF_stage_* performance or maternal trajectory

**a)** Mean *EF_stage_* score separated by “on” vs. “off-nest” behavior across observed maternal timepoints (REML, postpartum day effect, *P* = 0.0272, *n* = 4-6) for ELS-exposed dams. Multiple comparisons across postpartum days can be found in Table S14. **b)** Mean *EF_stage_* (REML, timepoint effect, *P* = 1.29E-08) scores across observed maternal timepoints normalized to the preconception baseline level for both control and ELS-exposed dams. Timepoint specific *n*/group noted beneath each stage. Multiple comparisons of individual timepoints can be found in Table S15.

Data are mean ± sem. **P* < 0.05, *****P* < 0.0001

Statistical analyses were completed using two-way RM REML. All statistical analyses of mean *EF* scores were performed on fourth root transformed data. For the normalized *EF* trajectory data**,** the transformation was performed prior to calculating the percent change from baseline. All data shown are non-transformed. If a significant effect of the repeated measure was found, a False Discovery Rate (FDR) correction for multiple comparisons was performed to specify which timepoints or measures were significantly different.

*EF* = *Electome Factor*, P = postpartum day, Pre = preconception, Ges = gestation, RM = repeated measures, REML = Restricted Maximum Likelihood, FDR = False Discovery Rate correction for multiple comparisons

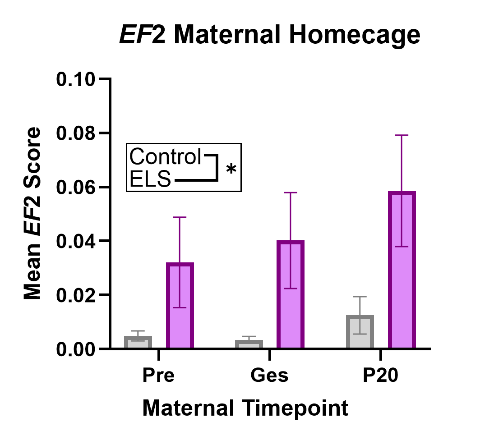

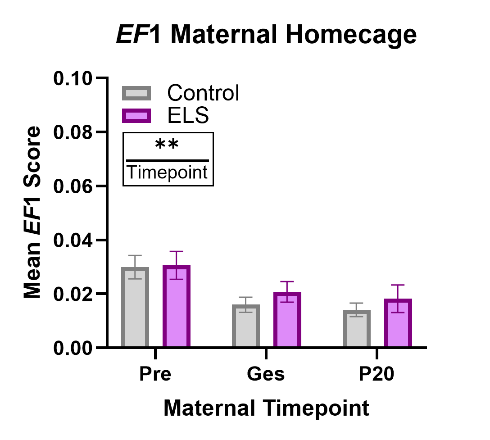

**b**

**a**

***EF*1**

***EF*2**

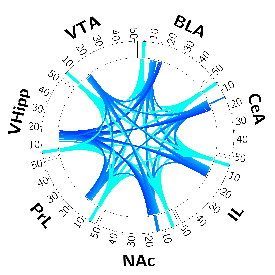

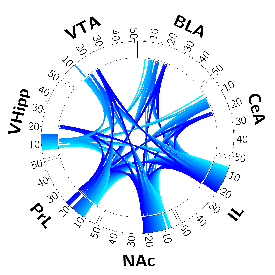

**c**

**d**

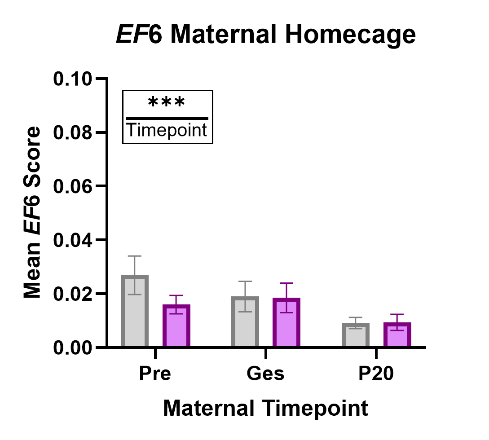

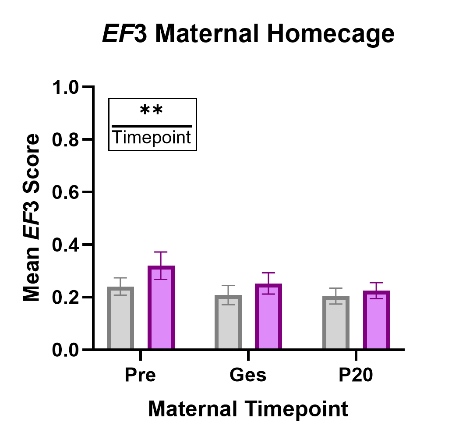

***EF*3**

***EF*6**

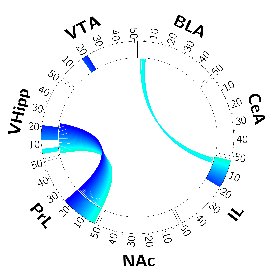

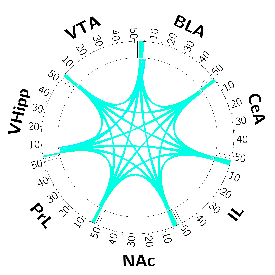

### Supplemental Figure S21: Stress-relevant *EF*s display dynamics in engagement over the maternal timecourse

**a)** Circos plot for the stress vulnerability predictive *EF*1, and mean *EF*1 scores across observed maternal-relevant timepoints (REML, timepoint effect, *P* = 0.0016, *n* = 13-20). Multiple comparisons of individual timepoints can be found in Table S16. **b)** Circos plot for the depressive-associated *EF*2, and mean *EF*2 scores across observed maternal-relevant timepoints (REML, ELS effect, *P* = 0.0397, *n* = 13-20). **c)** Circos plot for the depressive-associated *EF*3, and mean *EF*3 scores across observed maternal-relevant timepoints (REML, timepoint effect, *P* = 0.0045, *n* = 13-20). Multiple comparisons of individual timepoints can be found in Table S17. **d**) Circos plot for the stress-relevant *EF*6, and mean *EF*6 scores across observed maternal-relevant timepoints (REML, timepoint effect, *P* = 4.44E-04, *n* = 9-20). Multiple comparisons of individual timepoints can be found in Table S18.

Data shown are raw mean ± sem. **P*<0.05, ***P*<0.01, ****P*<0.001. Statistical analyses were performed using two-way RM REML (**a-d**) on fourth root transformed data. If a significant effect of the repeated measure was found, a False Discovery Rate (FDR) correction for multiple comparisons was performed.

PrL = Prelimbic cortex, IL = infralimbic cortex, NAc = nucleus accumbens, BLA = basolateral amygdala, CeA = central amygdala, MeA = medial amygdala, VHipp = ventral hippocampus, VTA = ventral tegmental area, *EF* = *Electome Factor*, ELS = early life stress, P = postpartum day, Pre = preconception, Ges = gestation, RM = repeated measures, REML = Restricted Maximum Likelihood, FDR = False Discovery Rate correction for multiple comparisons.

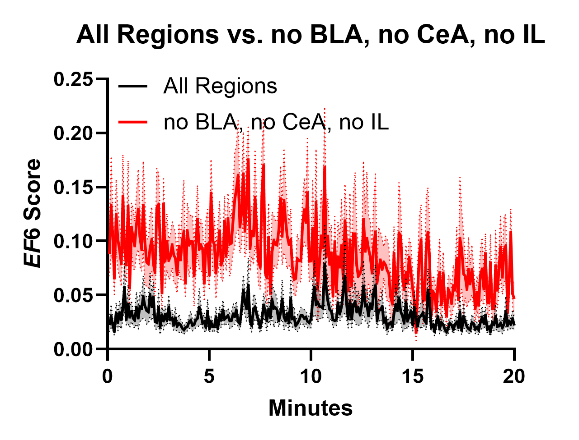

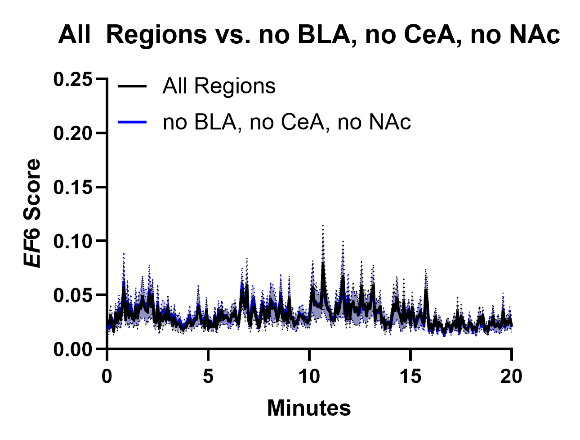

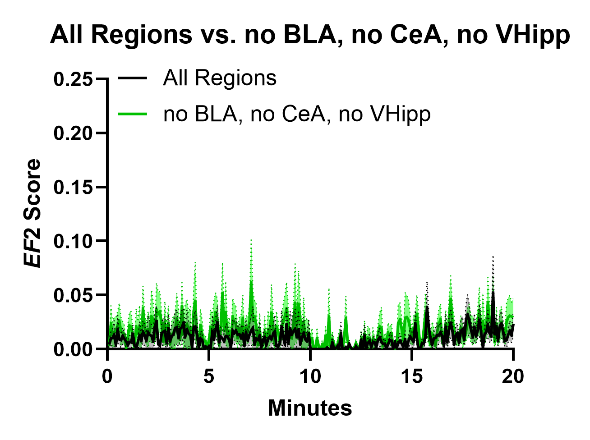

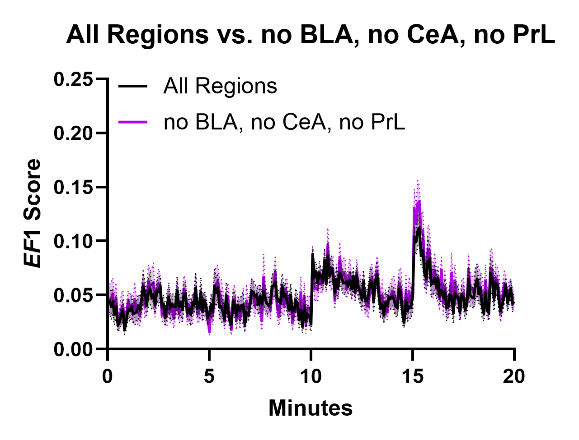

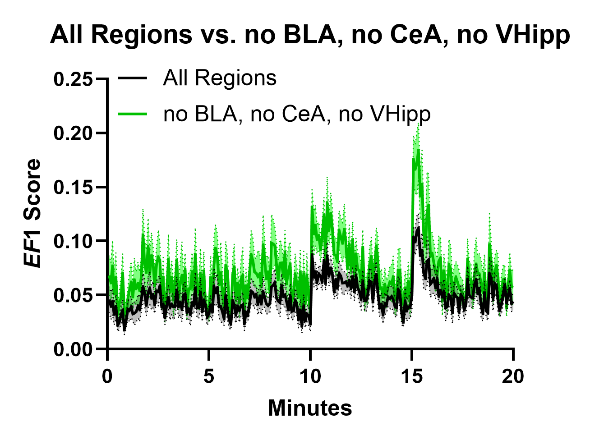

**b**

**a**

**c**

**d**

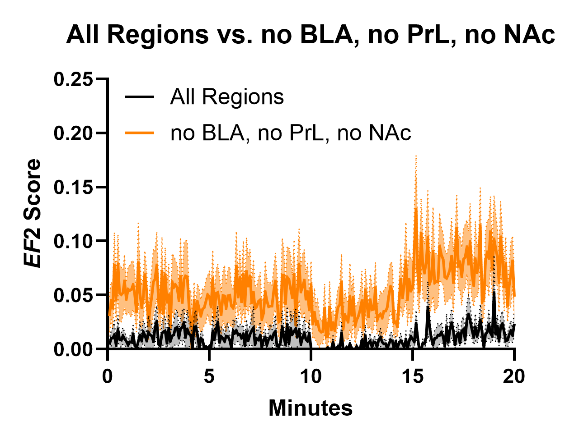

**f**

**e**

### Supplemental Figure S22: Testing the impact of missing LFP regions on stress *EF* score timecourses

Averaged preconception FIT stress *EF* score timecourse data when including all 7 regions (PrL, IL, NAc, BLA, CeA, VHipp, VTA), compared to parallel data missing a subset of three regions. An acceptable result for *EF*1 **(a)**, *EF*2 **(c)**, and *EF*6 **(e)** is shown alongside a corresponding unacceptable result **(b, d, f)** for the same three stress-relevant *EF*s (*n*=18). Combinations of missing regions that resulted in averaged timecourses diverging from the expected outcome were excluded from further analysis.

EF = *Electome Factor*, FIT = forced interaction test, PrL = Prelimbic cortex, IL = infralimbic cortex, NAc = nucleus accumbens, BLA = basolateral amygdala, CeA = central amygdala, VHipp = ventral hippocampus

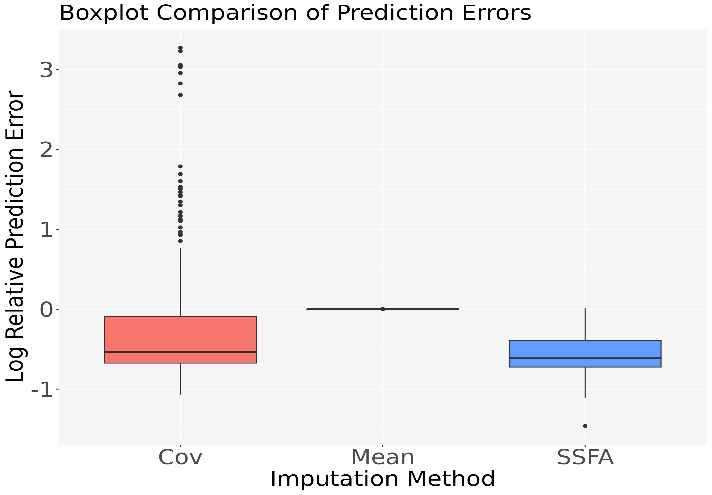

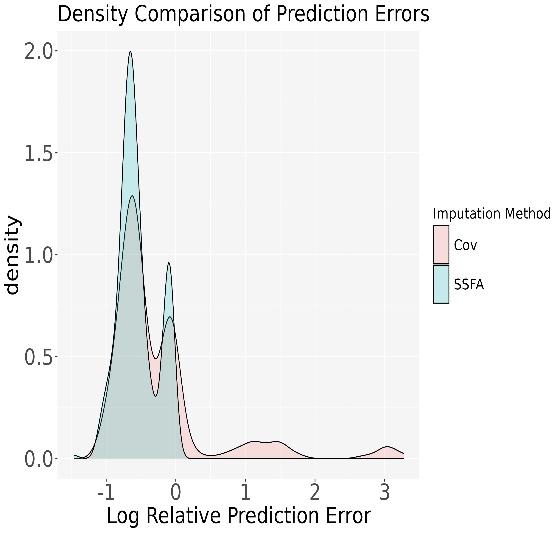

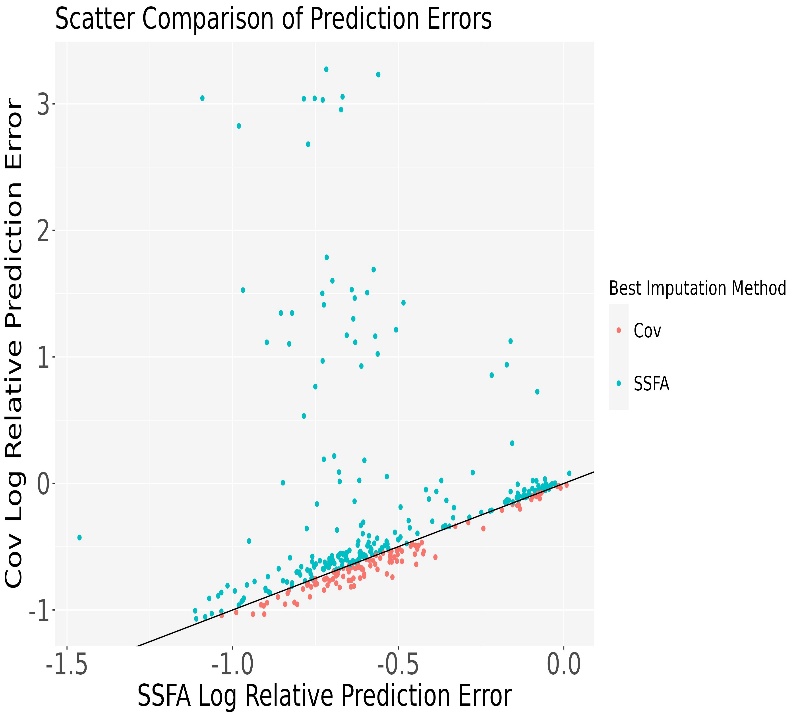

**a**

**b**

**c**

### Supplemental Figure S23: Testing the predictive performance of three data imputation methods

**a)** Distribution of the log relative prediction errors as boxplots for each method. **b)** Distribution of the log relative prediction errors for just the SSFA and traditional covariance estimate methods in terms of their densities. **c)** Plots comparing the log relative prediction errors for the SSFA and the traditional covariance estimate (Cov) methods against one another. Colored dots indicate which method had the smaller log relative prediction error. Each dot represents a cross-validation test; teal represents observations where SSFA had smaller RPE, and salmon represents observations where Cov had smaller RPE). The line indicates the point at which the two methods show equal levels of performance. The figure shows that SSFA generally outperforms Cov across observations, whereas Cov has a modest advantage for only a small subset. SSFA = simplified supervisory formula approach

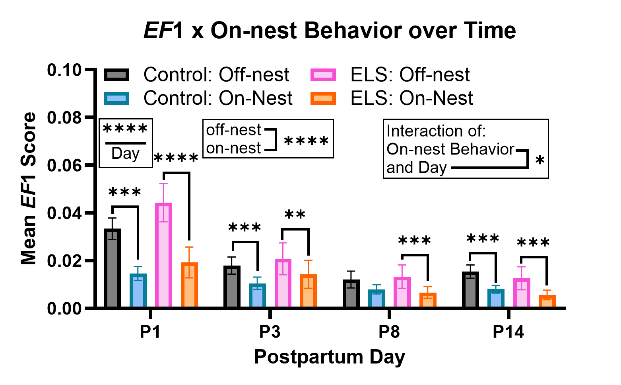

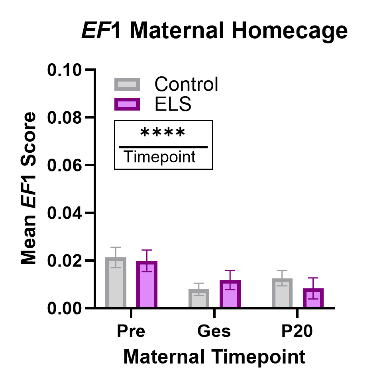

**c**

**b**

**a**

***EF*1**

***EF*6**

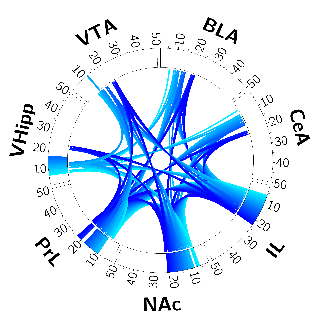

**e**

**f**

**d**

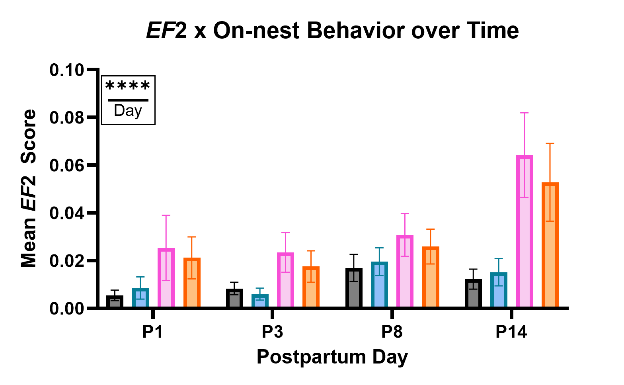

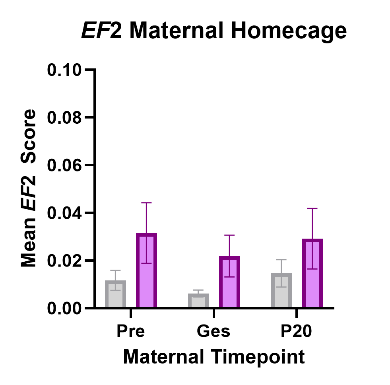

***EF*2**

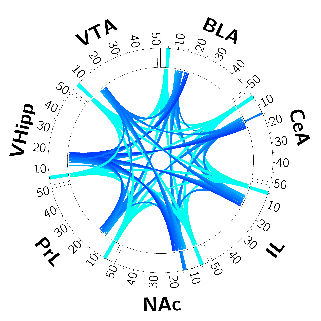

**i**

**h**

**g**

***EF*3**

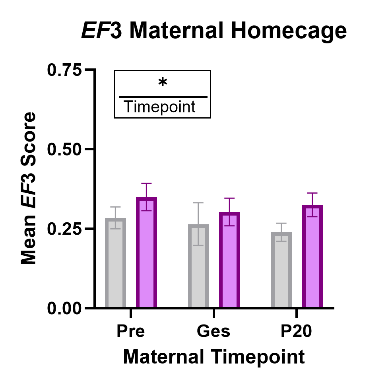

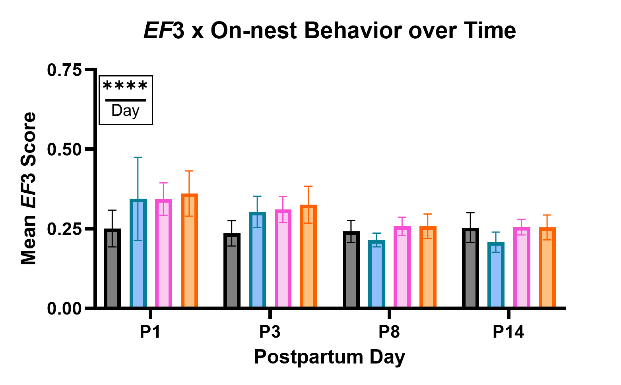

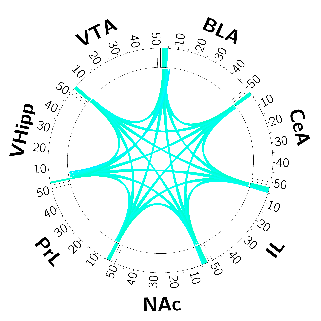

**l**

**k**

**j**

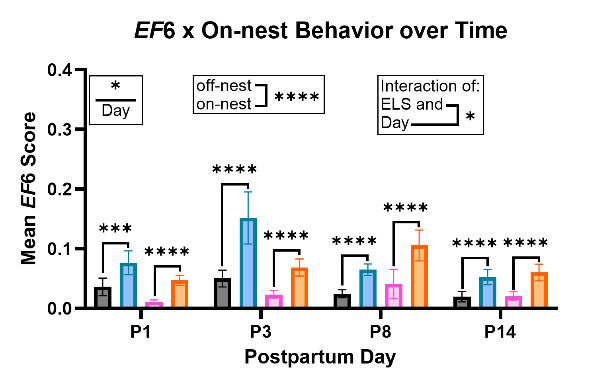

### Supplemental Figure S24: Stress-relevant *EF*s across maternal timepoints and postpartum behavior on the nest – imputation analysis

**a)** Circos plots presenting the network specific features (at 70% threshold) for the stress networks *EF*1, **d)** *EF*2, **g)** *EF*3, **j)** and *EF*6. **b)** Mean *EF*1, **e)** *EF*2, **h)** *EF*3, and **k)**, *EF*6 scores across observed maternal-relevant timepoints for control and ELS-exposed dams without pups present. **c)** Mean *EF*1, **f)** *EF*2, **i)** *EF*3, and **l)** *EF*6 scores across observed postpartum timepoints for control and ELS-exposed dams accounting for potential differences by engagement in “on” vs. “off-nest” behavior.

Data shown are raw mean ± sem. **p*<0.05, ***p*<0.01, ****p*<0.001, *****p*<0.0001

The data presented above show the same sets and comparisons as in Supplemental Fig. S21 and the main Figure 4. In the panels above, missing regional data that was imputed as described in the supplemental methods is included in the analysis. Statistics were performed on fourth root transformed data. The data presented in the figure are non-transformed.

Statistical analyses were performed using two-way RM REML (**b, e, h, k**) and three-way RM REML (**c, f, I, l**) on fourth root transformed data. If a significant effect of the repeated measure was found, a False Discovery Rate (FDR) correction for multiple comparisons was performed. For detailed discussion of the imputed dataset results and comparison to the “cleaned and complete” dataset, refer to the supplemental results and Tables S16-S25 and S27-S35. For *EF* and timepoint specific animal numbers, see Table S26.

PrL = Prelimbic cortex, IL = infralimbic cortex, NAc = nucleus accumbens, BLA = basolateral amygdala, CeA = central amygdala, MeA = medial amygdala, VHipp = ventral hippocampus, VTA = ventral tegmental area, *EF* = *Electome Factor*, Pre = preconception, Ges = gestation, P = postpartum day, ELS = early life stress, RM = repeated measures, REML = restricted maximum likelihood, FDR = False Discovery Rate correction for multiple comparisons

**b**

**a**

**d**

**c**

### Supplemental Figure S25: Correlations of time on nest with *EF_on-nest_* strength during postpartum behavior

Correlation plots for total percent time on the nest and *EF_on-nest_* score during behavior on the nest for control and ELS dams at **a)** P1, **b)** P3, **c)** P8 (significant positive correlation for control animals only at P8 (*P* = 0.0033) with a significant difference between regression line slopes (*P* = 0.0006) for control and ELS dams, and **d)** P14 (*n* = 4-8 animals/group).

Raw mean data are shown. ***P* < 0.01, ****P* < 0.001

Simple linear regression lines are displayed. Pearson’s correlations were performed and the slopes of the linear regression lines were compared for statistical difference. All statistical analyses of mean *EF* scores were performed on fourth root transformed data.

*EF* = *Electome Factor*, ELS = early life stress, P = postpartum day, *r* = Pearson’s correlation coefficient

**b**

**a**

**c**

**d**

### Supplemental Figure S26: Prior ELS-exposure does not robustly alter maternal behavior throughout most of postpartum

**a)** Illustration of a dam spending time on the nest. **b)** Percent total time “on-nest” for control and ELS-exposed dams across the observed postpartum timepoints. **c)** Mean number of visits to the nest per 1-hour of observation. One outlier, more than 7x the next highest mean value, was removed at P14. Nest visits of all durations were included in the analysis (REML, Day effect, *P* = 4.14E-11, *n*=11-19/group/stage). Multiple comparisons of individual timepoints can be found in Table S36. **d)** Mean duration of care visits to the nest during naturalistic maternal care observations at four postpartum timepoints. Only visits including observations of care behavior were included in the analysis (REML, Day effect, *P* = 4.18E-04; ELS x Day interaction, *P* = 0.0256, *n* =11-19/group/stage). Multiple comparisons of individual timepoints can be found in Table S37. Panel **a** was created in BioRender. Mitchell, SB. (2026) https://BioRender.com/x9540la.

Data shown are raw mean ± sem. **P* < 0.05, ***P < 0.001, *****P* < 0.0001

Statistical analyses were performed using two-way RM REML (**b-d**). All statistical analyses of mean *EF* scores were performed on fourth root transformed data. If a significant effect of the repeated measure was found, a False Discovery Rate (FDR) correction for multiple comparisons was performed to specify which timepoints/days were significantly different.

ELS = early life stress, P = postpartum day, RM = repeated measures, REML = Restricted Maximum Likelihood, FDR = False Discovery Rate correction for multiple comparisons.

**a**

**d**

**c**

**b**

**f**

**e**

**g**

**h**

### Supplemental Figure S27: ELS altered patterning of maternal care behavior, but not dam self-care at P3

**A)** Illustration of a mouse dam on the nest with pups. **b)** Number of care visits to the nest (*n* = 16-19/group). **c)** Mean duration of dam care visits (mins) to the nest (*P* = 0.0058, *n* = 16-19/group). **d)** Percentage of time on the nest that was spent nursing by dams (*P* = 0.0495, *n* = 16-18/group). **e)** Time (s) spent licking/grooming pups (*n* = 16-18/group). **f)** Time (s) by dams spent self-grooming (*n* = 16-18/group). **g)** Difference in time dams spent grooming self as compared to pups and (*n* = 16-18/group). **h)** Time (s) dams spent eating (*n* = 16-18/group) over a 3-h observation at P3. The illustration in Panel **a** was created using DALL – E via openai.com.

Data shown are raw mean ± sem. **P* < 0.05, ***P* < 0.01.

Statistics were performed using the unpaired t-test or non-parametric Mann Whitney test.

P = postpartum day, ELS = early life stress, mins = minutes, s = seconds, h = hour(s)

**h**

**g**

**f**

**e**

**d**

**c**

**b**

**a**

### Supplemental Figure S28: Blinded behavioral frequency scoring produced comparable results to the continuous unblinded maternal scoring approach

The percent of behavioral bins engaged in sub-typed behaviors at P3. Frequency of behavioral bins = once every 4-mins. Length of observation = 3-h. Recorded dam behaviors include **a)** nursing of pups, **b)** nest building, **c)** licking/grooming of pups, **d)** retrieval of pups, **e)** roaming of cage, **f)** digging in bedding/nesting material, **g)** eating, **h)** and licking/grooming of self (*n* = 16-19/group).

Data shown are raw mean ± sem.

Statistics were performed using the unpaired t-test or non-parametric Mann Whitney test.

P = postpartum day, ELS = early life stress, mins = minutes, h = hour(s)

**b**

**a**

**d**

**c**

### Supplemental Figure S29: Correlations of time on nest with *EF*1 network strength

Correlation plots for total percent time on the nest and the stress-relevant *EF*1 score for control and ELS-exposed dams at **a)** P1 (significant correlation for ELS animals only, *P* = 0.0072, *n* = 9-13), **b)** P3 (significant correlation for control animals only, *P* = 0.0057, *n* = 15-17), **c)** P8 (*n* = 15-16), and **d)** P14 (*n* = 13-15).

Raw mean data are shown. ***P* < 0.01

Statistics were performed on fourth root transformed data. Simple linear regression lines are displayed. Pearson’s correlations were performed and the slopes of the linear regression lines were compared for statistical difference.

*EF* = *Electome Factor*, P = postpartum day, ELS = early life stress, *r* = Pearson’s correlation coefficient

**a**

**c**

**b**

### Supplemental Figure S30: *EF*1 and the open field pup retrieval task

**a)** Schematic presentation of the open field pup retrieval task and the scoring method for pup grouping completed at P4. **b)** Mean stress-relevant *EF*1 scores for control and ELS-exposed dams during the two stages of the open field pup retrieval task (*n* = 5-6 animals/group). **c)** Open field pup retrieval scores for control and ELS-exposed dams.

Panel **a** was created in BioRender. Mitchell, SB. (2026) https://BioRender.com/nvdzqj6.

Data shown are raw mean ± sem.

Statistics were performed on fourth root transformed data using a two-way RM ANOVA **(b)**.

*EF* = *Electome Factor*, ELS = early life stress, RM = repeated measures, ANOVA = analysis of variance

**b**

**a**

**c**

### Supplemental Figure S31: *EF*6 during preconception and P4 pup retrieval tasks

**a)** The mean baseline *EF*6 score for control and ELS-exposed animals prior to the start of the pup retrieval screen (*n* = 5-7/group). **b)** Mean *EF*6 score for control and ELS-exposed animals during baseline, retrieval trials, and active retrieval behavior (AVOVA, task stage effect, *P* = 0.0053, *n* = 4/group, Table S39). The “baseline” stage refers to the habituation period prior to the start of the retrieval trials. The “trials” stage refers to the period of retrieval trials (10 x 1 pup removed from nest) over which no active retrieval behavior took place. The “retrieval” stage refers to periods of active retrieval behavior. **c)** Mean stress-relevant *EF*6 scores for control and ELS-exposed dams during the two stages of the open field pup retrieval task (*n* = 4-5/group).

Data shown are raw mean ± sem. ***P* < 0.01. Statistics were performed on fourth root transformed data using a two-way RM ANOVA **(b-c)**. *EF* = *Electome Factor*, ELS = early life stress, RM = repeated measures, ANOVA = analysis of variance

**b**

**a**

**d**

**c**

**f**

**e**

### Supplemental Figure S32: Novel maternal network strength during subtyped behaviors

**a)** Mean *EF_on-nest_* scores and **b)** mean *EF_stage_* scores for control and ELS-exposed dams during three parts of a pup retrieval trial at P4 (*n* = 4 animals/group). The “baseline” stage refers to the habituation period prior to the start of the retrieval trials. The “trials” stage refers to the period of retrieval trials (10 x 1 pup removed from nest) over which no active retrieval behavior took place. The “retrieval” stage refers to periods of active retrieval behavior. **c)** The strength of *EF_on-nest_* during licking/grooming of pups by dams vs. all other behaviors (two-way RM ANOVA, behavior effect, *P* = 0.0001, *n* = 6-8) at P3. **d)** The strength of *EF_stage_* during licking/grooming of pups by dams vs. all other behaviors (two-way RM ANOVA, behavior effect, *P* = 0.0015, *n* = 6-8) at P3. **e)** The strength of *EF_on-nest_* during self-grooming behavior by dams vs. all other behaviors at P3 (*n* = 6-8/group). **f)** The strength of *EF_stage_* during self-grooming behavior by dams vs. all other behaviors at P3 (*n* = 6-8/group). Both individual and paired representations of the data are shown.

Data shown are raw mean ± sem. ***P* < 0.01, ****P* < 0.001

Statistics were performed on fourth root transformed data using two-way RM ANOVAs (**a-f**).

*EF* = *Electome Factor*, P = postpartum day, RM = repeated measures, ANOVA = Analysis of Variance

**FIT Timecourse**

**a**

**b**

**Mean *EF*1 Score**

**c**

**d**

**FIT Timecourse**

**Mean *EF*2 Score**

### Supplemental Figure S33: Stress *EFs* and a negative affect task before and after the maternal transition - imputation analysis

**a)** *EF*1 and **c)** *EF*2 summary data for all stages of the FIT for both control and ELS-exposed dams at two timepoints: preconception and late postpartum. **b)** Mean *EF*1 and **d)** *EF*2 scores across the FIT timecourse for all comparisons of interest. The dotted lines denote changes in the stage of the FIT.

Data shown are raw mean ± sem. **P* < 0.05, ***P* < 0.01, ****P* < 0.001, *****P* < 0.0001

The data presented show the same sets and comparisons as in the main Figure 6. In the panels above, missing regional data that was imputed as described in the supplemental methods is included in the analysis.

Statistics were performed on fourth root transformed data. For panels **a**, and **c**, main effects and interactions were assessed using a three-way RM mixed-effects model (REML). Differences between specific task stages were evaluated and multiple comparisons were accounted for using False Discovery Rate (FDR) correction. For detailed discussion of the imputed dataset results and comparison to the “cleaned and complete” dataset, refer to the supplemental results and Tables S40-S47. For *EF* and timepoint specific animal numbers, see Table S26.

*EF* = *Electome Factor*, Pre = preconception, Post = postpartum, FIT = forced interaction test, RM = repeated measures, REML = restricted maximum likelihood, FDR = False Discovery Rate correction for multiple comparisons

### Table S1 – Surgical coordinates for targeted electrode implantation

| **Regional Bundle** | **AP** | **ML** | **DV (brain)** |
| --- | --- | --- | --- |
| PFC (PrL) | 1.36 | 0.00 ± 0.375 | 1.94 |
| PFC (IL) | 1.86 | 0.00 ± 0.375 | 2.19 |
| NAc | 1.56 | -1.37 | 3.41 |
| AMY (BLA) | -1.665 | -2.93 | 3.8 |
| AMY (CeA) | -1.415 | -2.43 | 3.55 |
| AMY (MeA) | -1.665 | -1.93 | 4.8 |
| VTA | -3.22 | -0.49 | 4.14 |
| VHipp | -3.61 | 2.93 | 3.41 |

PFC = prefrontal cortex, PrL = prelimbic cortex, IL = infralimbic cortex, NAc = nucleus accumbens, AMY = amygdala, BLA = basolateral amygdala, CeA = central amygdala, MeA = medial amygdala, VTA = ventral tegmental area, VHipp = ventral hippocampus

Coordinates provided indicate the central point of each bundle measured from Bregma. The DV measurement is taken from the surface of the brain.

### Table S2 – *EF_on-nest_* scores by “off-nest” and “on-nest” behavior across postpartum observation days [Controls]

| **Contrast** | **Postpartum Day** | **Discovery** | **FDR Adj. P-value** |
| --- | --- | --- | --- |
| “off-nest” vs. “on-nest” | **1** | **Yes** | **1.84E-06** |
| “off-nest” vs. “on-nest” | **3** | **Yes** | **7.52E-07** |
| “off-nest” vs. “on-nest” | **8** | **Yes** | **7.52E-07** |
| “off-nest” vs. “on-nest” | **14** | **Yes** | **1.84E-06** |

### Table S3 – *EF_on-nest_* scores by on-nest behavior and “on-nest” subtyped behavior (nursing) at P3 [Controls]

| **Contrast** | **Discovery** | **FDR Adj. P-value** |
| --- | --- | --- |
| **“off-nest” vs. Nursing** | **Yes** | **0.0002** |
| “off-nest” vs. “on-nest” other | No | 0.3024 |
| **Nursing vs. “on-nest” other** | **Yes** | **0.0001** |

### Table S4 – *EF_on-nest_* percent change from baseline: Multiple comparisons by maternal timepoint [Controls]

| **Contrast** | **Discovery** | **FDR Adj. P-value** |
| --- | --- | --- |
| Pre vs. Ges | No | 0.2330 |
| **Pre vs. P1** | **Yes** | **0.0043** |
| **Pre vs. P3** | **Yes** | **0.0012** |
| **Pre vs. P8** | **Yes** | **0.0006** |
| **Pre vs. P14** | **Yes** | **0.0022** |
| Pre vs. P20 | No | 0.3061 |
| Ges vs. P1 | No | 0.0552 |
| **Ges vs. P3** | **Yes** | **0.0194** |
| **Ges vs. P8** | **Yes** | **0.0082** |
| **Ges vs. P14** | **Yes** | **0.0358** |
| Ges vs. P20 | No | 0.8774 |
| P1 vs. P3 | No | 0.7945 |
| P1 vs. P8 | No | 0.5500 |
| P1 vs. P14 | No | 0.8774 |
| **P1 vs. P20** | **Yes** | **0.0392** |
| P3 vs. P8 | No | 0.7956 |
| P3 vs. P14 | No | 0.8261 |
| **P3 vs. P20** | **Yes** | **0.0133** |
| P8 vs. P14 | No | 0.6107 |
| **P8 vs. P20** | **Yes** | **0.0057** |
| **P14 vs. P20** | **Yes** | **0.0248** |

Pre = preconception, Ges = gestation, P = postpartum day

### Table S5 – *EF_lick-groom_* percent change from baseline: Multiple comparisons by maternal timepoint [Controls]

| **Contrast** | **Discovery** | **FDR Adj. P-value** |
| --- | --- | --- |
| **Pre vs. Ges** | **Yes** | **0.0045** |
| Pre vs. P1 | No | 0.9208 |
| Pre vs. P3 | No | 0.1518 |
| **Pre vs. P8** | **Yes** | **0.0297** |
| **Pre vs. P14** | **Yes** | **0.0045** |
| **Pre vs. P20** | **Yes** | **0.0045** |
| **Ges vs. P1** | **Yes** | **0.0089** |
| Ges vs. P3 | No | 0.1294 |
| Ges vs. P8 | No | 0.4432 |
| Ges vs. P14 | No | 0.9568 |
| Ges vs. P20 | No | 0.9452 |
| P1 vs. P3 | No | 0.2784 |
| P1 vs. P8 | No | 0.0727 |
| **P1 vs. P14** | **Yes** | **0.0089** |
| **P1 vs. P20** | **Yes** | **0.0081** |
| P3 vs. P8 | No | 0.4432 |
| P3 vs. P14 | No | 0.1294 |
| P3 vs. P20 | No | 0.1085 |
| P8 vs. P14 | No | 0.4432 |
| P8 vs. P20 | No | 0.4136 |
| P14 vs. P20 | No | 0.9486 |

Pre = preconception, Ges = gestation, P = postpartum day

### Table S6 – *EF_stage_* percent change from baseline: Multiple comparisons by maternal timepoint [Controls]

| **Contrast** | **Discovery** | **FDR Adj. P-value** |
| --- | --- | --- |
| **Pre vs. Ges** | **Yes** | **9.29E-05** |
| **Pre vs. P1** | **Yes** | **3.75E-07** |
| **Pre vs. P3** | **Yes** | **1.92E-05** |
| **Pre vs. P8** | **Yes** | **1.92E-05** |
| **Pre vs. P14** | **Yes** | **4.72E-05** |
| **Pre vs. P20** | **Yes** | **0.0003** |
| Ges vs. P1 | No | 0.1520 |
| Ges vs. P3 | No | 0.4177 |
| Ges vs. P8 | No | 0.4177 |
| Ges vs. P14 | No | 0.5637 |
| Ges vs. P20 | No | 0.9473 |
| P1 vs. P3 | No | 0.5637 |
| P1 vs. P8 | No | 0.6066 |
| P1 vs. P14 | No | 0.4231 |
| P1 vs. P20 | No | 0.1520 |
| P3 vs. P8 | No | 0.9473 |
| P3 vs. P14 | No | 0.8024 |
| P3 vs. P20 | No | 0.4232 |
| P8 vs. P14 | No | 0.7558 |
| P8 vs. P20 | No | 0.4177 |
| P14 vs. P20 | No | 0.5637 |

Pre = preconception, Ges = gestation, P = postpartum day

### Table S7 – *EF_stage_* scores by “off-nest” and “on-nest” behavior across postpartum observation days [Controls]

| **Contrast** | **Postpartum Day** | **Discovery** | **FDR Adj. P-value** |
| --- | --- | --- | --- |
| “off-nest” vs. “on-nest” | 1 | No | 0.2417 |
| “off-nest” vs. “on-nest” | 3 | No | 0.2417 |
| “off-nest” vs. “on-nest” | 8 | No | 0.2417 |
| “off-nest” vs. “on-nest” | 14 | No | 0.3931 |

### Table S8 – Homecage *EF*1 scores: Multiple comparisons by maternal timepoint [Controls]

| **Contrast** | **Discovery** | **FDR Adj. P-value** |
| --- | --- | --- |
| **Pre vs. Ges** | **Yes** | **0.0414** |
| **Pre vs. P20** | **Yes** | **0.0414** |
| Ges vs. P20 | No | 0.8441 |

Pre = preconception, Ges = gestation, P = postpartum day

### Table S9 – Homecage *EF*6 scores: Multiple comparisons by maternal timepoint [Controls]

| **Contrast** | **Discovery** | **FDR Adj. P-value** |
| --- | --- | --- |
| Pre vs. Ges | No | 0.1684 |
| **Pre vs. P20** | **Yes** | **0.0242** |
| Ges vs. P20 | No | 0.1684 |

Pre = preconception, Ges = gestation, P = postpartum day

### Table S10 – *EF_on-nest_* scores by “off-nest” and “on-nest” behavior across postpartum observation days [ELS]

| **Contrast** | **Postpartum Day** | **Discovery** | **FDR Adj. P-value** |
| --- | --- | --- | --- |
| “off-nest” vs. “on-nest” | 1 | No | 0.1247 |
| “off-nest” vs. “on-nest” | 3 | No | 0.1632 |
| “off-nest” vs. “on-nest” | 8 | No | 0.0657 |
| “off-nest” vs. “on-nest” | 14 | No | 0.0808 |

### Table S11 – *EF_on-nest_* percent change from baseline: Multiple comparisons by maternal timepoint

| **Contrast** | **Discovery** | **FDR Adj. P-value** |
| --- | --- | --- |
| Pre vs. Ges | No | 0.4488 |
| **Pre vs. P1** | **Yes** | **0.0078** |
| **Pre vs. P3** | **Yes** | **0.0078** |
| **Pre vs. P8** | **Yes** | **0.0014** |
| **Pre vs. P14** | **Yes** | **0.0137** |
| Pre vs. P20 | No | 0.5584 |
| Ges vs. P1 | No | 0.0608 |
| Ges vs. P3 | No | 0.0608 |
| **Ges vs. P8** | **Yes** | **0.0084** |
| Ges vs. P14 | No | 0.1101 |
| Ges vs. P20 | No | 0.1444 |
| P1 vs. P3 | No | 0.8213 |
| P1 vs. P8 | No | 0.6779 |
| P1 vs. P14 | No | 0.7405 |
| **P1 vs. P20** | **Yes** | **0.0022** |
| P3 vs. P8 | No | 0.5168 |
| P3 vs. P14 | No | 0.8213 |
| **P3 vs. P20** | **Yes** | **0.0020** |
| P8 vs. P14 | No | 0.4412 |
| **P8 vs. P20** | **Yes** | **3.20E-04** |
| **P14 vs. P20** | **Yes** | **0.0033** |

Pre = preconception, Ges = gestation, P = postpartum day

### Table S12 – *EF_licking_* percent change from baseline: Multiple comparisons by maternal timepoint

| **Contrast** | **Discovery** | **FDR Adj. P-value** |
| --- | --- | --- |
| Pre vs. Ges | No | 0.9897 |
| Pre vs. P1 | No | 0.9897 |
| Pre vs. P3 | No | 0.7563 |
| Pre vs. P8 | No | 0.0825 |
| Pre vs. P14 | No | 0.2997 |
| Pre vs. P20 | No | 0.5584 |
| Ges vs. P1 | No | 0.9897 |
| Ges vs. P3 | No | 0.7563 |
| Ges vs. P8 | No | 0.0825 |
| Ges vs. P14 | No | 0.2997 |
| Ges vs. P20 | No | 0.5584 |
| P1 vs. P3 | No | 0.7563 |
| P1 vs. P8 | No | 0.1042 |
| P1 vs. P14 | No | 0.3184 |
| P1 vs. P20 | No | 0.6085 |
| **P3 vs. P8** | **Yes** | **0.0442** |
| P3 vs. P14 | No | 0.1181 |
| P3 vs. P20 | No | 0.7563 |
| P8 vs. P14 | No | 0.6698 |
| **P8 vs. P20** | **Yes** | **0.0277** |
| P14 vs. P20 | No | 0.0825 |

Pre = preconception, Ges = gestation, P = postpartum day

### Table S13 – *EF_lick-groom_* percent change from baseline: Multiple comparisons by maternal timepoint

| **Contrast** | **Discovery** | **FDR Adj. P-value** |
| --- | --- | --- |
| Pre vs. Ges | **Yes** | **1.08E-04** |
| Pre vs. P1 | No | 0.1258 |
| Pre vs. P3 | **Yes** | **0.0489** |
| Pre vs. P8 | **Yes** | **1.08E-04** |
| Pre vs. P14 | **Yes** | **8.81E-05** |
| Pre vs. P20 | **Yes** | **1.08E-04** |
| Ges vs. P1 | **Yes** | **0.0425** |
| Ges vs. P3 | **Yes** | **0.0454** |
| Ges vs. P8 | No | 0.8635 |
| Ges vs. P14 | No | 0.7955 |
| Ges vs. P20 | No | 0.9184 |
| P1 vs. P3 | No | 0.8295 |
| P1 vs. P8 | **Yes** | **0.04912** |
| P1 vs. P14 | **Yes** | **0.0252** |
| P1 vs. P20 | **Yes** | **0.0425** |
| P3 vs. P8 | No | 0.0626 |
| P3 vs. P14 | **Yes** | **0.0252** |
| P3 vs. P20 | **Yes** | **0.0454** |
| P8 vs. P14 | No | 0.6548 |
| P8 vs. P20 | No | 0.8295 |
| P14 vs. P20 | No | 0.8295 |

Pre = preconception, Ges = gestation, P = postpartum day

### Table S14 – *EF_stage_* scores by on-nest behavior: Multiple comparisons by postpartum observation day [ELS]

| **Contrast** | **Discovery** | **FDR Adj. P-value** |
| --- | --- | --- |
| P1 vs. P3 | No | 0.6849 |
| P1 vs. P8 | No | 0.0580 |
| P1 vs. P14 | No | 0.0951 |
| P3 vs. P8 | No | 0.0580 |
| P3 vs. P14 | No | 0.0981 |
| P8 vs. P14 | No | 0.6849 |

P = postpartum day

### Table S15 – *EF_stage_* percent change from baseline: Multiple comparisons by maternal timepoint

| **Contrast** | **Discovery** | **FDR Adj. P-value** |
| --- | --- | --- |
| **Pre vs. Ges** | **Yes** | **3.44E-06** |
| **Pre vs. P1** | **Yes** | **5.23E-08** |
| **Pre vs. P3** | **Yes** | **5.23E-08** |
| **Pre vs. P8** | **Yes** | **2.03E-06** |
| **Pre vs. P14** | **Yes** | **1.11E-05** |
| **Pre vs. P20** | **Yes** | **1.63E-04** |
| Ges vs. P1 | No | 0.1437 |
| Ges vs. P3 | No | 0.3253 |
| Ges vs. P8 | No | 0.8093 |
| Ges vs. P14 | No | 0.9247 |
| Ges vs. P20 | No | 0.4567 |
| P1 vs. P3 | No | 0.5881 |
| P1 vs. P8 | No | 0.2293 |
| P1 vs. P14 | No | 0.1437 |
| **P1 vs. P20** | **Yes** | **0.0307** |
| P3 vs. P8 | No | 0.4565 |
| P3 vs. P14 | No | 0.3253 |
| P3 vs. P20 | No | 0.0861 |
| P8 vs. P14 | No | 0.7835 |
| P8 vs. P20 | No | 0.3434 |
| P14 vs. P20 | No | 0.5138 |

Pre = preconception, Ges = gestation, P = postpartum day

### Table S16 – *EF*1 maternal homecage: Multiple comparisons by maternal timepoint

| **Contrast** | **Discovery** | **FDR Adj. P-value** |
| --- | --- | --- |
| **Pre vs. Ges** | **Yes** | **0.0116** |
| **Pre vs. P20** | **Yes** | **0.0023** |
| Ges vs. P20 | No | 0.3564 |

Pre = preconception, Ges = gestation, P = postpartum day

### Table S17 – *EF*3 maternal homecage: Multiple comparisons by maternal timepoint

| **Contrast** | **Discovery** | **FDR Adj. P-value** |
| --- | --- | --- |
| **Pre vs. Ges** | **Yes** | **0.0205** |
| **Pre vs. P20** | **Yes** | **0.0065** |
| Ges vs. P20 | No | 0.4227 |

Pre = preconception, Ges = gestation, P = postpartum day

### Table S18 – *EF*6 Maternal Homecage: Multiple comparisons by maternal timepoint

| **Contrast** | **Discovery** | **FDR Adj. P-value** |
| --- | --- | --- |
| Pre vs. Ges | No | 0.4962 |
| **Pre vs. P20** | **Yes** | **5.10E-04** |
| **Ges vs. P20** | **Yes** | **0.0024** |

Pre = preconception, Ges = gestation, P = postpartum day

### Table S19 – *EF*1 maternal homecage observations: Multiple comparisons by postpartum day

| **Contrast** | **Discovery** | **FDR Adj. P-value** |
| --- | --- | --- |
| P1 vs. P3 | No | 0.1168 |
| **P1 vs. P8** | **Yes** | **3.07E-07** |
| **P1 vs. P14** | **Yes** | **3.07E-07** |
| **P3 vs. P8** | **Yes** | **2.04E-05** |
| **P3 vs. P14** | **Yes** | **1.44E-05** |
| P8 vs. P14 | No | 0.7596 |

P = postpartum day

### Table S20 – *EF*1 maternal homecage observations: Multiple comparisons by “off-nest” and “on-nest” behavior

| **Contrast** | **Postpartum Day** | **Experimental Condition** | **Discovery** | **FDR Adj. P-value** |
| --- | --- | --- | --- | --- |
| “off-nest” vs. “on-nest” | 1 | Control | No | 0.1161 |
| “off-nest” vs. “on-nest” | **1** | **ELS** | **Yes** | **3.74E-04** |
| “off-nest” vs. “on-nest” | 3 | Control | No | 0.1708 |
| “off-nest” vs. “on-nest” | **3** | **ELS** | **Yes** | **0.0198** |
| “off-nest” vs. “on-nest” | 8 | Control | No | 0.2861 |
| “off-nest” vs. “on-nest” | **8** | **ELS** | **Yes** | **0.0141** |
| “off-nest” vs. “on-nest” | **14** | **Control** | **Yes** | **0.0154** |
| “off-nest” vs. “on-nest” | **14** | **ELS** | **Yes** | **0.0154** |

ELS = early life stress

### Table S21 – *EF*2 maternal homecage observations: Multiple comparisons by postpartum day

| **Contrast** | **Discovery** | **FDR Adj. P-value** |
| --- | --- | --- |
| P1 vs. P3 | No | 0.9349 |
| P1 vs. P8 | No | 0.0644 |
| **P1 vs. P14** | **Yes** | **0.0015** |
| **P3 vs. P8** | **Yes** | **0.0321** |
| **P3 vs. P14** | **Yes** | **3.20E-04** |
| P8 vs. P14 | No | 0.0843 |

P = postpartum day

### Table S22 – *EF*3 maternal homecage observations: Multiple comparisons by postpartum day

| **Contrast** | **Discovery** | **FDR Adj. P-value** |
| --- | --- | --- |
| P1 vs. P3 | No | 0.4910 |
| **P1 vs. P8** | **Yes** | **0.0175** |
| **P1 vs. P14** | **Yes** | **0.0128** |
| **P3 vs. P8** | **Yes** | **0.0493** |
| **P3 vs. P14** | **Yes** | **0.0175** |
| P8 vs. P14 | No | 0.5645 |

P = postpartum day

### Table S23 – *EF*3 maternal homecage observations: Multiple comparisons by “off-nest” and “on-nest” behavior

| **Contrast** | **Postpartum Day** | **Experimental Condition** | **Discovery** | **FDR Adj. P-value** |
| --- | --- | --- | --- | --- |
| “off-nest” vs. “on-nest” | 1 | Control | No | 0.1472 |
| “off-nest” vs. “on-nest” | 1 | ELS | No | 0.1781 |
| “off-nest” vs. “on-nest” | 3 | Control | No | 0.0757 |
| “off-nest” vs. “on-nest” | 3 | ELS | No | 0.7388 |
| “off-nest” vs. “on-nest” | 8 | Control | No | 0.0757 |
| “off-nest” vs. “on-nest” | 8 | ELS | No | 0.5586 |
| “off-nest” vs. “on-nest” | **14** | **Control** | **Yes** | **0.0326** |
| “off-nest” vs. “on-nest” | 14 | ELS | No | 0.5586 |

ELS = early life stress

### Table S24 – *EF*6 maternal homecage observations: Multiple comparisons by postpartum day

| **Contrast** | **Discovery** | **FDR Adj. P-value** |
| --- | --- | --- |
| P1 vs. P3 | No | 0.8551 |
| P1 vs. P8 | No | 0.8551 |
| P1 vs. P14 | No | 0.0616 |
| P3 vs. P8 | No | 0.8551 |
| **P3 vs. P14** | **Yes** | **0.0158** |
| **P8 vs. P14** | **Yes** | **0.0158** |

P = postpartum day

### Table S25 – *EF*6 maternal homecage observations: Multiple comparisons by “off-nest” and “on-nest” behavior

| **Contrast** | **Postpartum Day** | **Experimental Condition** | **Discovery** | **FDR Adj. P-value** |
| --- | --- | --- | --- | --- |
| “off-nest” vs. “on-nest” | **1** | **Control** | **Yes** | **7.89E-04** |
| “off-nest” vs. “on-nest” | **1** | **ELS** | **Yes** | **5.77E-06** |
| “off-nest” vs. “on-nest” | **3** | **Control** | **Yes** | **4.90E-07** |
| “off-nest” vs. “on-nest” | **3** | **ELS** | **Yes** | **5.85E-07** |
| “off-nest” vs. “on-nest” | **8** | **Control** | **Yes** | **4.29E-06** |
| “off-nest” vs. “on-nest” | **8** | **ELS** | **Yes** | **5.94E-10** |
| “off-nest” vs. “on-nest” | **14** | **Control** | **Yes** | **5.77E-06** |
| “off-nest” vs. “on-nest” | **14** | **ELS** | **Yes** | **5.30E-06** |

ELS = early life stress

### Table S26 – Animal number comparisons for all correctly placed, imputed, and “cleaned and complete” (C&C) datasets

|  | **Control** | | | | | |
| --- | --- | --- | --- | --- | --- | --- |
|  | **All Correct** | **Imputed** | **C&C: *EF*1** | **C&C: *EF*2** | **C&C: *EF*3** | **C&C: *EF*6** |
| **Pre-FIT** | 10 | 20 | 20 | 18 | 20 | 16 |
| **Pre-HC** | 10 | 20 | 20 | 18 | 20 | 16 |
| **Ges** | 9 | 18 | 17 | 16 | 17 | 13 |
| **P1** | 7 | 11 | 13 | 12 | 13 | 10 |
| **P3** | 8 | 17 | 17 | 15 | 17 | 13 |
| **P8** | 8 | 15 | 15 | 15 | 15 | 11 |
| **P14** | 8 | 13 | 13 | 13 | 13 | 11 |
| **P20** | 9 | 13 | 13 | 13 | 13 | 10 |
| **Post-FIT** | 8 | 12 | 12 | 12 | 12 | 10 |
|  | **ELS** | | | | | |
|  | **All Correct** | **Imputed** | **C&C: *EF*1** | **C&C: *EF*2** | **C&C: *EF*3** | **C&C: *EF*6** |
| **Pre-FIT** | 8 | 21 | 20 | 20 | 21 | 20 |
| **Pre-HC** | 8 | 21 | 20 | 20 | 21 | 20 |
| **Ges** | 6 | 16 | 15 | 14 | 16 | 14 |
| **P1** | 4 | 11 | 9 | 9 | 10 | 10 |
| **P3** | 6 | 15 | 15 | 14 | 16 | 14 |
| **P8** | 6 | 17 | 16 | 16 | 17 | 16 |
| **P14** | 5 | 15 | 15 | 14 | 16 | 14 |
| **P20** | 5 | 15 | 14 | 14 | 15 | 13 |
| **Post-FIT** | 2 | 13 | 12 | 11 | 13 | 11 |

Pre = preconception, Ges = gestation, P = postpartum day, Post = postpartum, FIT = forced interaction test, HC = homecage, *EF* = *Electome Factor*, C&C = “Cleaned and Complete”. Note that stress *EF*s 1-6 don’t require all regions to accurately calculate *EF* scores (see Supplemental Results and Fig. S22).

### Table S27 – *EF*1 Maternal homecage: Multiple comparisons by maternal timepoint [Imputed]

| **Contrast** | **Discovery** | **FDR Adj. P-value** |
| --- | --- | --- |
| **Pre vs. Ges** | **Yes** | **4.96E-07** |
| **Pre vs. P20** | **Yes** | **3.07E-07** |
| Ges vs. P20 | No | 0.4888 |

Pre = preconception, Ges = gestation, P = postpartum day

### Table S28 – *EF*1 maternal homecage observations: Multiple comparisons by postpartum day [Imputed]

| **Contrast** | **Discovery** | **FDR Adj. P-value** |
| --- | --- | --- |
| **P1 vs. P3** | **Yes** | **5.77E-06** |
| **P1 vs. P8** | **Yes** | **5.64E-13** |
| **P1 vs. P14** | **Yes** | **2.23E-10** |
| **P3 vs. P8** | **Yes** | **3.61E-04** |
| **P3 vs. P14** | **Yes** | **0.0094** |
| P8 vs. P14 | No | 0.3575 |

P = postpartum day

### Table S29 – *EF*1 maternal homecage observations: Multiple comparisons by “off-nest” and “on-nest” behavior [Imputed]

| **Contrast** | **Postpartum Day** | **Experimental Condition** | **Discovery** | **FDR Adj. P-value** |
| --- | --- | --- | --- | --- |
| “off-nest” vs. “on-nest” | **1** | **Control** | **Yes** | **1.78E-04** |
| “off-nest” vs. “on-nest” | **1** | **ELS** | **Yes** | **1.14E-05** |
| “off-nest” vs. “on-nest” | **3** | **Control** | **Yes** | **8.94E-04** |
| “off-nest” vs. “on-nest” | **3** | **ELS** | **Yes** | **0.0032** |
| “off-nest” vs. “on-nest” | 8 | Control | No | 0.2190 |
| “off-nest” vs. “on-nest” | **8** | **ELS** | **Yes** | **8.94E-04** |
| “off-nest” vs. “on-nest” | **14** | **Control** | **Yes** | **8.76E-04** |
| “off-nest” vs. “on-nest” | **14** | **ELS** | **Yes** | **8.94E-04** |

ELS = early life stress

### Table S30 – *EF*2 maternal homecage observations: Multiple comparisons by postpartum day [Imputed]

| **Contrast** | **Discovery** | **FDR Adj. P-value** |
| --- | --- | --- |
| P1 vs. P3 | No | 0.7647 |
| **P1 vs. P8** | **Yes** | **0.0303** |
| **P1 vs. P14** | **Yes** | **9.29E-04** |
| **P3 vs. P8** | **Yes** | **0.0045** |
| **P3 vs. P14** | **Yes** | **3.28E-05** |
| P8 vs. P14 | No | 0.1043 |

P = postpartum day

### Table S31 – *EF*3 Maternal homecage: Multiple comparisons by maternal timepoint [Imputed]

| **Contrast** | **Discovery** | **FDR Adj. P-value** |
| --- | --- | --- |
| **Pre vs. Ges** | **Yes** | **0.0446** |
| Pre vs. P20 | No | 0.1487 |
| Ges vs. P20 | No | 0.5232 |

Pre = preconception, Ges = gestation, P = postpartum day

### Table S32 – *EF*3 maternal homecage observations: Multiple comparisons by postpartum day [Imputed]

| **Contrast** | **Discovery** | **FDR Adj. P-value** |
| --- | --- | --- |
| P1 vs. P3 | No | 0.8680 |
| **P1 vs. P8** | **Yes** | **0.0071** |
| **P1 vs. P14** | **Yes** | **0.0045** |
| **P3 vs. P8** | **Yes** | **0.0015** |
| **P3 vs. P14** | **Yes** | **9.62E-04** |
| P8 vs. P14 | No | 0.7841 |

P = postpartum day

### Table S33 – *EF*6 maternal homecage: Multiple comparisons by maternal timepoint [Imputed]

| **Contrast** | **Discovery** | **FDR Adj. P-value** |
| --- | --- | --- |
| Pre vs. Ges | No | 0.8941 |
| **Pre vs. P20** | **Yes** | **0.0145** |
| **Ges vs. P20** | **Yes** | **0.0145** |

Pre = preconception, Ges = gestation, P = postpartum day

### Table S34 – *EF*6 maternal homecage observations: Multiple comparisons by postpartum day [Imputed]

| **Contrast** | **Discovery** | **FDR Adj. P-value** |
| --- | --- | --- |
| P1 vs. P3 | No | 0.7502 |
| P1 vs. P8 | No | 0.7502 |
| P1 vs. P14 | No | 0.0907 |
| P3 vs. P8 | No | 0.7502 |
| **P3 vs. P14** | **Yes** | **0.0131** |
| **P8 vs. P14** | **Yes** | **0.0156** |

P = postpartum day

### Table S35 – *EF*6 maternal homecage observations: Multiple comparisons by “off-nest” and “on-nest” behavior [Imputed]

| **Contrast** | **Postpartum Day** | **Experimental Condition** | **Discovery** | **FDR Adj. P-value** |
| --- | --- | --- | --- | --- |
| “off-nest” vs. “on-nest” | **1** | **Control** | **Yes** | **1.31E-04** |
| “off-nest” vs. “on-nest” | **1** | **ELS** | **Yes** | **7.94E-07** |
| “off-nest” vs. “on-nest” | **3** | **Control** | **Yes** | **5.94E-10** |
| “off-nest” vs. “on-nest” | **3** | **ELS** | **Yes** | **1.54E-06** |
| “off-nest” vs. “on-nest” | **8** | **Control** | **Yes** | **5.41E-07** |
| “off-nest” vs. “on-nest” | **8** | **ELS** | **Yes** | **1.48E-11** |
| “off-nest” vs. “on-nest” | **14** | **Control** | **Yes** | **1.73E-06** |
| “off-nest” vs. “on-nest” | **14** | **ELS** | **Yes** | **2.90E-07** |

ELS = early life stress

### Table S36 – Nest visits per hour: Multiple comparisons by maternal timepoint

| **Contrast** | **Discovery** | **FDR Adj. P-value** |
| --- | --- | --- |
| P1 vs. P3 | No | 0.0623 |
| **P1 vs. P8** | **Yes** | **7.27E-06** |
| **P1 vs. P14** | **Yes** | **2.83E-10** |
| **P3 vs. P8** | **Yes** | **0.0014** |
| **P3 vs. P14** | **Yes** | **3.58E-08** |
| **P8 vs. P14** | **Yes** | **0.0044** |

P = postpartum day

### Table S37 – Duration of care visits: Multiple comparisons by maternal timepoint

| **Contrast** | **Discovery** | **FDR Adj. P-value** |
| --- | --- | --- |
| P1 vs. P3 | No | 0.8924 |
| P1 vs. P8 | No | 0.9680 |
| **P1 vs. P14** | **Yes** | **0.0013** |
| P3 vs. P8 | No | 0.8924 |
| **P3 vs. P14** | **Yes** | **0.0013** |
| **P8 vs. P14** | **Yes** | **0.0013** |

P = postpartum day

### Table S38 – *EF*1 and P4 Pup Retrieval: Multiple comparisons by task stage

| **Contrast** | **Discovery** | **FDR Adj. P-value** |
| --- | --- | --- |
| **Baseline vs. Trials** | **Yes** | **<0.0001** |
| **Baseline vs. Retrieval** | **Yes** | **<0.0001** |
| **Trials vs. Retrieval** | **Yes** | **0.0222** |

### Table S39 – *EF*6 and P4 Pup Retrieval: Multiple comparisons by task stage

| **Contrast** | **Discovery** | **FDR Adj. P-value** |
| --- | --- | --- |
| **Baseline vs. Trials** | **Yes** | **0.0089** |
| **Baseline vs. Retrieval** | **Yes** | **0.0043** |
| **Trials vs. Retrieval** | **Yes** | **0.0424** |

### Table S40 – *EF*1 FIT: Multiple comparisons by FIT stage [Cleaned and Complete]

| **Contrast** | **Discovery** | **FDR Adj. P-value** |
| --- | --- | --- |
| **Homecage vs. Empty** | **Yes** | **3.19E-09** |
| **Homecage vs. Aggressor** | **Yes** | **3.14E-13** |
| Empty vs. Aggressor | No | 0.0680 |

Pre = preconception, Post = postpartum

### Table S41 – *EF*1 FIT: Multiple comparisons by maternal timepoint [Cleaned and Complete]

| **Contrast** | **Task Stage** | **Experimental Condition** | **Discovery** | **FDR Adj. P-value** |
| --- | --- | --- | --- | --- |
| Pre vs. Post | **Homecage** | **Control** | **Yes** | **0.0392** |
| Pre vs. Post | **Homecage** | **ELS** | **Yes** | **0.0438** |
| Pre vs. Post | Empty | Control | No | 0.2471 |
| Pre vs. Post | **Empty** | **ELS** | **Yes** | **0.0392** |
| Pre vs. Post | Aggressor | Control | No | 0.2241 |
| Pre vs. Post | Aggressor | ELS | No | 0.0502 |

Pre = preconception, Post = postpartum, ELS = early life stress

### Table S42 – *EF*2 FIT: Multiple comparisons by FIT stage [Cleaned and Complete]

| **Contrast** | **Discovery** | **FDR Adj. P-value** |
| --- | --- | --- |
| **Homecage vs. Empty** | **Yes** | **0.0053** |
| **Homecage vs. Aggressor** | **Yes** | **1.61E-05** |
| **Empty vs. Aggressor** | **Yes** | **3.72E-11** |

### Table S43 – *EF*2 FIT: Multiple comparisons by maternal timepoint [Cleaned and Complete]

| **Contrast** | **Task Stage** | **Experimental Condition** | **Discovery** | **FDR Adj. P-value** |
| --- | --- | --- | --- | --- |
| Pre vs. Post | Homecage | Control | No | 0.0514 |
| Pre vs. Post | Homecage | ELS | No | 0.0809 |
| Pre vs. Post | **Empty** | **Control** | **Yes** | **0.0284** |
| Pre vs. Post | Empty | ELS | No | 0.0809 |
| Pre vs. Post | Aggressor | Control | No | 0.1148 |
| Pre vs. Post | **Aggressor** | **ELS** | **Yes** | **0.0284** |

Pre = preconception, Post = postpartum, ELS = early life stress

### Table S44 – *EF*1 FIT: Multiple comparisons by FIT stage [Imputed]

| **Contrast** | **Discovery** | **FDR Adj. P-value** |
| --- | --- | --- |
| Homecage vs. Empty | **Yes** | **2.57E-14** |
| Homecage vs. Aggressor | **Yes** | **7.75E-18** |
| Empty vs. Aggressor | No | 0.1386 |

### Table S45 – *EF*1 FIT: Multiple comparisons by maternal timepoint [Imputed]

| **Contrast** | **Task Stage** | **Experimental Condition** | **Discovery** | **FDR Adj. P-value** |
| --- | --- | --- | --- | --- |
| Pre vs. Post | Homecage | Control | No | 0.2344 |
| Pre vs. Post | **Homecage** | **ELS** | **Yes** | **0.0016** |
| Pre vs. Post | Empty | Control | No | 0.5944 |
| Pre vs. Post | Empty | ELS | No | 0.0690 |
| Pre vs. Post | Aggressor | Control | No | 0.0690 |
| Pre vs. Post | Aggressor | ELS | No | 0.2204 |

Pre = preconception, Post = postpartum, ELS = early life stress

### Table S46 – *EF*2 FIT: Multiple comparisons by FIT stage [Imputed]

| **Contrast** | **Discovery** | **FDR Adj. P-value** |
| --- | --- | --- |
| **Homecage vs. Empty** | **Yes** | **1.47E-04** |
| **Homecage vs. Aggressor** | **Yes** | **4.58E-05** |
| **Empty vs. Aggressor** | **Yes** | **3.28E-13** |

### Table S47 – *EF*2 FIT: Multiple comparisons by maternal timepoint [Imputed]

| **Contrast** | **Task Stage** | **Experimental Condition** | **Discovery** | **FDR Adj. P-value** |
| --- | --- | --- | --- | --- |
| Pre vs. Post | Homecage | Control | No | 0.8357 |
| Pre vs. Post | Homecage | ELS | No | 0.8357 |
| Pre vs. Post | **Empty** | **Control** | **Yes** | **0.0188** |
| Pre vs. Post | **Empty** | **ELS** | **Yes** | **0.0425** |
| Pre vs. Post | Aggressor | Control | No | 0.8357 |
| Pre vs. Post | **Aggressor** | **ELS** | **Yes** | **0.0295** |

Pre = preconception, Post = postpartum, ELS = early life stress

### Table S48 – Supervision weights used in maternal *EF* training

| ***EF* Model** | **Mu (µ)** |
| --- | --- |
| *EF_on-nest_* | 0.05 |
| *EF_on-nest-1-Hz_* | 0.045 |
| *EF_licking_* | 0.07 |
| *EF_lick-groom_* | 0.50 |
| *EF_stage_* | 0.025 |
| *EF_stage-1-Hz_* | 0.03 |

1	Talbot, A., Dunson, D., Dzirasa, K. & Carlson, D. Estimating a brain network predictive of stress and genotype with supervised autoencoders. *J R Stat Soc Ser C Appl Stat* **72**, 912–936 (2023). <https://doi.org/10.1093/jrsssc/qlad035>

2	Hultman, R. *et al.* Brain-wide Electrical Spatiotemporal Dynamics Encode Depression Vulnerability. *Cell* **173**, 166–180 e114 (2018). <https://doi.org/10.1016/j.cell.2018.02.012>

3	Hultman, I. *et al*. Sparse Separable Factor Analysis in the Complex Domain with an Application to Local Field Potential Data. *arXiv* (2026). <https://doi.org/10.48550/arXiv.2608.21551>

4	Gerasimenko, M. *et al.* Receptor for advanced glycation end-products (RAGE) plays a critical role in retrieval behavior of mother mice at early postpartum. *Physiol Behav* **235**, 113395 (2021). <https://doi.org/10.1016/j.physbeh.2021.113395>

5	Firestein, M. R. *et al.* - Rates of Positive M-CHAT-R Screenings by Pandemic Birth and Prenatal SARS-CoV-2 Exposure. (2024).

6	Schaffler, M. D. J., Micah; Hing, Ben; Kahler, Paul; Hultman, Ian; Srivastava, Sanvesh; Arnold, Justin; Blendy, Julie N; Hultman, Rainbo; Abdus-Saboor, Ishmail A critical role for touch neurons in a skin-brain pathway for stress resilience. (2022). <https://doi.org/doi>: <https://doi.org/10.1101/2022.05.23.493062>

7	Barba-Muller, E., Craddock, S., Carmona, S. & Hoekzema, E. Brain plasticity in pregnancy and the postpartum period: links to maternal caregiving and mental health. *Arch Womens Ment Health* **22**, 289–299 (2019). <https://doi.org/10.1007/s00737-018-0889-z>

8	Kikusui, T. & Mori, Y. Behavioural and neurochemical consequences of early weaning in rodents. *J Neuroendocrinol* **21**, 427–431 (2009). <https://doi.org/10.1111/j.1365-2826.2009.01837.x>

9	Kaiser, R. H. *et al.* Childhood stress, grown-up brain networks: corticolimbic correlates of threat-related early life stress and adult stress response. *Psychol Med* **48**, 1157–1166 (2018). <https://doi.org/10.1017/S0033291717002628>

10	Rombaut, C., Roura-Martinez, D., Lepolard, C. & Gascon, E. Brief and long maternal separation in C57Bl6J mice: behavioral consequences for the dam and the offspring. *Front Behav Neurosci* **17**, 1269866 (2023). <https://doi.org/10.3389/fnbeh.2023.1269866>

11	Murgatroyd, C. A., Pena, C. J., Podda, G., Nestler, E. J. & Nephew, B. C. Early life social stress induced changes in depression and anxiety associated neural pathways which are correlated with impaired maternal care. *Neuropeptides* **52**, 103–111 (2015). <https://doi.org/10.1016/j.npep.2015.05.002>
